# Auditory Gamma-Frequency Entrainment Abolishes Working Memory Deficits in a Rodent Model of Autism

**DOI:** 10.1101/2025.06.12.659284

**Authors:** Jorge Cardoso, Marta Luis, Catarina Mesquita, Luísa V. Lopes, Miguel Remondes

**Affiliations:** Gulbenkian Institute for Molecular Medicine Lisbon, 1649-028, Portugal; Faculdade de Medicina Universidade de Lisboa Lisbon, 1649-028, Portugal; Faculdade de Medicina Veterinária Universidade Lusófona, Lisbon, 1749-024, Portugal

## Abstract

Individuals on the autism spectrum often exhibit atypical responses to sensory stimuli and difficulties with behavioral regulation, reflecting altered functional patterns in core sensory-motor circuits. While impairments in higher cognitive functions are a hallmark of autism, they remain underexplored in rodent models compared to basic sensory-motor deficits. Furthermore, these sensory processing differences suggest the potential for using patterned auditory stimulation to modulate neural activity and mitigate symptoms. In this study, we investigated higher cognitive function in the valproic acid (VPA) rodent model of autism and evaluated whether auditory entrainment could ameliorate associated impairments. Pregnant dams received an intraperitoneal injection of VPA on embryonic day 12.5 (E12.5), and offspring were subsequently tested on a delayed non-match to place (DNMP) task to assess working memory. VPA-exposed animals showed significantly impaired performance and altered trial-by-trial learning dynamics, indicating deficits in working memory-based decision making. Notably, gamma-frequency auditory stimulation delivered during DNMP sessions led to increased gamma-frequency oscillations, and enhanced power in other bands (theta, beta) across brain regions, as assessed by LFP recordings in freely moving animals. More importantly, such stimulation eliminated the observed working memory impairments, enhancing performance both during and after entrainment. This provides strong evidence for the therapeutic potential of sensory entrainment in reversing autism-related cognitive deficits by modulating dysfunctional neural circuits.

## Introduction

A critical component of adaptive behavior is the ability to make decisions based on the spatial structure of one’s environment - a capacity broadly referred to as spatial cognition or spatial goal-directed behavior (GDB). This cognitive function begins with the selection of relevant sensory inputs from a continuous stream of stimuli and culminates in the storage of spatial representations within cortico-hippocampal circuits, forming cognitive maps characterized by diverse neural activity patterns sensitive to both egocentric (self-referenced) and allocentric (world-referenced) spatial variables ^1–3^.

Emerging evidence suggests that the neural circuits connecting early sensory processing, memory formation, and anterior executive regions involved in behavioral control are altered in autism spectrum disorder (ASD). These alterations may contribute to the hallmark inflexibility observed in ASD—manifesting as rigid, self-centered rather than adaptive, context-sensitive behaviors, thoughts, and emotions ^4^. While a wide range of spontaneous and stimulus-driven behaviors have been documented in animals prenatally exposed to valproic acid (VPA)—a well-established rodent model of ASD—higher cognitive functions, such as spatial GDB, remain relatively unexplored in this model. If VPA exposure indeed mirrors core ASD phenotypes, deficits in early cognitive functions, including the ability to utilize allocentric spatial information to guide behavior, would be expected. Recent studies in neurodegenerative models have shown that patterned sensory stimulations - specifically sensory entrainment - can mitigate cognitive deficits ^5^. Given the reliance of higher cognitive processes on effective sensory integration, we hypothesize that similar strategies may also alleviate cognitive impairments associated with ASD. In particular, we propose that auditory gamma-frequency entrainment could enhance spatial working memory and goal-directed decision making in VPA-exposed rodents.

To test these hypotheses, we employed a Delayed Non-Match to Place (DNMP) task designed to robustly engage working memory (WM) over a 20-second delay. In subsequent sessions, animals performed this task while exposed to auditory gamma-frequency (40 Hz) entrainment. The DNMP paradigm requires animals to recall a prior spatial decision, and the time it was made, in order to disambiguate it from similar, previously rewarded decisions ^6–11^. In our established version of this task ^12,13^, animals navigate one arm of a T-maze during a “Sample” run, followed by a 20-second delay period. In the ensuing “Test” run, both arms are made available, and a reward is given for choosing the arm opposite to the one previously sampled. We applied this paradigm to animals prenatally exposed to either saline or valproic acid (VPA; 400 or 600 mg/kg body weight, intraperitoneally administered to pregnant dams), modeling varying degrees of ASD-like traits. Importantly, animals were assigned to groups without prior behavioral screening, ensuring an unbiased approach to testing VPA’s impact on memory-guided decision making.

To evaluate behavioral performance, we measured both trial-level accuracy (correct/error) and session-level success (% correct trials), analyzing outcomes across individuals, sessions, and groups (CTRL, VPA400, VPA600). To examine the impact of VPA exposure on task performance, we used a multivariable Generalized Linear Mixed Model (GLMM), accounting for both categorical and continuous predictors. Additionally, we conducted trial-by-trial analyses using logistic regression and state-space modeling to assess learning dynamics and estimate the probability of correct choices over time.

Given that perseverative behavior—a hallmark of ASD—is often maladaptive in tasks requiring cognitive flexibility, we also quantified individual perseveration biases. A bias index was computed for each animal by calculating the difference in left versus right arm choices during Test trials, normalized by the total number of trials.

To determine whether auditory entrainment could modulate the underlying neural dynamics associated with these behaviors, we recorded multi-unit neural activity using a multi-tetrode drive implanted in brain areas implicated in ASD pathogenesis. In a dedicated experimental group (VPA600), animals underwent three consecutive blocks of six DNMP sessions, with gamma-frequency auditory stimulation applied during the middle block. Behavioral changes across these phases were assessed using the performance metrics described above, enabling us to evaluate both behavioral and neural effects of sensory entrainment.

## Results

### Group-level performance reveals working memory impairment following prenatal VPA exposure

In the DNMP task, animals first complete a “Sample” trial, during which one arm of the T-maze is blocked. In the subsequent “Test” trial, both arms are accessible, and rats are rewarded for choosing the arm opposite to the one previously visited during the Sample trial (Figure 1A).

**Figure 1.**
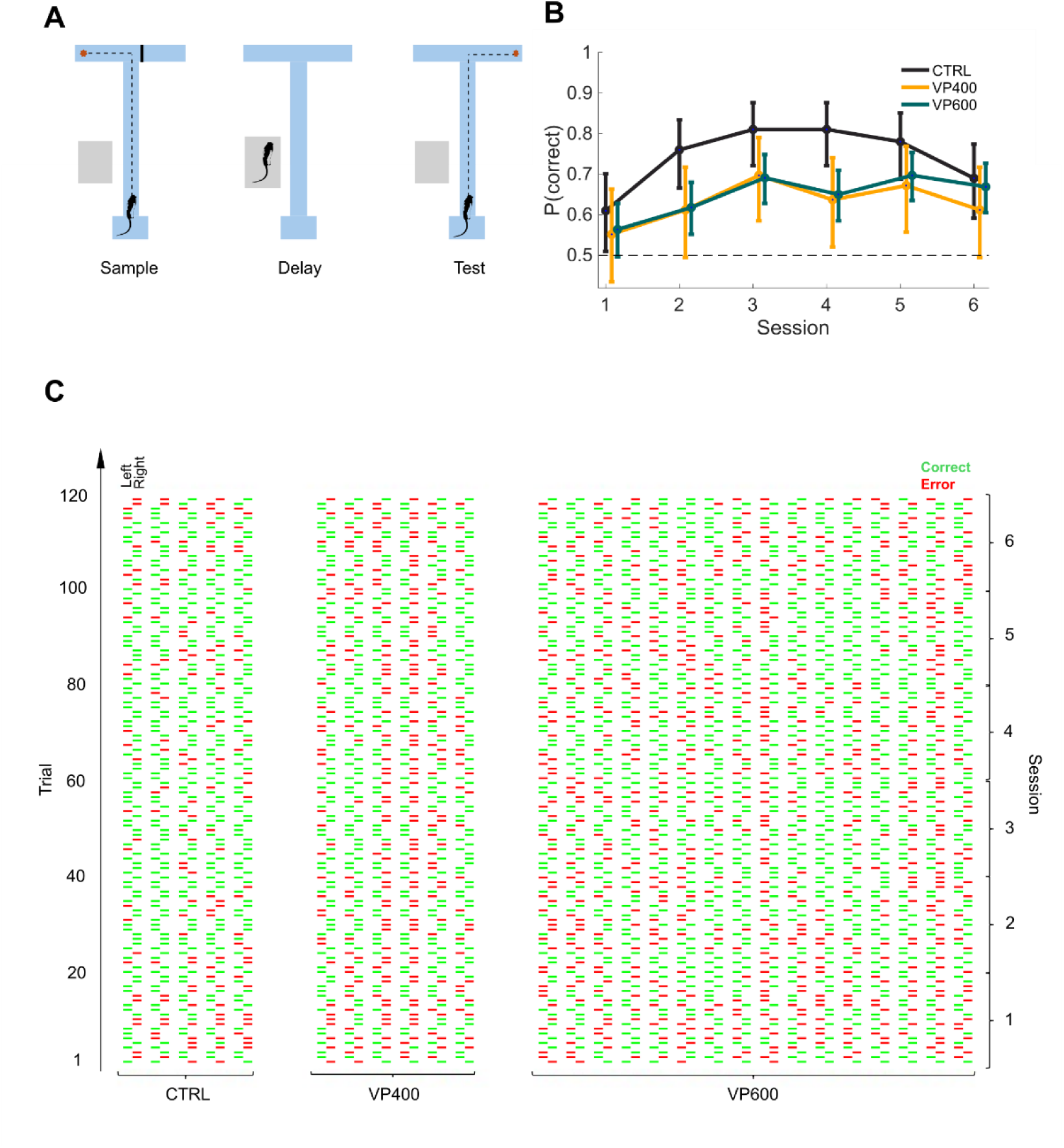
Delayed non-match to place (DNMP) performance in CTRL and VPA-exposed animals. **(A)** Experimental design showing DNMP task protocol across six sessions (S1–S6). **(B)** Average performance (probability of correct trials) across sessions for CTRL, VP400, and VP600 groups. **(C)** Raw behavioral data from individual animals across all sessions. Error trials are marked red and correct trials green. Tick orientation corresponds to the maze arm choice (left vs right).

A group-level analysis of performance across all animals (Figure 1B) revealed a general improvement over sessions 1 to 6 (χ² = 30.39, *p* < 0.001; Fixed Effect Omnibus test). However, animals exposed in utero to valproic acid (both VP600 and VP400) displayed significantly impaired performance compared to control animals (CTRL), with a robust effect of treatment (Figure 1B–C; Table 2; GLMM: χ² = 15.05, *p* < 0.001; Fixed Effect Omnibus test). No significant performance difference was observed between the VP600 and VP400 groups (*p* < 0.001, Bonferroni post hoc test). Interestingly, this performance gap disappeared by session 6, with all groups achieving accuracy well above chance (50%). Notably, the CTRL group showed a slight decline in performance by session 6.

These findings suggest that prenatal VPA exposure impairs spatial working memory (SWM) in the DNMP task. To better understand the underlying learning dynamics, we next analyzed performance progression within sessions (S1–S6) and across individual trials.

### VPA-exposed animals exhibit disrupted trial-by-trial learning dynamics and reduced within-session performance stability

To assess within-session behavioral dynamics in individual animals, we analyzed how trial-by-trial performance evolved over time. Specifically, we aimed to estimate the probability of a correct choice on each trial based on preceding trials. For this, we employed two complementary approaches: Binary Logistic Regression (BLR) and State-Space Modeling (SSM), allowing us to track trial-wise learning and performance maintenance across sessions S1–S6 (see Supplementary Figure 1 for raw data, BLR, and SSM outputs for each animal across groups).

In CTRL animals, we observed not only a clear improvement across sessions, as shown earlier, but also a consistent within-session increase in the probability of correct responses. Both BLR and SSM analyses converged on these results, with later sessions showing high and stable performance. Group-level analyses further confirmed this pattern: in 3.2 out of 6 sessions, the 95% confidence intervals (CIs) of the mean trial-wise correct probability exceeded the 50% chance level, indicating reliable and above-chance performance (Figure 2A). On average, SSM yielded stable trial-by-trial estimates in 4.8 of 6 sessions per animal (Figure S1; Table 1).

**Figure 2.**
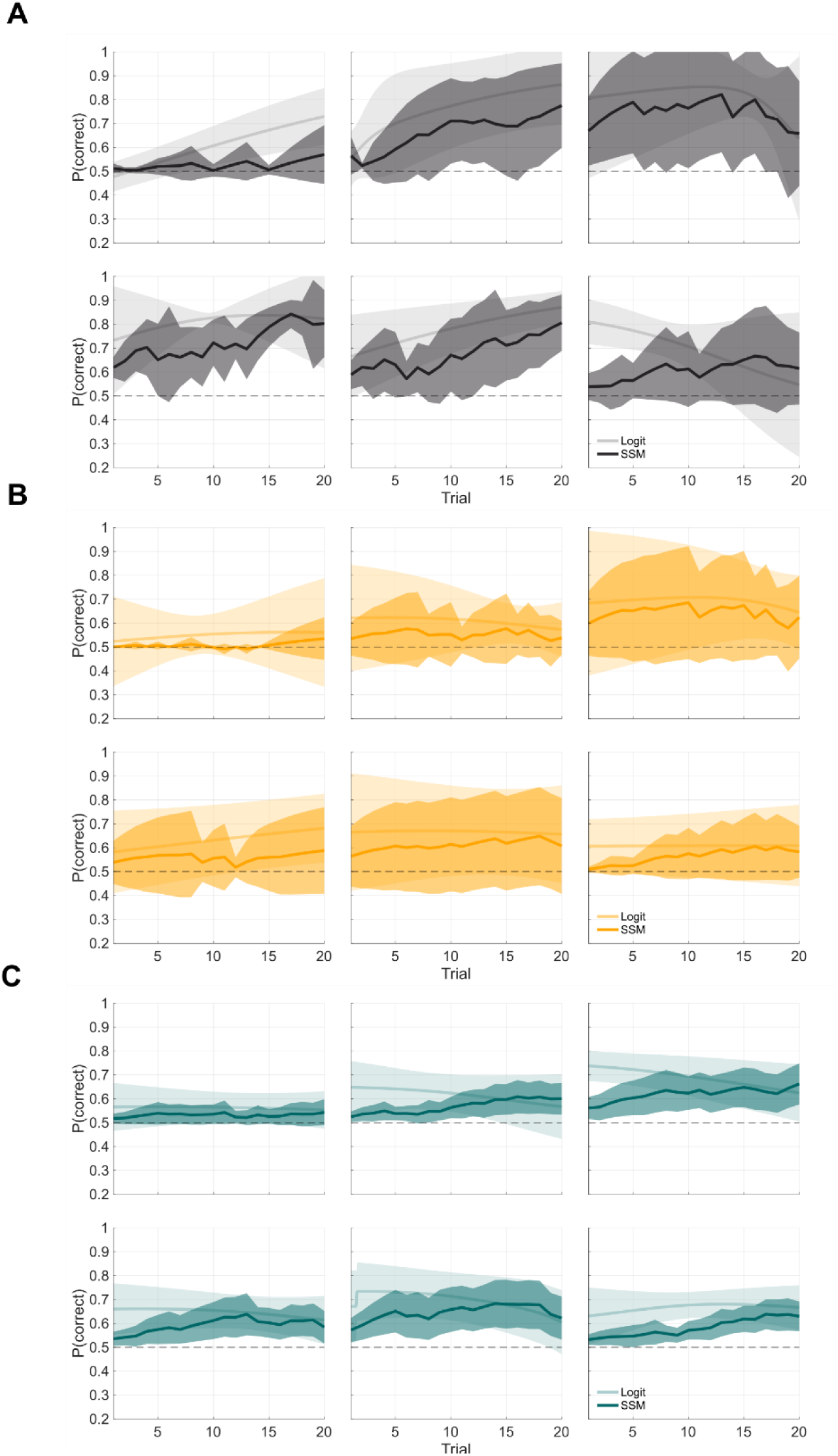
Trial-by-trial dynamics of performance in CTRL and VPA animals. State-space Modeling (darker shade) overlaid on Binary Logistic Regression (BLR) (light shade) trial-by-trial probability estimates, +/-95% CI showing average group-level performance over time, on each of the 6 DNMP sessions (individual plots), for CTRL (grey in panel A), VP400 (yellow in panel B) and VP600 (green in panel C) animals.

**Table 1.**
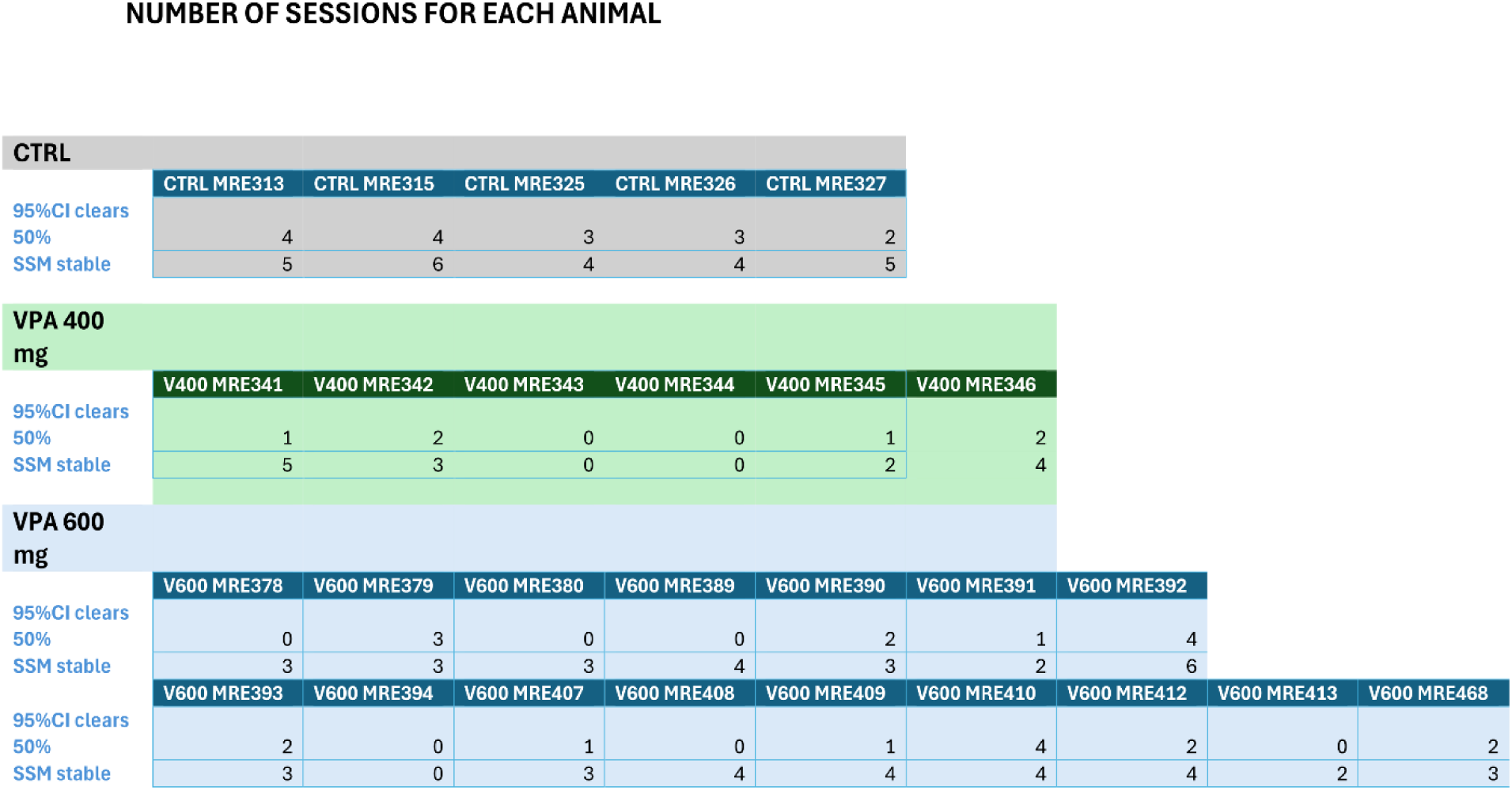
Trial-by-trial Logit and SSM estimateś robustness across individual animals and sessions.

**Table 2.**
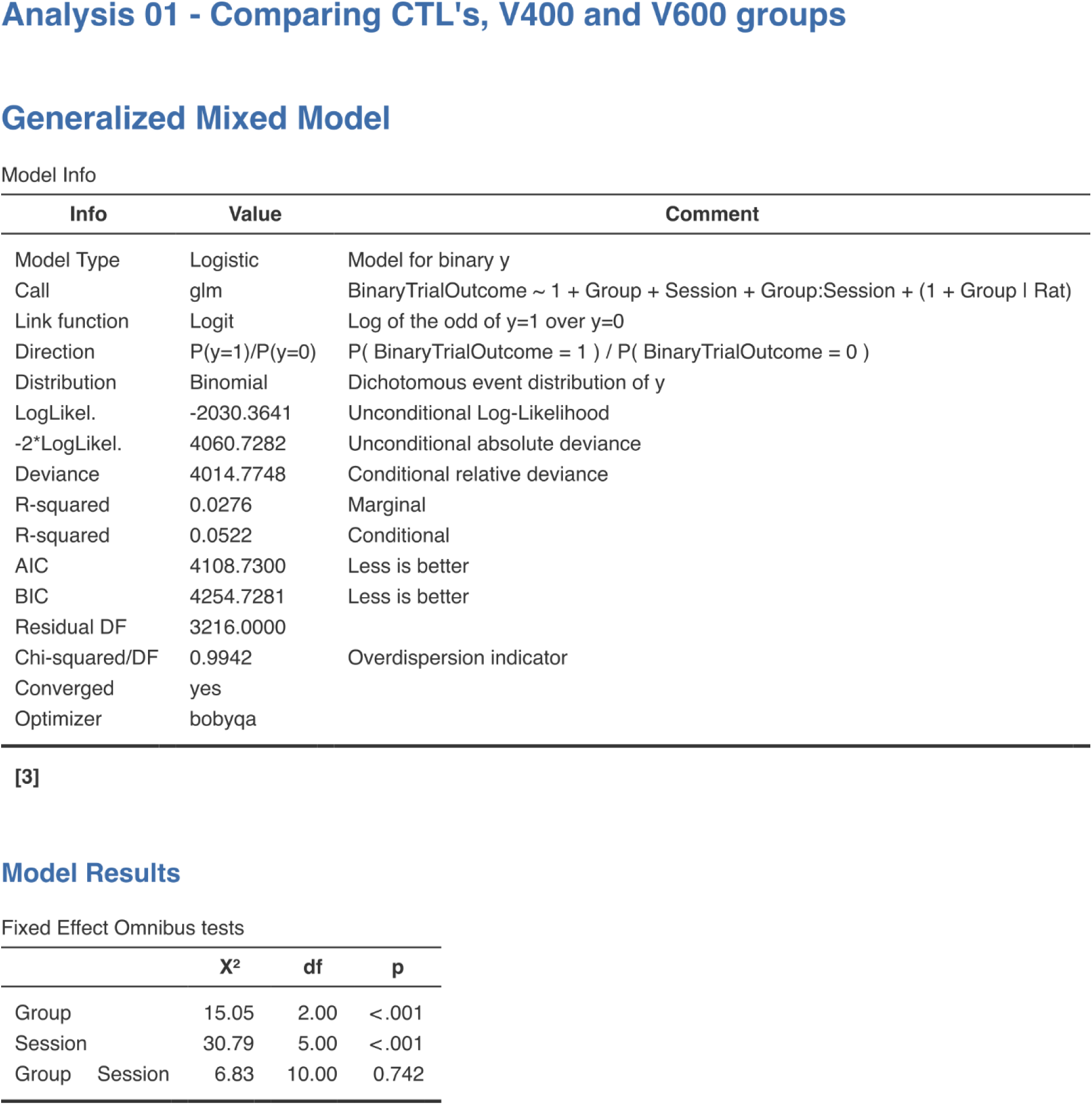
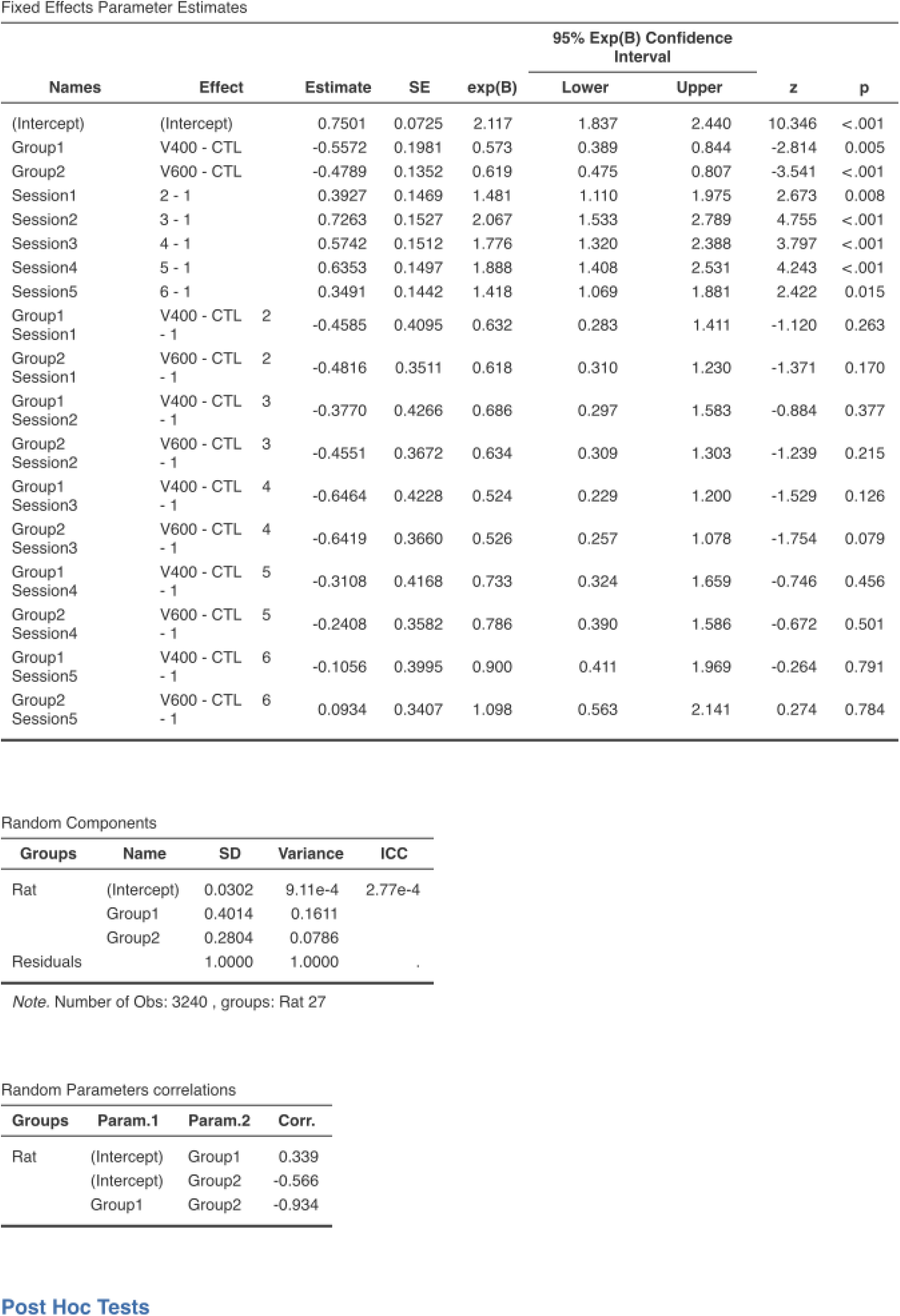
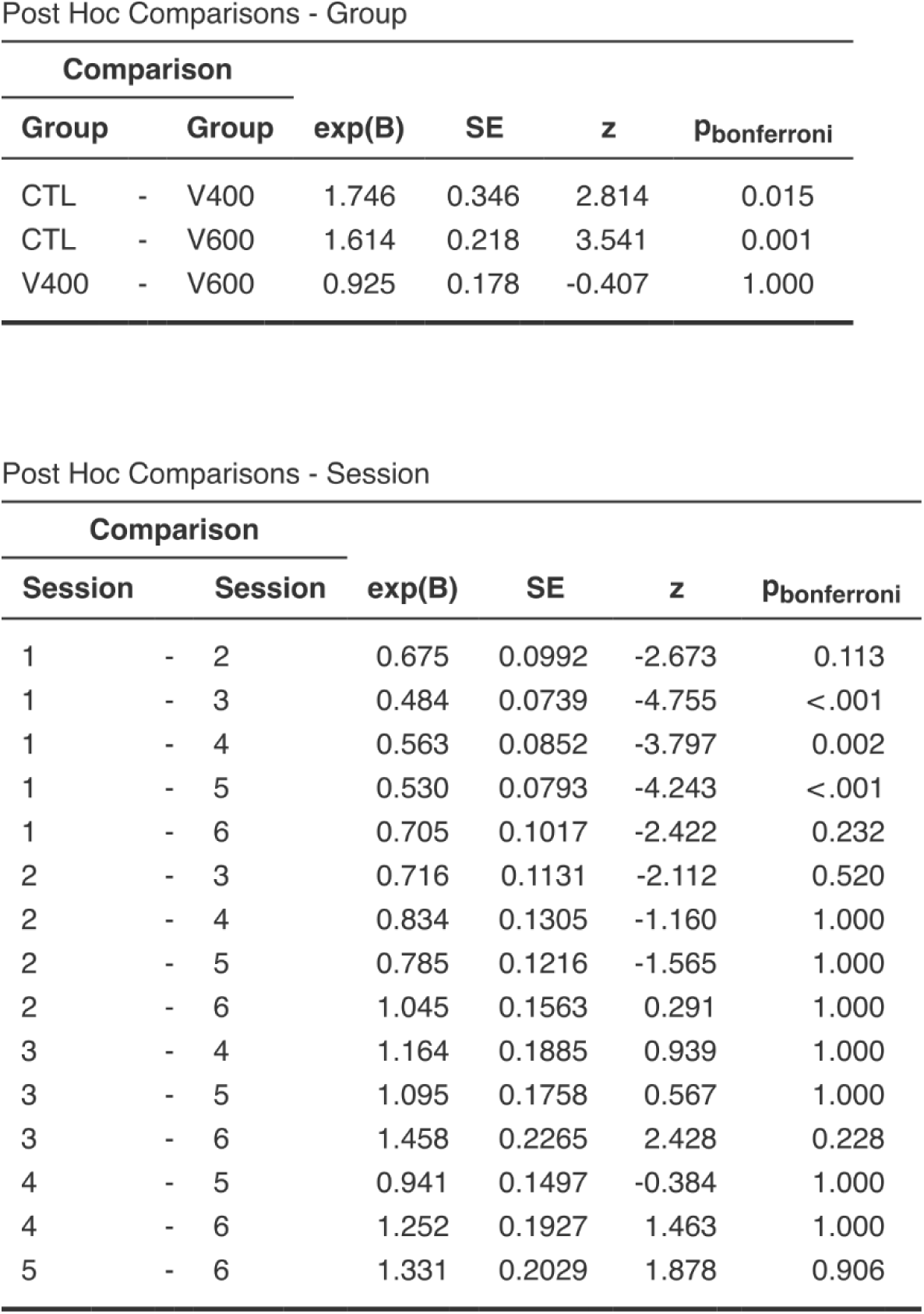
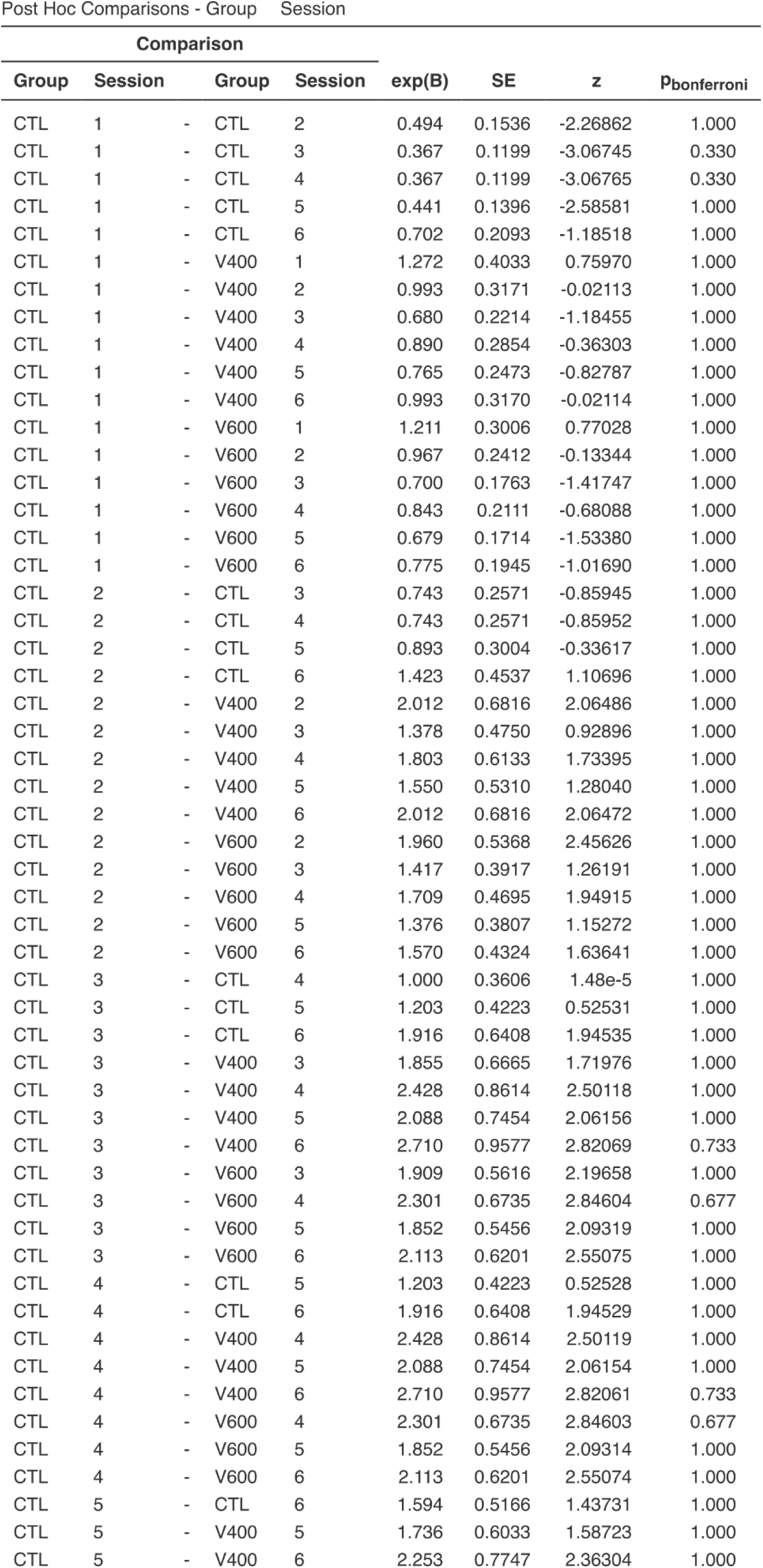
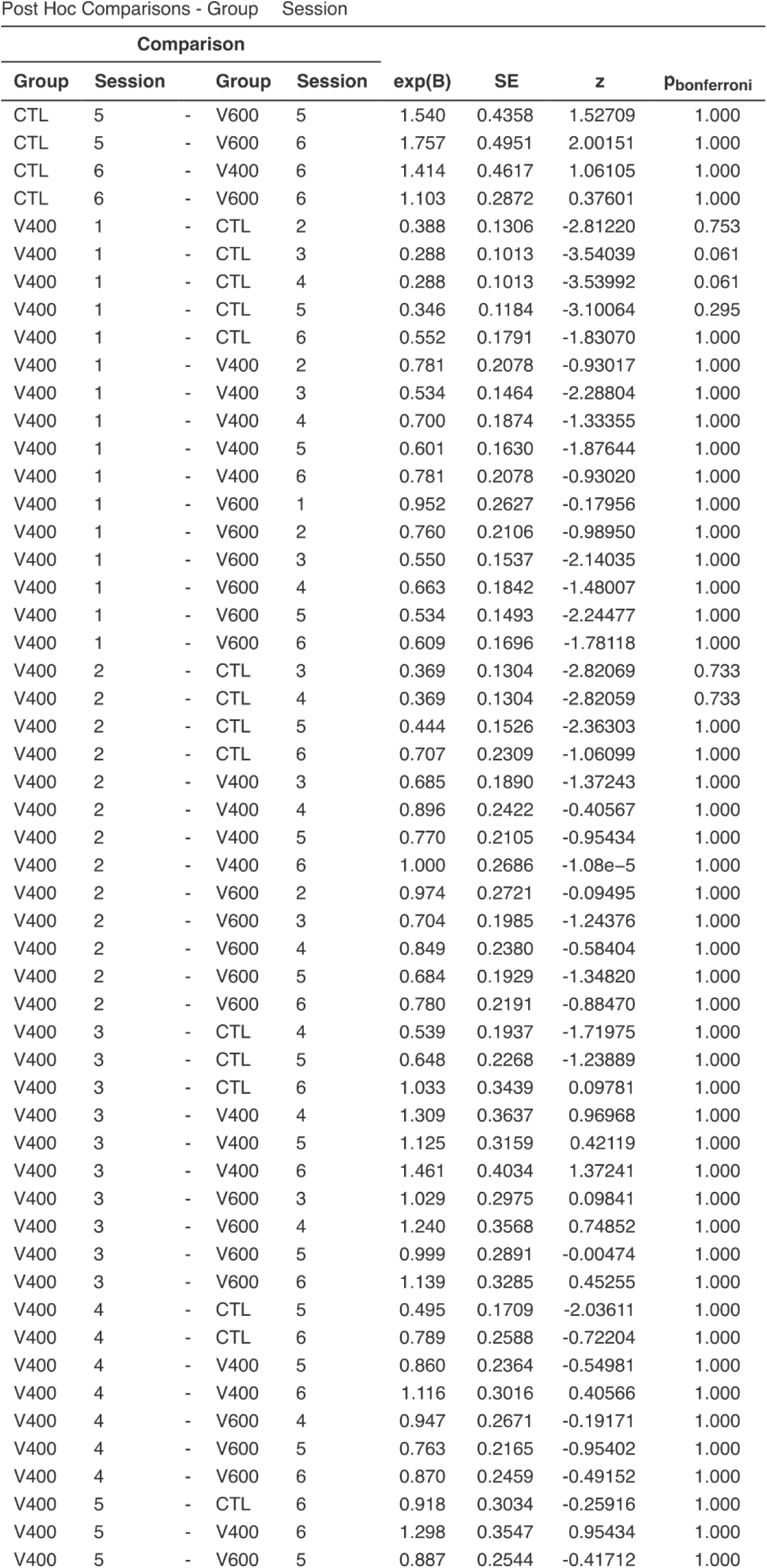
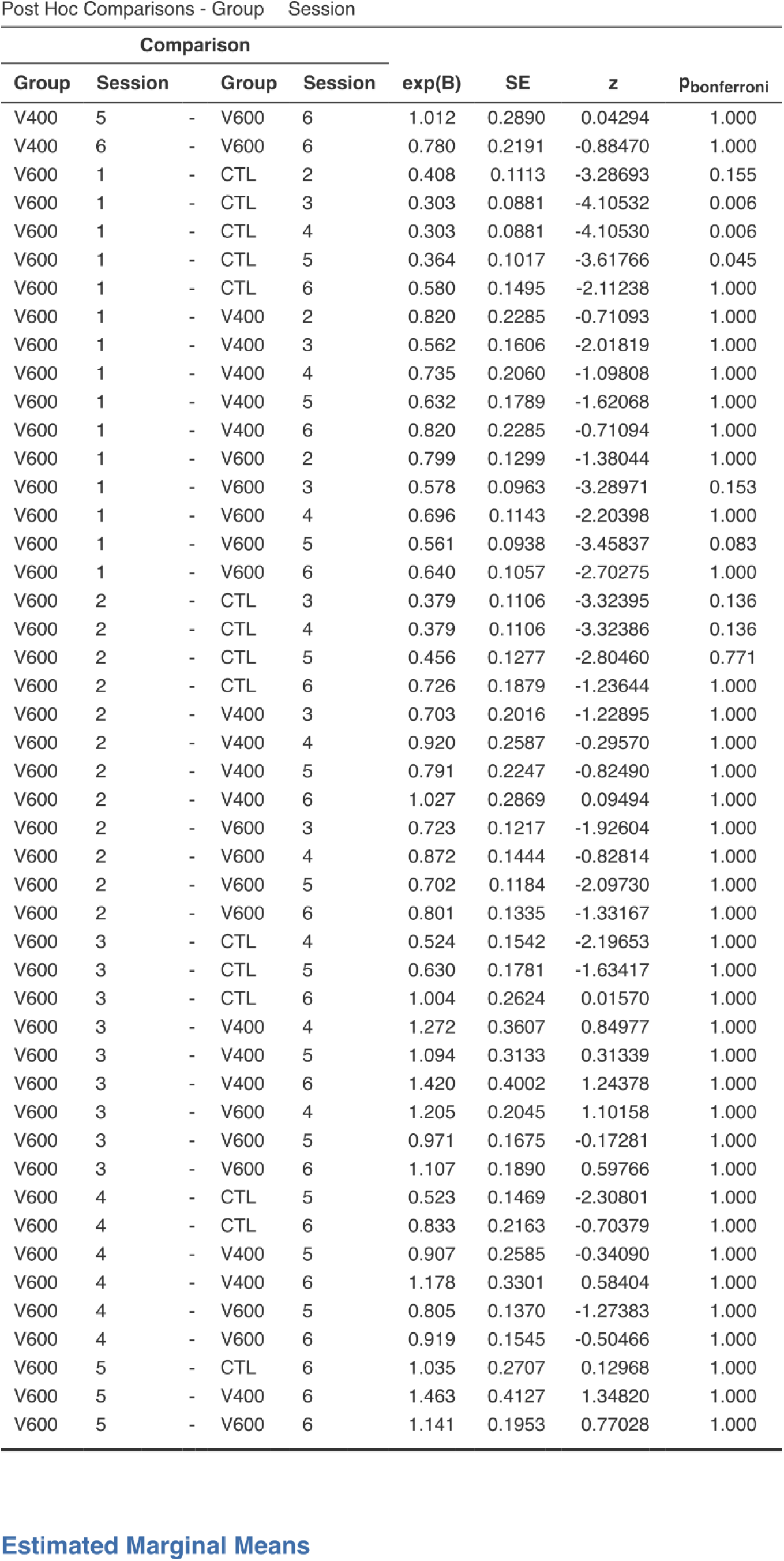
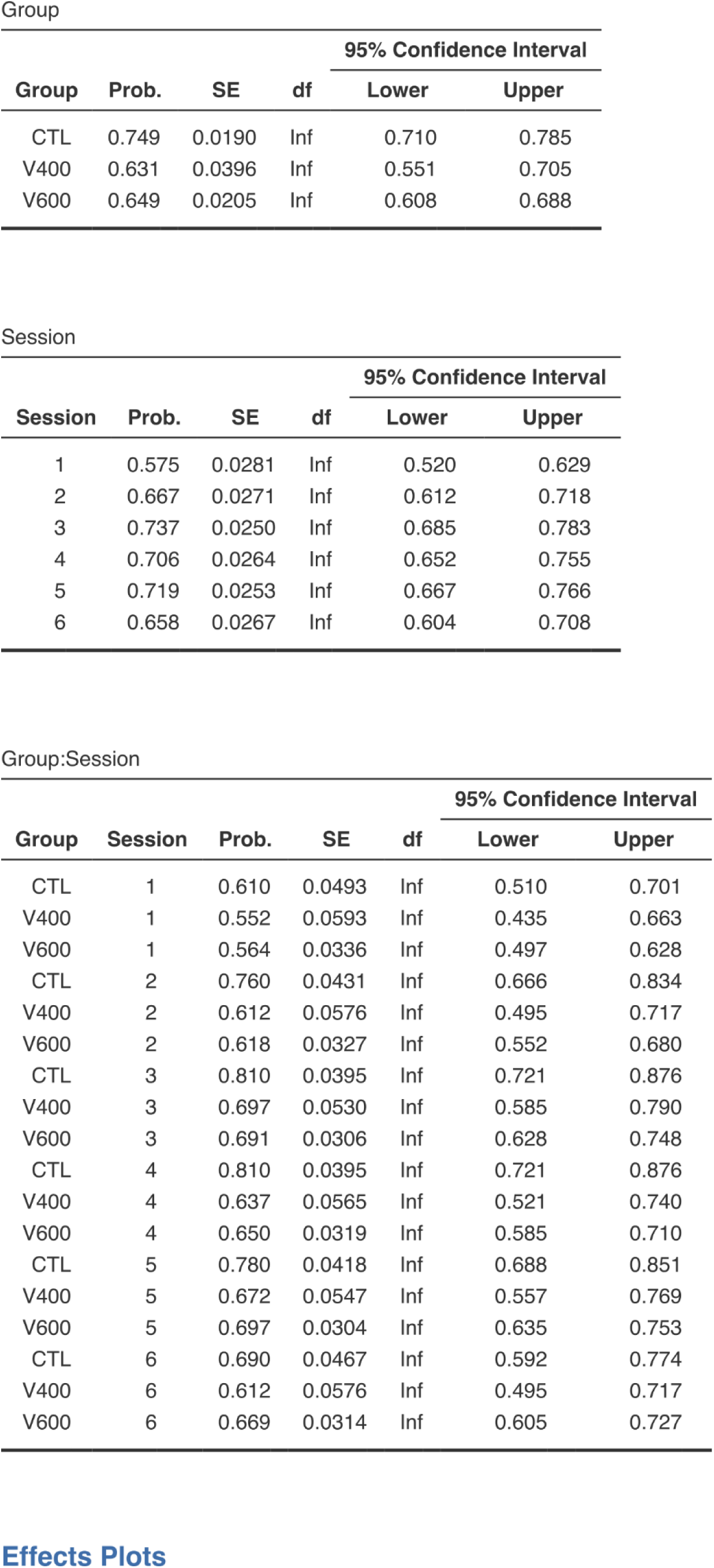
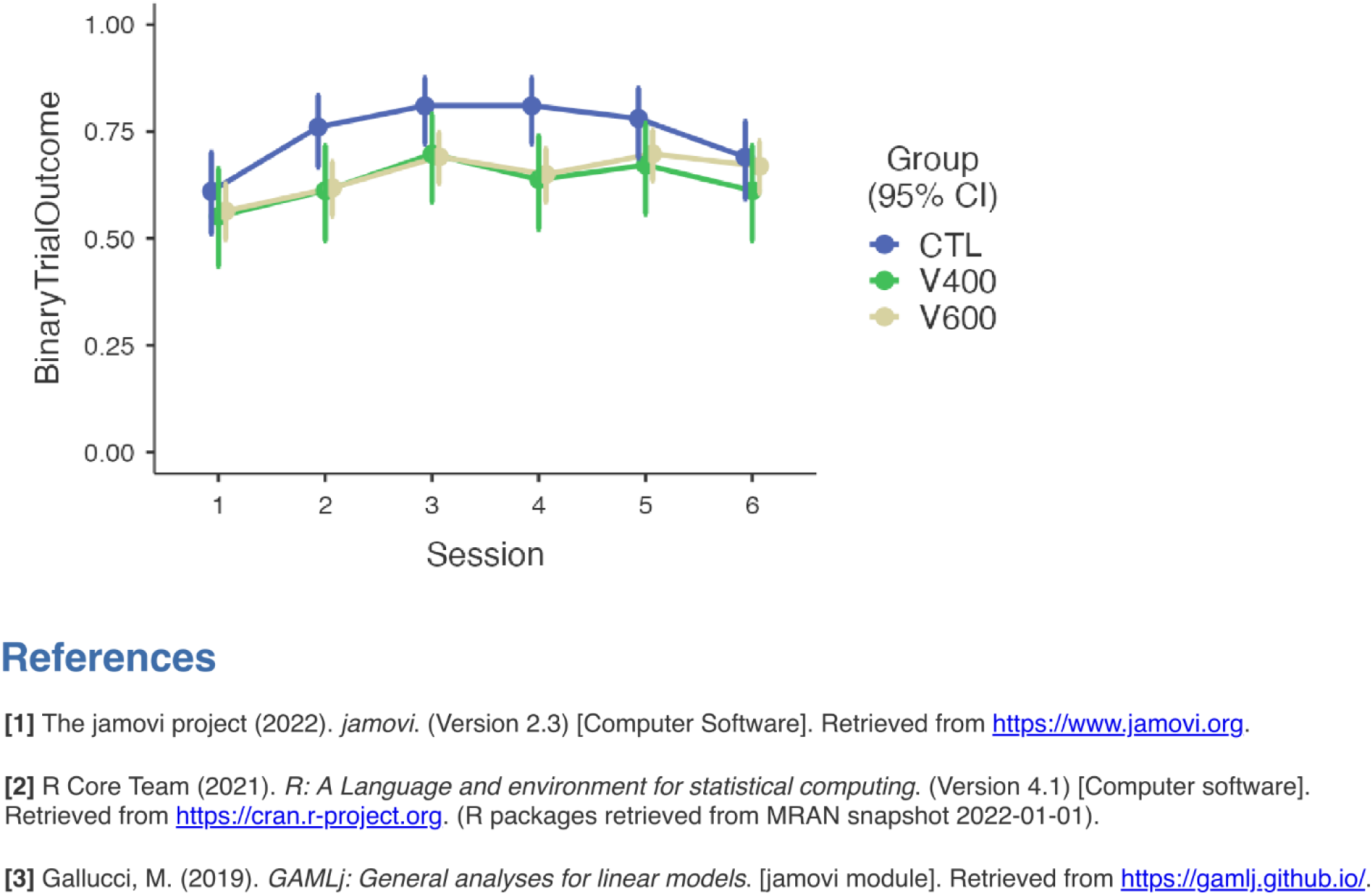
GLMM table for the comparison between CTLR, VP400 and VP600 animals.

**Table 3.**
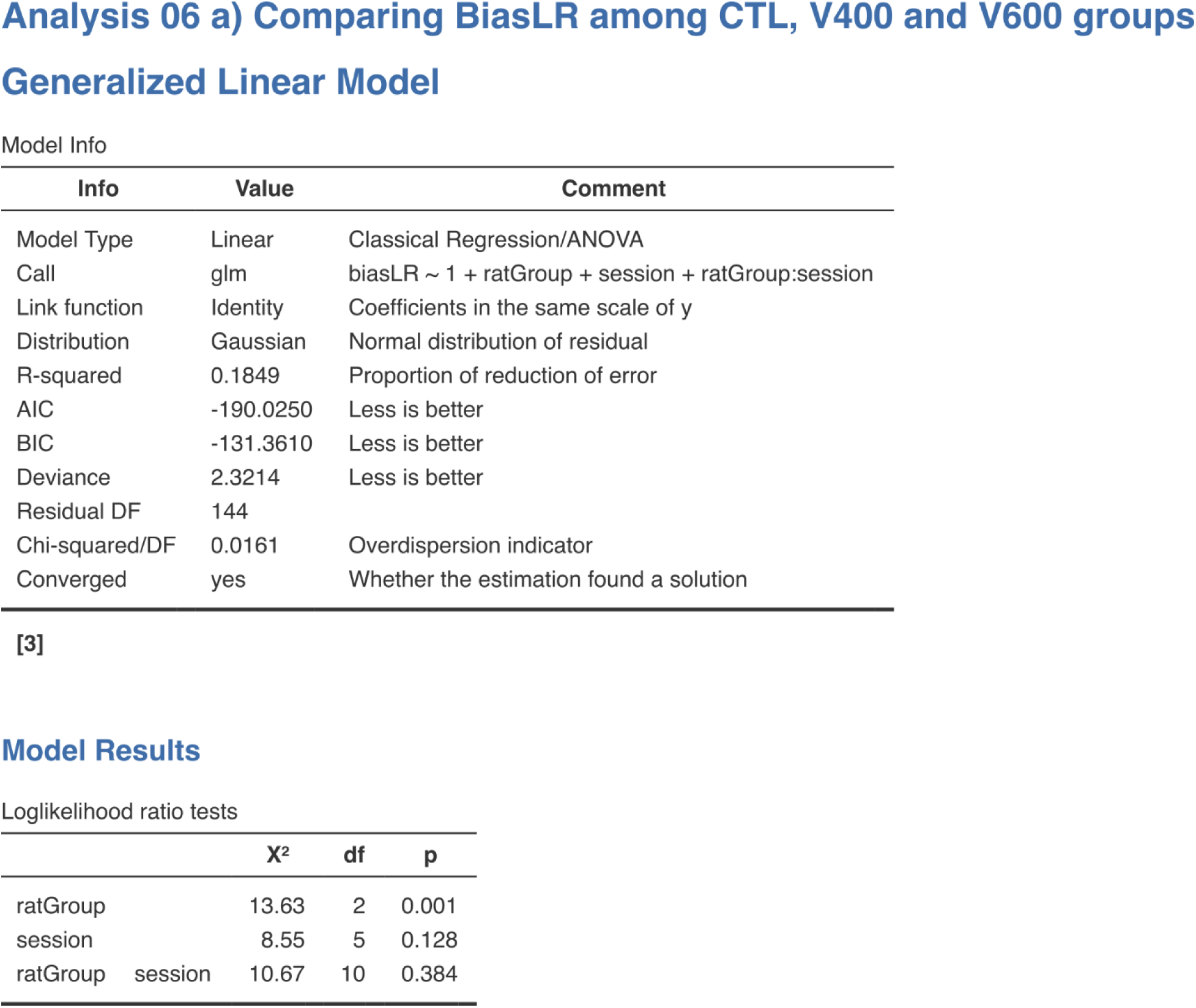
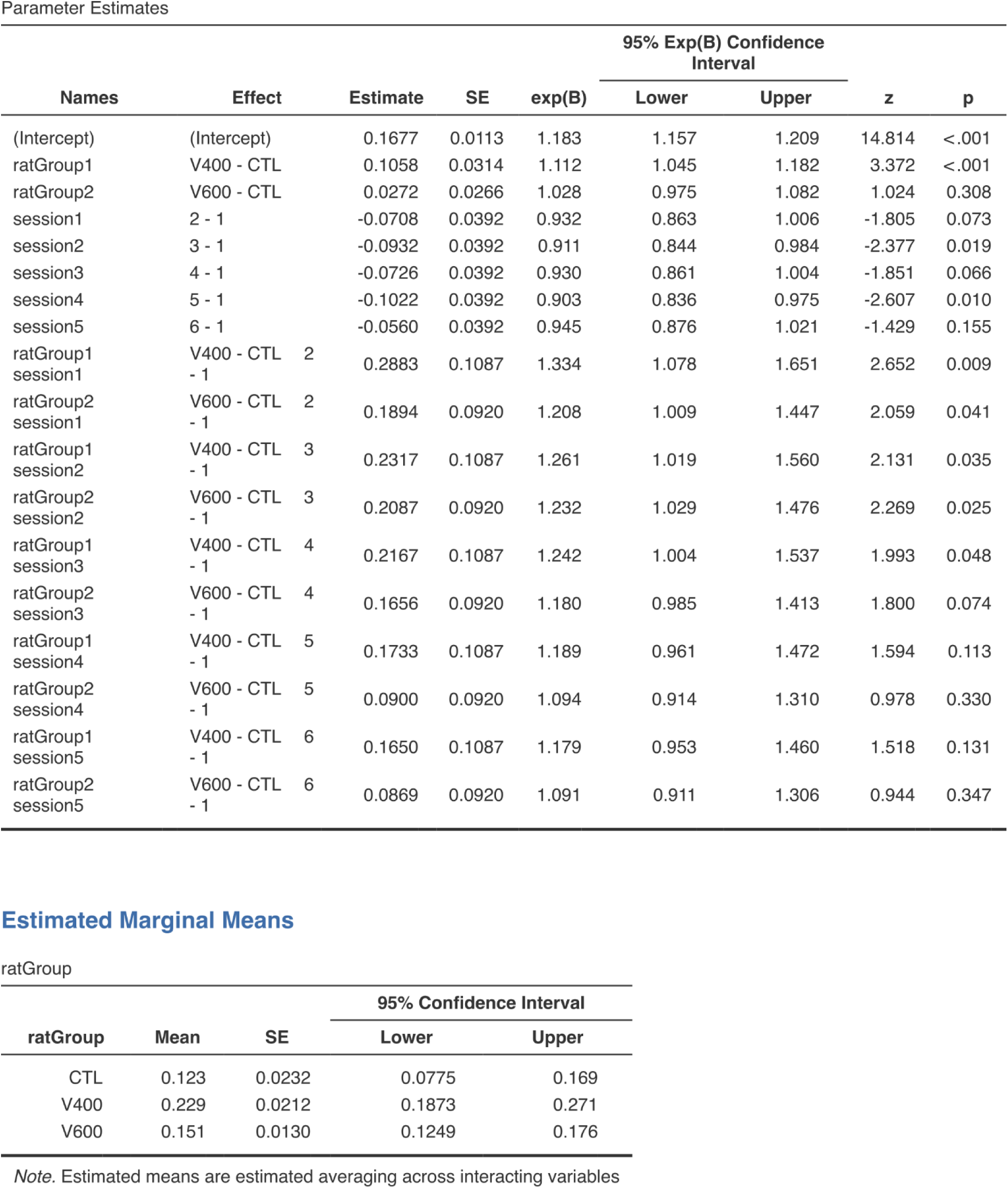
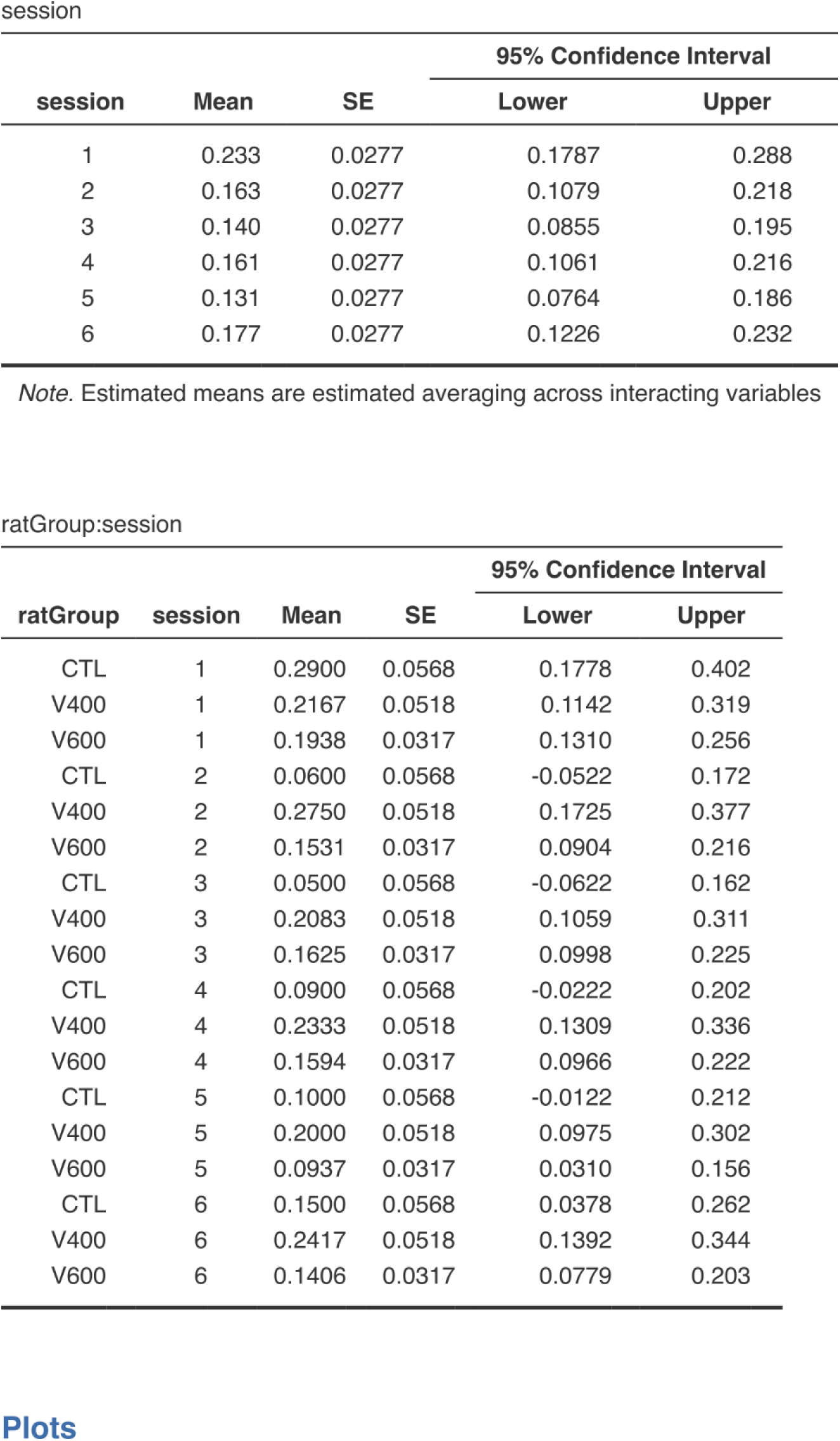
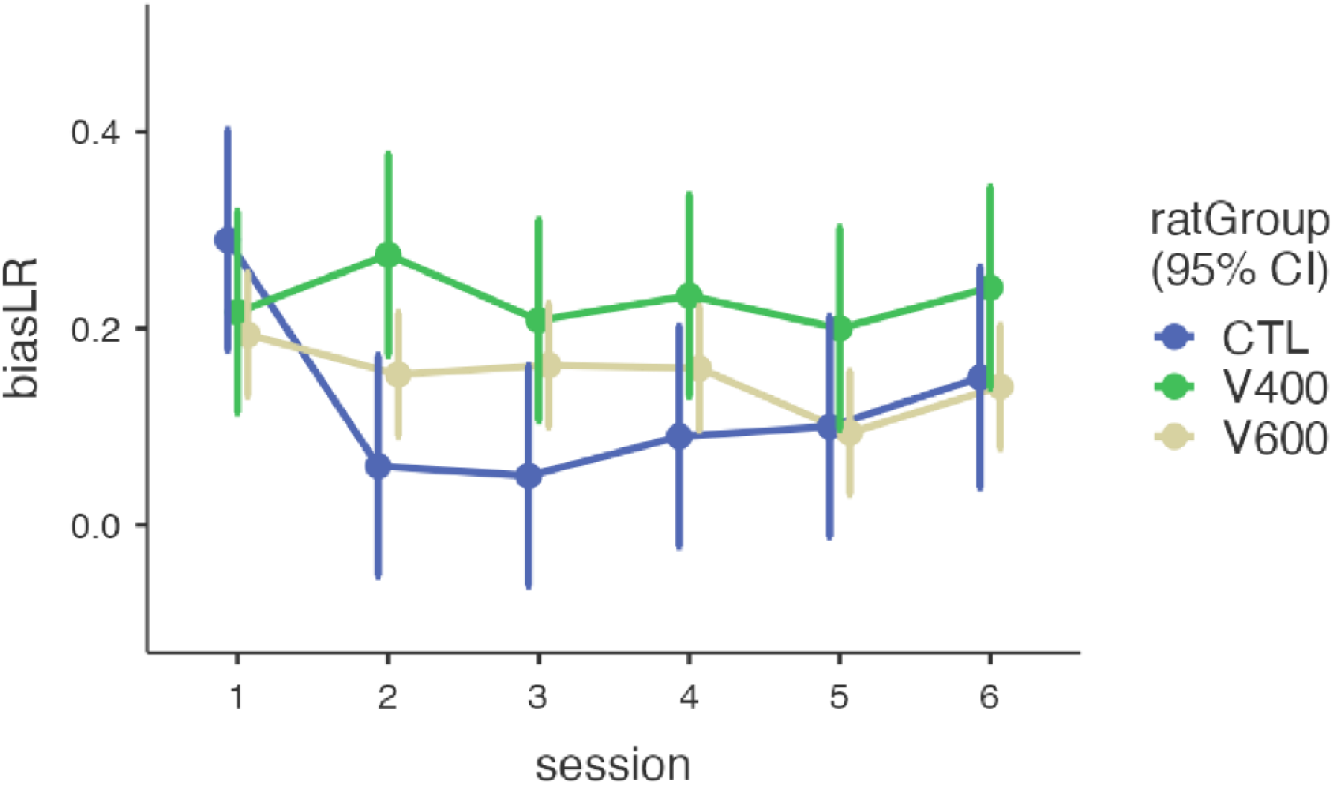
GLMM table for the comparison of Left-right Bias (perseverative behavior) between CTLR, VP400 and VP600 animals.

In contrast, the VP400 group displayed markedly weaker performance. Mean correct trial probabilities were lower, and within-session learning curves were flatter or absent. Only 2.1 out of 6 sessions showed group-level 95% CIs above chance (Figure 2B), and SSM failed to provide stable performance estimates in 2.3 of 6 sessions per animal, on average, reflecting unreliable learning (Figure S1 – data for individual animals; Table 1 – group level analysis and SSM reliability).

The VP600 group, with the largest sample size, showed slightly narrower confidence intervals but significantly lower performance overall. Incremental within-session learning was infrequent, and even in later sessions (S5–S6), group-level performance remained well below CTRL levels (Figure 2C). Only 1.38 of 6 sessions, on average, yielded group-level 95% CIs above chance performance, and SSM produced stable trial-wise estimates in just 3.2 of 6 sessions per animal (Figure S1 – data for individual animals; Figure S2 – Logit and SSM data, with 95% CIs for each individual animal, Table 1 – group level analysis and SSM reliability).

Together, these findings indicate that while VPA-exposed animals are eventually able to reach a performance plateau, their overall success is significantly diminished compared to controls. More importantly, analysis of trial-by-trial dynamics reveals that past trials in VPA animals are poor predictors of future behavior. This inconsistency prevents reliable estimation of learning progression, as only a minority of sessions show performance distinguishable from chance—unlike the more robust and predictable behavior observed in CTRL animals (Figure S1 – data for individual animals; Table 1 – group level analysis and SSM reliability).

### VPA-exposed animals exhibit increased choice perseveration

A common behavioral feature in individuals on the autism spectrum is a diminished ability to promptly perceive or respond to external stimuli, and to flexibly guide behavior based on changing circumstances. One potential manifestation of this is a failure to associate actions with their outcomes—such as associating a specific choice with the subsequent delivery or absence of a reward—and to adjust behavior, thus persevering in one particular trajectory regardless of reward contingencies.

To test this hypothesis, we analyzed raw DNMP data for signs of choice perseveration, operationalized as a persistent Left/Right Bias (LRB) during decision-making. Our analysis revealed a significant group effect: CTRL animals exhibited lower LRB compared to VPA-exposed groups (Figure 3; Table 2; GLMM: χ² = 13.63, *p* < 0.001; Fixed Effect Omnibus test), indicating increased perseverative tendencies in VPA animals.

**Figure 3.**
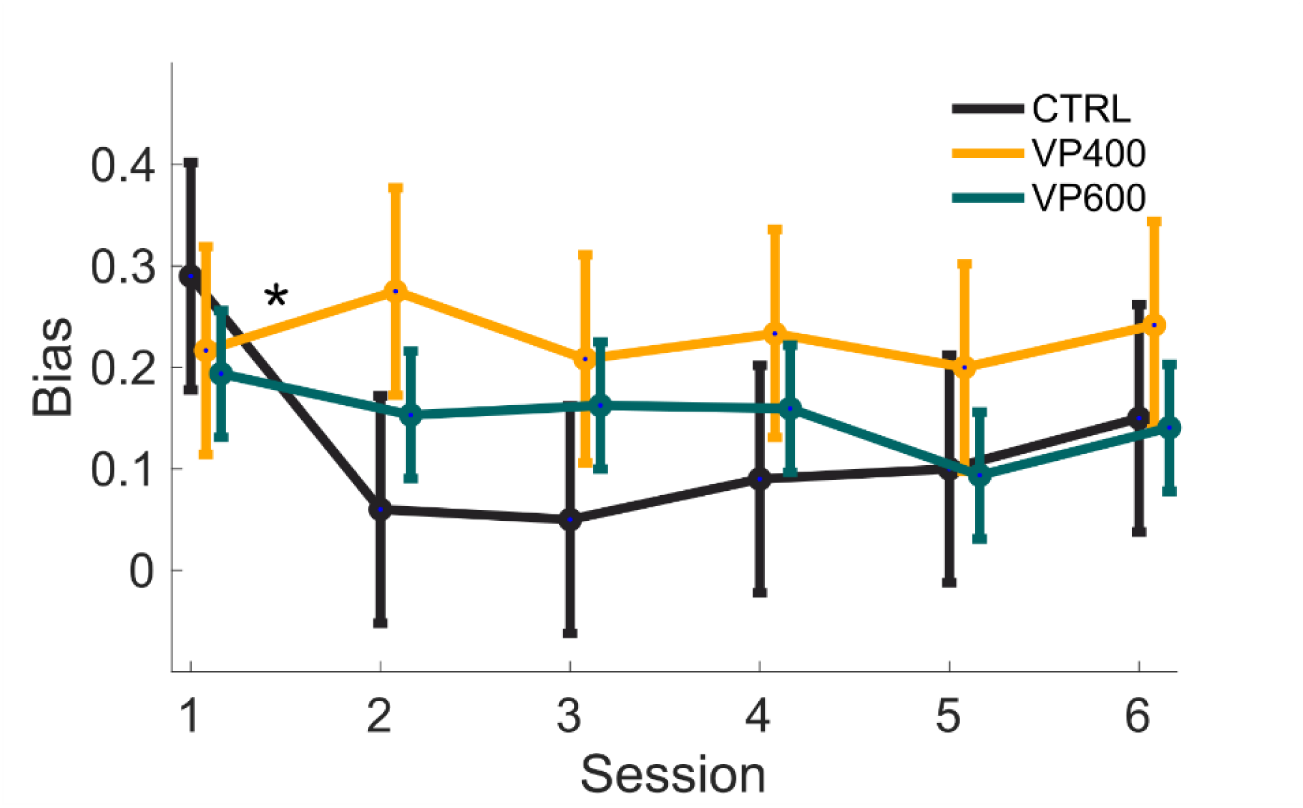
Choice perseveration (left/right bias, LRB) across groups and sessions. Group-level comparison of session-by-session LRB showing increased bias in VP400 and VP600 animals relative to CTRL (GLMM, p<0.001). CTRL animals exhibit a significant reduction of LRB after S1, while VPA groups retain elevated bias across sessions.

Further analysis revealed significant group-by-session interactions. While CTRL animals showed a reduction in LRB after the first DNMP session (S1)—suggesting adaptive learning and behavioral flexibility—both VPA groups (VP400 and VP600) maintained elevated LRB across multiple subsequent sessions. Specifically, compared to S1, significant LRB persistence was observed in VP400 and VP600 animals in sessions S2 (VP400: *p* = 0.009; VP600: *p* = 0.041) and S3 (VP400: *p* = 0.035; VP600: *p* = 0.025). In session S4, only VP400 animals continued to show a significant bias (*p* = 0.048), whereas VP600 animals did not (NS) (Figure 3; Table 2; GLMM: χ² = 13.63, *p* < 0.001; Fixed Effect Omnibus test).

These findings support the notion that prenatal VPA exposure impairs the reduction of perseverative behavior across learning sessions. Unlike CTRL animals, VPA-exposed animals fail to flexibly adapt their choices based on past outcomes, consistent with behavioral inflexibility commonly observed in ASD.

### 40Hz sensory entrainment significantly increases gamma-frequency neural oscillations across brain regions

Functional imaging studies in individuals with ASD have consistently reported underconnectivity in parietal regions ^14,15^, while EEG data reveal disrupted gamma-band oscillations and reduced gamma coherence ^16,17^, as well as diminished fronto-temporal coherence in the 3–6 Hz range. Notably, the latter findings have shown responsiveness to sensory stimulation therapies ^18–20^. In parallel, prenatal VPA exposure in rodent models leads to downregulation of Kv10.2 channel expression in the hippocampus, which results in elevated high-frequency oscillatory activity—a phenotype that can be reversed through compensatory gene overexpression ^21^. These findings parallel MEG data in ASD patients showing increased power across delta, theta, alpha, and broadband gamma bands (20– 120 Hz) in multiple cortical regions, including temporal, parietal, frontal, occipital, and midline areas, with stronger deviations associated with greater symptom severity ^22,23^. Such gamma-frequency dysregulation is likely mediated by deficits in PV+ GABAergic interneurons, whose density is reduced in frontal, motor, and auditory cortices of ASD individuals (reviewed in ^24^). Crucially, abnormal gamma oscillations in ASD patients have been shown to respond to behavioral interventions (e.g., PEERS) ^25,26^ and to transcranial stimulation protocols ^27,28^, resulting in durable improvements in both neural synchrony and behavior during adolescence. Based on this converging evidence, we hypothesized that auditory gamma-frequency sensory entrainment would mitigate the working memory (WM) deficits observed in VPA-exposed animals.

A wealth of evidence suggests that cortical networks in humans and animals can be reliably entrained to external stimuli—particularly light ^29^ and sound ^30,31^, delivered at gamma frequencies relevant to cognition. Recent rodent studies further demonstrate that 40 Hz gamma entrainment, via either visual or multimodal (visual and auditory) stimuli, improves cognitive function and induces broad changes in neuropathology and neuroplasticity in Alzheimer’s disease models, with persistent effects following the intervention^32–36^.

To test whether 40 Hz auditory stimulation could induce similar effects in the VPA rodent model, we recorded local field potentials (LFPs) in two animals—one VP600 and one CTRL—each implanted with multi-tetrode hyperdrives targeting the hippocampus and cortical regions. During recording sessions, animals received 40 Hz patterned auditory stimulation, and their LFPs were analyzed using Fast Fourier Transform (FFT) and multi-taper spectral analysis ^37^. Our analyses confirmed that auditory entrainment at 40 Hz enhances gamma-band power, as well as activity in other relevant frequency bands such as theta and beta (Figure 4). Importantly, we observed sustained changes in oscillatory power that persisted beyond the period of active stimulation. These results demonstrate that 40 Hz auditory stimulation effectively entrains neural circuits in awake, behaving rats, and induces lasting changes in neural dynamics, providing a possible mechanistic basis for therapeutic effects.

**Figure 4.**
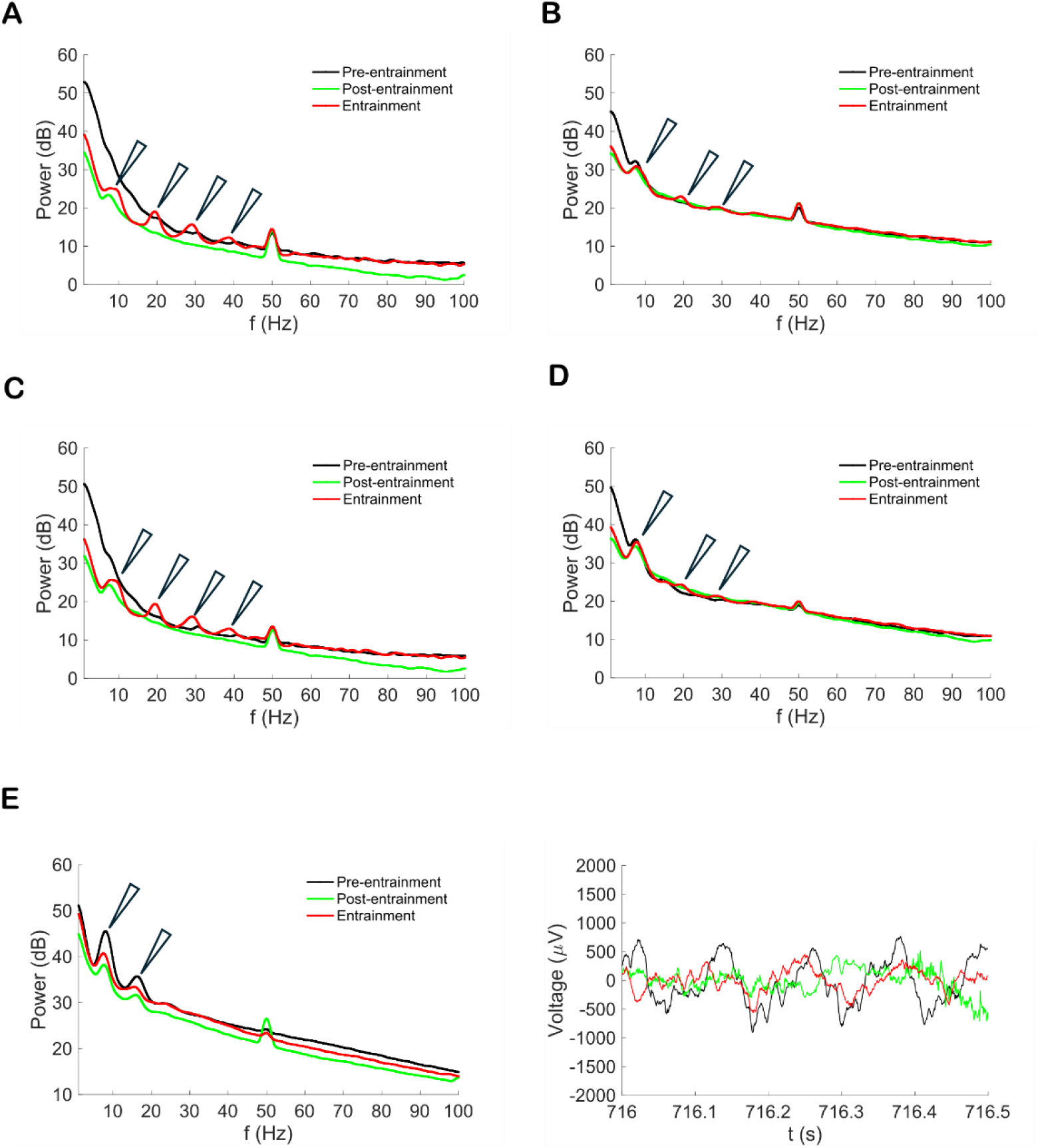
Neural entrainment to 40 Hz auditory stimulation in awake behaving rats. Power Spectral Density plots showing power changes at physiologically relevant frequencies under entrainment, often enduring beyond the entrainment stimulus delivery. (A) Auditory cortex, (B) Posterior Parietal cortex, (C) LPMR Thalamus, (D) Medial occipital secondary (Oc2M) cortex, (E) Hippocampal CA1 subfield during DNMP trials, raw data samples of the right

### Auditory gamma entrainment restores working memory performance in VPA-exposed rats

Building on our finding that 40 Hz auditory stimulation reliably alters neural activity in both control and VPA-exposed animals, we next tested the therapeutic potential of this intervention on working memory (WM) deficits observed in VPA600 animals. To this end, a separate cohort of VPA600 animals was run through three consecutive sets of six DNMP sessions (Figure 5B). The middle block of sessions included auditory entrainment, delivered continuously throughout each behavioral session using a 40 Hz-patterned sound stimulus.

**Figure 5.**
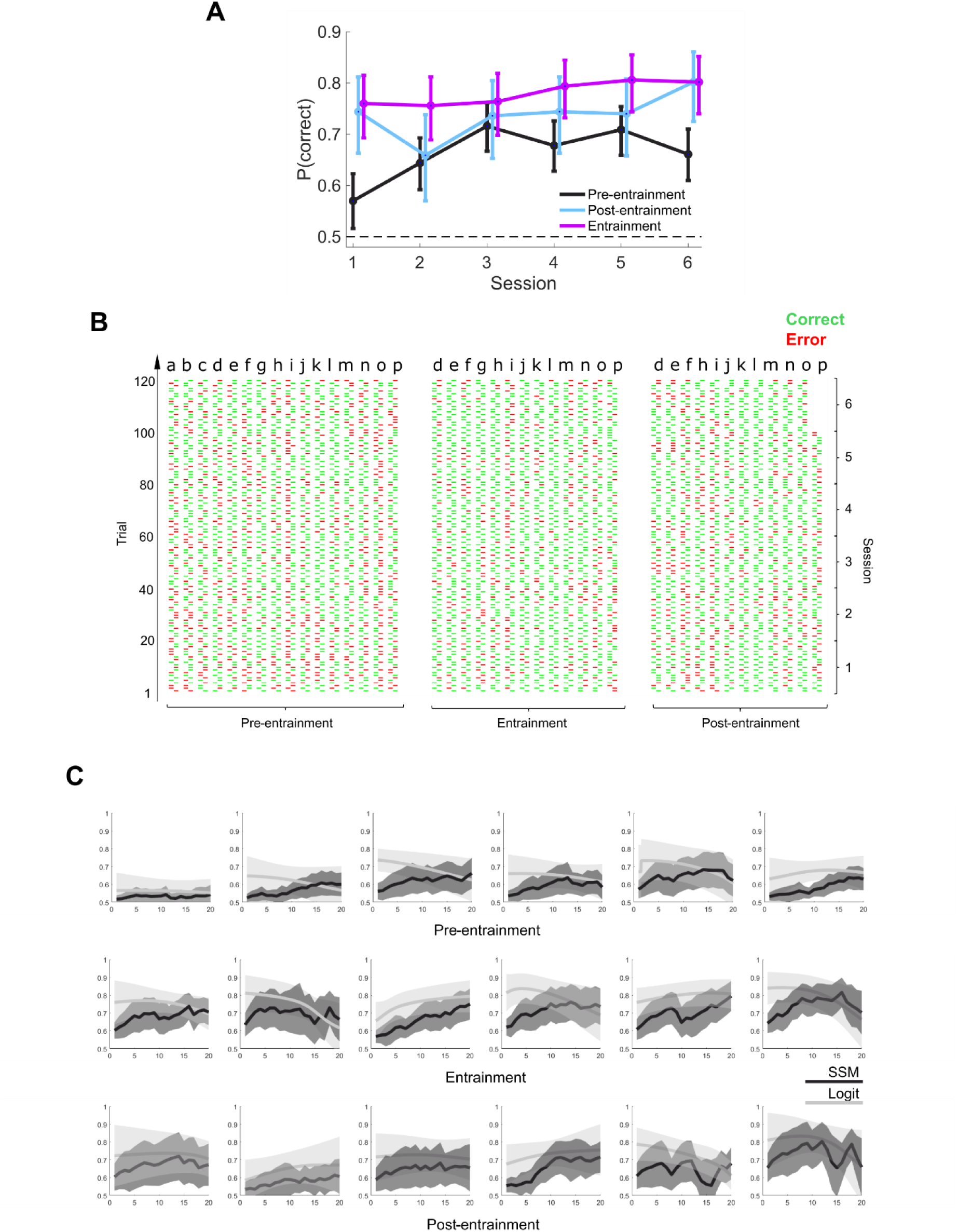
Sensory entrainment (SE) rescues WM deficits in VPA600 animals. **(A)** Group-level performance comparison during showing improved correct response rates during entrainment when compared to pre-entrainment (p<0.001), enduring into post-entrainment. **(B)** Raw behavioral data from individual animals across all sessions (similar to Figure 1C), during the three epochs of the entrainment experiment: pre-entrainment (sessions 1–6), entrainment (sessions 7–12), and post-entrainment (sessions 13–18). **(C)** Overlaid BLR (light shade) and SSM (strong shade) mean +/-95%CI, trial-wise estimates for VP600 animals, for the 6 sessions, across the three epochs of the experiment (similar to Figure 2). Entrainment epoch shows CTRL-like increases in trial-by-trial probability of correct responses. This improvement outlives the delivery of the entrainment stimulus, during post-entrainment.

As shown in Figure 5A, auditory entrainment produced a robust and statistically significant improvement in DNMP performance across sessions. Specifically, performance during entrainment sessions was markedly enhanced compared to baseline: the average probability of a correct trial increased from 0.66 pre-entrainment to 0.78 during entrainment (GLMM, *p*<0.001). Strikingly, this performance was statistically indistinguishable from that of the CTRL animals (GLMM, *p* = 0.497), suggesting a full rescue of the WM deficit.

These findings were further supported by trial-by-trial analyses using both Binary Logistic Regression (Logit) and State-Space Modeling (SSM) (Figure 5C). During the entrainment sessions, VPA600 animals displayed trial-wise performance dynamics that closely resembled those of CTRL animals, with both methods showing increased and more consistent estimates of the probability of correct responses. These results strongly suggest that auditory gamma entrainment enhances WM performance to levels comparable to those of unexposed controls. Please refer to Figure S1 for individual animalś raw data and analysis.

### VPA-exposed animals exhibit trajectory-perseverative behavior

Lastly, we investigated whether sensory entrainment (SE) also modulates perseverative behavior, operationalized as Left/Right Bias (LRB), in VPA-exposed animals. While LRB differences across pre-entrainment, entrainment, and post-entrainment sessions in the VP600 group were not statistically significant (Figure 6A, GLMM, *p* = 0.068), the data suggested a potential trend worth exploring further.

**Figure 6.**
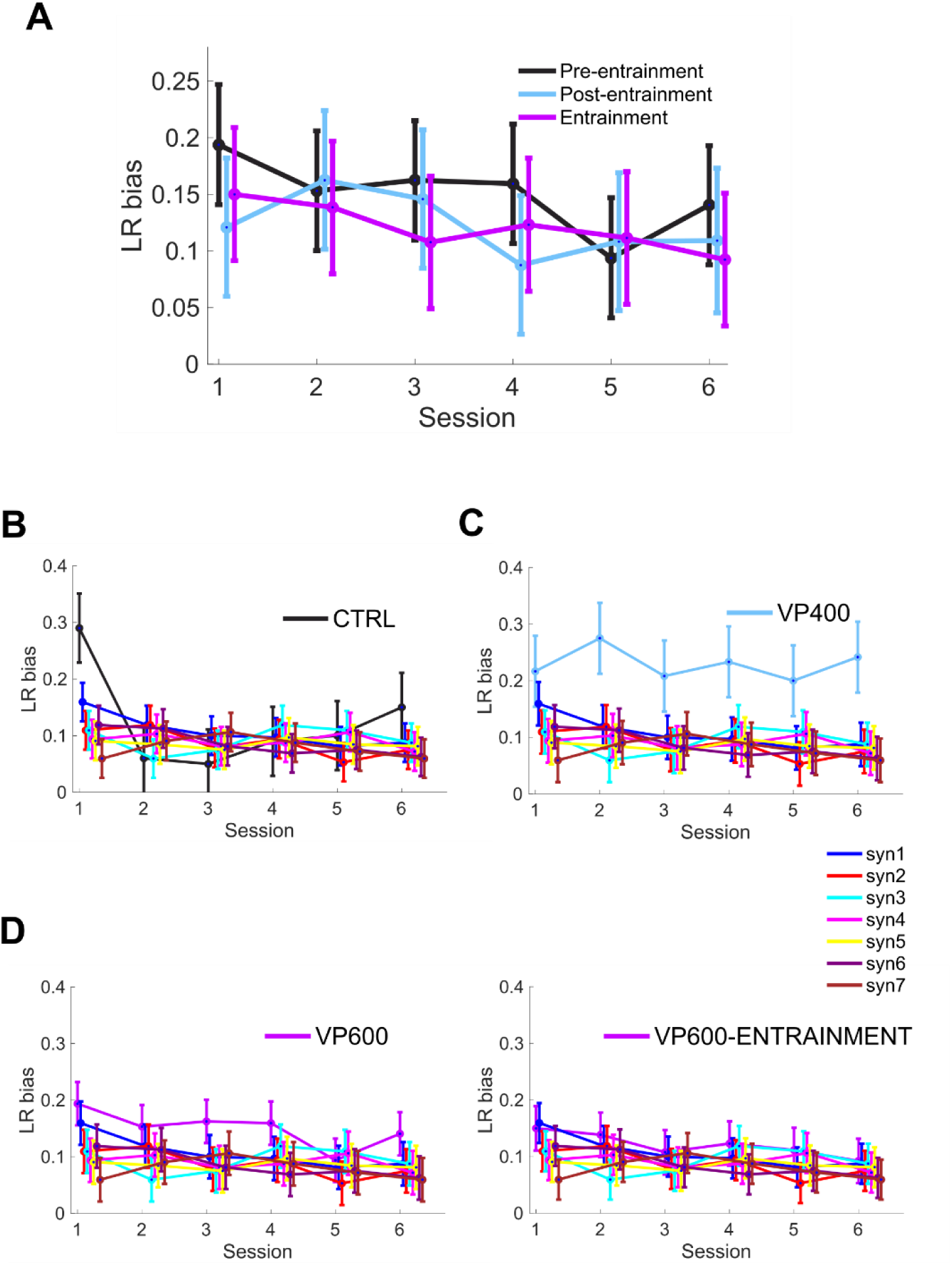
Sensory entrainment reduces LRB in VPA600 animals. **(A)** LRB scores across pre-entrainment, entrainment, and post-entrainment epochs in VP600 group. **(B)** Comparison of CTRL levels with synthetic data shows a high LRB on Session1 followed by much lower levels, indistinguishable from unbiased simulations. **(C, D-left panel)** LRB levels in VP400 and VP600 animals is significantly higher than that of synthetic data. **(D-right panel)** LRB levels in VP600 animals decreases during entrainment, compared to synthetic data.

To contextualize this behavior and assess whether LRB presence or absence can emerge by chance during DNMP training, we used the original trial structure (i.e., blocked arm, trial count, group sizes, and sessions) to create seven replicates in which choice responses were randomly assigned from an unbiased distribution, allowing us to establish a comparison baseline, and thus assess the emergence and persistence of LRB across treatment groups (CTRL, VP400, and VP600 and VP600-entrainment).

Compared to synthetic data, CTRL animals exhibit LRB restricted to the first session, after which it becomes similar to levels resulting from synthetic data, in agreement with the previous data (Figure 6B). VP400 animals exhibited high LRB levels relative to the synthetic control group (Figure 6C, GLMM, group effect *p* < 0.001; session-by-group interaction *p* = 0.597, n.s.), with only modest attenuation across sessions (*p* = 0.064). VP600 animals showed similar results (GLMM, group effect *p* < 0.001; session-by-group interaction *p* = 0.367, n.s.), with a significant attenuation across sessions (*p* = 0.004).

Notably, sensory entrainment reduced LRB in VP600 animals (GLMM, entrainment effect *p* = 0.004; session effect *p* = 0.003; session-by-group interaction *p* = 0.463, n.s.). During entrainment sessions, LRB in VP600 animals was statistically indistinguishable from that of one of the synthetic replicates (*p* = 0.229) and significantly reduced compared to baseline sessions (individual session *p* values ranging from 0.001 to 0.013). These findings indicate that auditory gamma entrainment may transiently normalize perseverative tendencies, in addition to rescuing WM impairments.

## Discussion

We have shown that prenatal exposure to Valproic Acid (VPA), a well-established rodent model of ASD, induces robust and persistent deficits in SWM performance in a delayed non-match to place (DNMP) task. Importantly, we found that gamma-frequency sensory entrainment (SE) not only modulated neural activity in a frequency-specific and lasting manner but also rescued both cognitive and behavioral deficits in VPA-exposed animals.

### Impaired WM and Learning Dynamics in VPA-Exposed Rats

Across sessions, VPA animals exhibited significantly poorer performance in the DNMP task compared to controls, with both VP400 and VP600 groups showing slower learning curves and reduced final performance levels. However, this impairment was not simply a reflection of delayed acquisition. Our trial-by-trial analyses using Binary Logistic Regression (BLR) and State Space Modeling (SSM) revealed that VPA animals lacked consistent within-session improvements and showed noisy or unstable probability estimates of correct responses, suggesting a fundamental disruption in the integration of recent experience to guide future choices.

This inability to progressively improve performance likely reflects underlying disruptions in distributed networks critical for goal-directed behavior (GDB), where allocentric representations must be reinstated from memory and translated into egocentric action plans ^38–41^. This transformation relies on synchronized activity across the hippocampus (HIPP), posterior parietal cortex (PPC), retrosplenial cortex (RSC), and prefrontal areas— circuits known to be affected in ASD. Specifically, anatomical studies show increased local neuronal density and reduced long-range white matter connections in ASD brains, including decreased myelination and structural thinning of PPC, temporal, and frontal cortices ^42–47^. These features are consistent with our behavioral findings and lend support to the hypothesis of impaired distributed coordination in ASD models.

### Perseverative Behavior as a Signature of Cognitive Rigidity

Perseverative responding, quantified here as left/right bias (LRB), was significantly elevated in VPA-exposed animals and showed limited improvement with training - contrasting with the rapid reduction of LRB observed in control animals after session 1. This suggests that VPA animals struggle to update behavior considering changing contingencies, a form of cognitive rigidity common in ASD.

Such rigidity may stem from failures to integrate reinforcement signals with prior action choices. Indeed, ASD patients often show reduced flexibility in reversal learning tasks and diminished engagement of prefrontal-striatal circuits responsible for adaptive choice updating. Moreover, abnormal connectivity patterns—such as reduced long-range frontoparietal coherence and increased local connectivity—have been extensively reported in ASD populations, potentially impairing the ability to coordinate activity across functionally-relevant brain areas.

### Restoring Function Through Gamma-Frequency Sensory Entrainment

Gamma-band oscillations (30–80 Hz) are thought to play a key role in coordinating distributed neural activity and enabling top-down control, attention, and WM maintenance. Disturbances in gamma rhythms are among the most robust electrophysiological findings in ASD, with reports of reduced gamma coherence and impaired phase locking to rhythmic stimuli in both EEG and MEG studies ^14–17^. Notably, gamma oscillations are tightly regulated by PV+ GABAergic interneurons, which are reduced in number in several cortical regions in ASD ^24^.

In our study, auditory 40 Hz stimulation effectively entrained neural activity in behaving rats, enhancing not only gamma but also theta and beta band power. These changes persisted beyond stimulus exposure, suggesting durable network-level modulation. Importantly, when VPA600 animals were subjected to gamma entrainment during DNMP sessions, we observed a complete rescue of WM performance—rendering their performance statistically indistinguishable from that of controls. Trial-wise modeling further confirmed that entrained animals exhibited stabilized learning curves and trial probabilities, matching CTRL-like profiles.

Moreover, SE also attenuated perseverative bias in VPA600 animals, reducing LRB to levels indistinguishable from unbiased synthetic controls. This suggests that gamma entrainment not only improves task performance but may restore behavioral flexibility by re-engaging circuits underlying adaptive decision-making.

### Mechanistic and Translational Implications

The beneficial effects of gamma entrainment observed here echo findings from Alzheimer’s disease (AD) models, where 40 Hz sensory stimulation rescues cognitive performance and alters neuropathology. Similar interventions in ASD patients—such as transcranial stimulation and behavioral training - have shown promising results in normalizing gamma activity and improving cognitive and social function.

Our results suggest that gamma entrainment may counteract the “local-over-connectivity, long-range underconnectivity” pattern present in ASD by promoting coherent, distributed neural activity. This could facilitate the reinstatement of allocentric memory traces and enable more effective goal-directed action planning. Furthermore, entrainment may modulate PV+ interneuron function, stabilizing oscillatory regimes critical for SWM and behavioral flexibility.

Our findings demonstrate that prenatal VPA exposure impairs SWM and induces perseverative behavior in rats, reflecting key features of ASD. Trial-by-trial modeling reveals unstable learning dynamics in VPA animals, and the presence of persistent choice bias. However, auditory gamma-frequency sensory entrainment restores both cognitive performance and behavioral flexibility, offering a promising non-invasive intervention to modulate large-scale brain activity. These results support the hypothesis that rhythmic sensory stimulation may rescue functional deficits in ASD by restoring oscillatory coherence and long-range network coordination.

## Acknowledgements

We are indebted to GIMM’s Bioimaging, Comparative Pathology, and Rodent facilities for critical help. We want to thank all the members of the Lopes Lab for fruitful discussions and help.

FCT granted a PhD Fellowship (BD-202009547) to JC, and an Exploratory Grant (IF/00201/2013), an IMM Director’s Fund Award, an Investigator FCT Position at IMM-JLA (IF/00201/2013), a Research Grant (PTDC/MED-NEU/29325/2017), the 2022.03699.CEECIND Coordinator Investigator contract to LVL, and the 2022.00811.CEECIND Principal Investigator contract to MR.

## Author contributions

JC and MR designed the experiments, JC, ML and CM conducted the experiments, JC and MR analyzed the data, and JC, LL and MR wrote the manuscript.

## Methods

### Lead Contact and Materials Availability

Further information and requests for resources and reagents should be directed to and will be fulfilled by the Lead Contact, Miguel Remondes, DVM, PhD. This study did not generate new unique reagents.

### Experimental Model Generation

All procedures were performed in accordance with EU and Institutional guidelines. Female and male rats aged 3-6 m, were obtained from Charles River France, kept in a 12hr light/dark cycle, fed *ad libitum*, housed in groups until mating cycles begun.

### TIMED-MATING PROTOCOL

Pairs of males and females of the same age were formed, and single-caged.for 48 hours to reduce stress and prepare females for estrus synchronization. After the separation period, cages were swapped between animals of the same pair to expose females to the scent of the male thus-timing estrus. Estrus was detected using periodic vaginal electrical impedance measurements. Once in peak estrus the pair was rejoined and allowed to mate overnight for a period of 8 hours. In the following morning, the pair was separated, and the female was inspected for the presence of a vaginal plug, in which case the female was labeled’possibly pregnant’ at day 0.5 (E0.5, embryonic day 5), and weighted daily thereafter.

### VPA TREATMENT

Twelve days later, on E12.5, pregnant females received an IP injection of either saline (CTRL), 400 mg/ Kg (VP400), or 600 mg/ Kg (VP600) of valproic acid sodium salt (VPA, Sigma P4543) diluted in saline, and were housed individually. The day the pups were born was recorded as post-natal day 0 (P0), and the bedding was left undisturbed for 7 days. After weaning (P21), pups were housed in groups of up to 3 rats per cage, and assigned experimental groups according to treatment, CTRL (saline), VP400 (400 mg/ kg) and VP600 (600 mg/ kg).

### DELAYED NON-MATCHING TO PLACE TASK

In preparation for the DNMP task, the above animals (now 6 month old) were food-restricted to 85% BW. The DNMP task was conducted in a T-maze featuring a starting platform (25×30 cm), a central corridor (170 cm) and two lateral exit arms (88 cm each) and whose ends a chocolate reward was placed. An external opaque resting box (57×39×42cm) was used to temporarily house the rat during the inter-trial maze setup. Each trial consisted in three runs: (1) Sample, (2) Delay and (3) Test. (1) Sample Run: one of the lateral arms (randomly defined) was baited with reward while the other arm was blocked. (2) After the rat visited the baited arm and consumed the reward, it was taken out of the maze and placed in the resting box for 20 seconds (the Delay). (3) Test Run: once the delay period had elapsed, the animal was returned to the maze, both whose lateral arms were accessible again. The rat was now rewarded for selecting the arm non-matching the one baited before (during Sample). Test trials were labelled accordingly Correct or Error. All sessions were recorded using a Flea 3 PointGreyTM at 30 fps acquisition rate (top view) mounted on a cable tray fixed on the ceiling above the T-maze and stored in the acquisition computer running Bonsai software.

### Sound Entrainment Protocol

During the entrainment sessions a synthetically generated sound, consisting of a carrier tone of 8 kHz amplitude-modulated at 40 Hz, was continuously delivered by a speaker (Blow BT-950) placed inside the room, above and aligned with the T-maze central corridor, and modified until the Sound Pressure Level (SPL) reached 70 dB at the maze level.

### Electrophysiological Recordings

Local field potential recordings were performed in awake-behaving CTRL and VP600 rats, using a multi-tetrode drive chronically implanted. The data was acquired at 20K sampling rate using Intan RHD2164 amplifier boards connected to the Open-Ephys board, as described earlier ^48^.

### Behavioral Data Acquisition and Statistical Analysis

Trial outcomes were recorded (“correct” or “error”) and classified as 0 or 1 across all groups, sessions and animals. Pooled, and, trial-by-trial data were analyzed using Matlab and Jamovi. The individual animals’ behavior curve were computed for each individual session, from the individual trial outcomes, using two complementary methods: 1) State Space Model (SSM) ^49^ applied to estimate the individual subject learning curve using binomial expectation maximization and 2) a predictive binary logistic regression model. In both cases we computed the 95% confidence interval displayed in the main and supplementary plots, in both individual animals and groups. Generalized Linear Mixed Models (GLMM) were employed to analyze the behavioral outcome results. Model selection was based on the Akaike Information Criterion (AIC), with the chosen model being the one that converged and exhibited the lowest AIC value. To assess performance differences between VPA and CTRL groups, we used a GLMM with a binomial distribution and logit link function, with individual subjects as the random effects variable. The same GLMM, followed by a Bonferroni post-hoc test, was conducted to analyze the ratio of correct trials between groups during the experiment.

