## Supplemental Figure 1 for "Auditory Gamma-Frequency Entrainment Abolishes Working Memory Deficits in a Rodent Model of Autism"

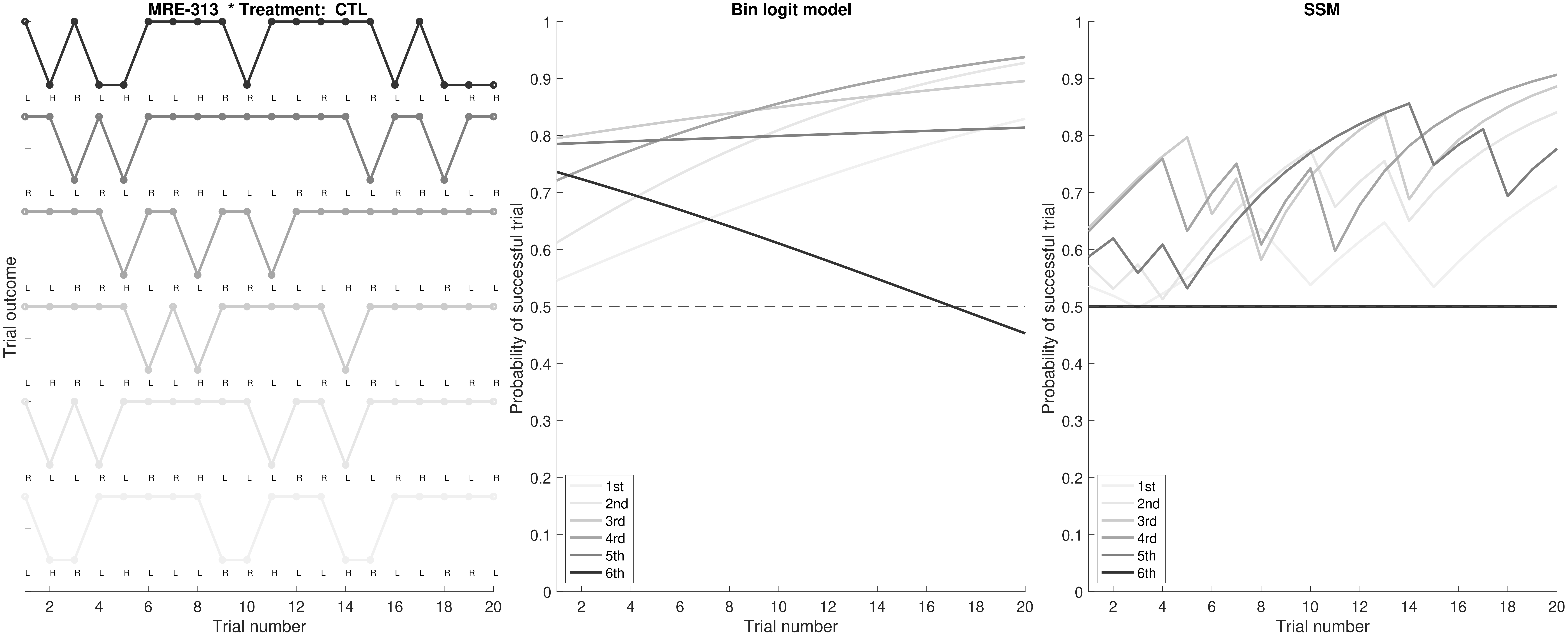

MRE-313 * Treatment: CTL
Trial outcome
L R R L R L L R R R L L R L R L L L R R
R L L R L R R R L L L R R R L R R L L R
L L R R R L R L R R L L L R R L L R L L
L R R L R L L R R R L L R L R L L R R R
R L L R L R R R L L L R R R L R R L L R
L R R L R L L L R R R L L R R L L R R L
2 4 6 8 10 12 14 16 18 20
Trial number
Bin logit model
Probability of successful trial
0 0.1 0.2 0.3 0.4 0.5 0.6 0.7 0.8 0.9 1
1st
2nd
3rd
4rd
5th
6th
2 4 6 8 10 12 14 16 18 20
Trial number
SSM
Probability of successful trial
0 0.1 0.2 0.3 0.4 0.5 0.6 0.7 0.8 0.9 1
1st
2nd
3rd
4rd
5th
6th
2 4 6 8 10 12 14 16 18 20
Trial number

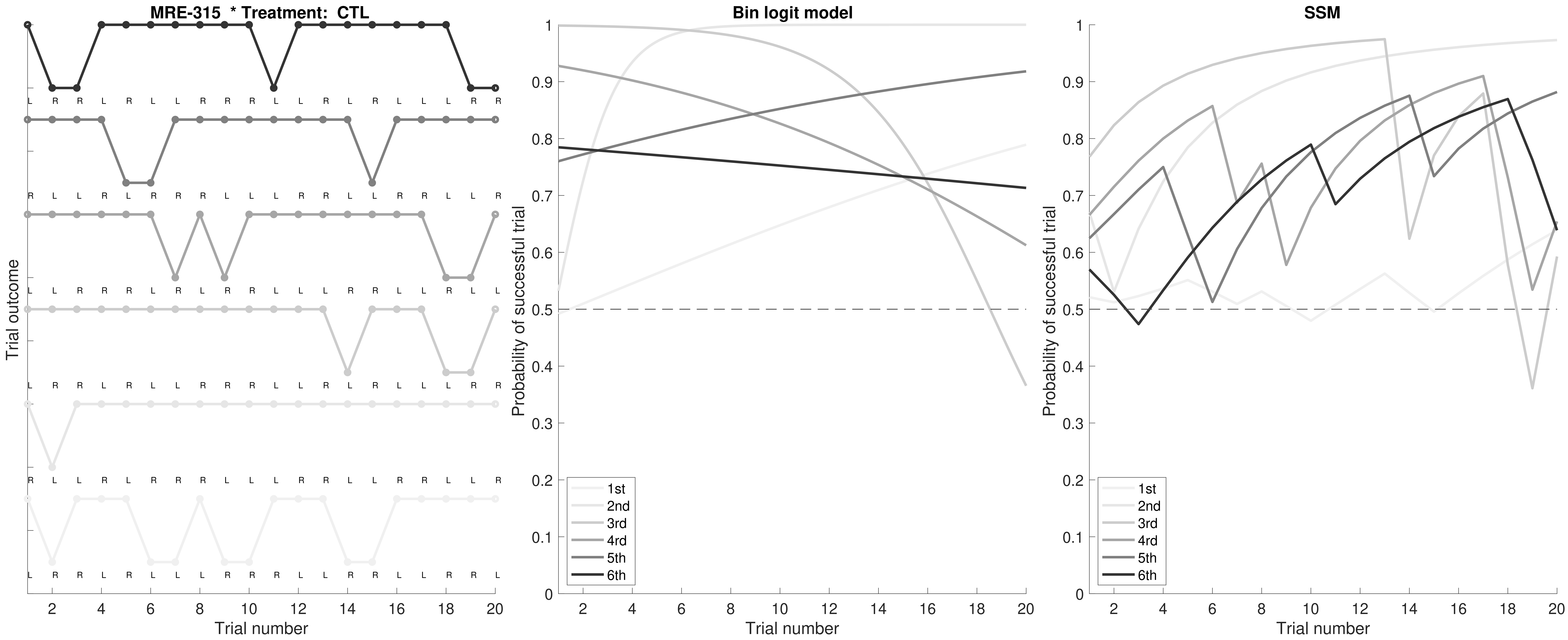

MRE-315 * Treatment: CTL
Trial outcome
Trial number
Bin logit model
Probability of successful trial
Trial number
SSM
Probability of successful trial
Trial number
1st
2nd
3rd
4rd
5th
6th

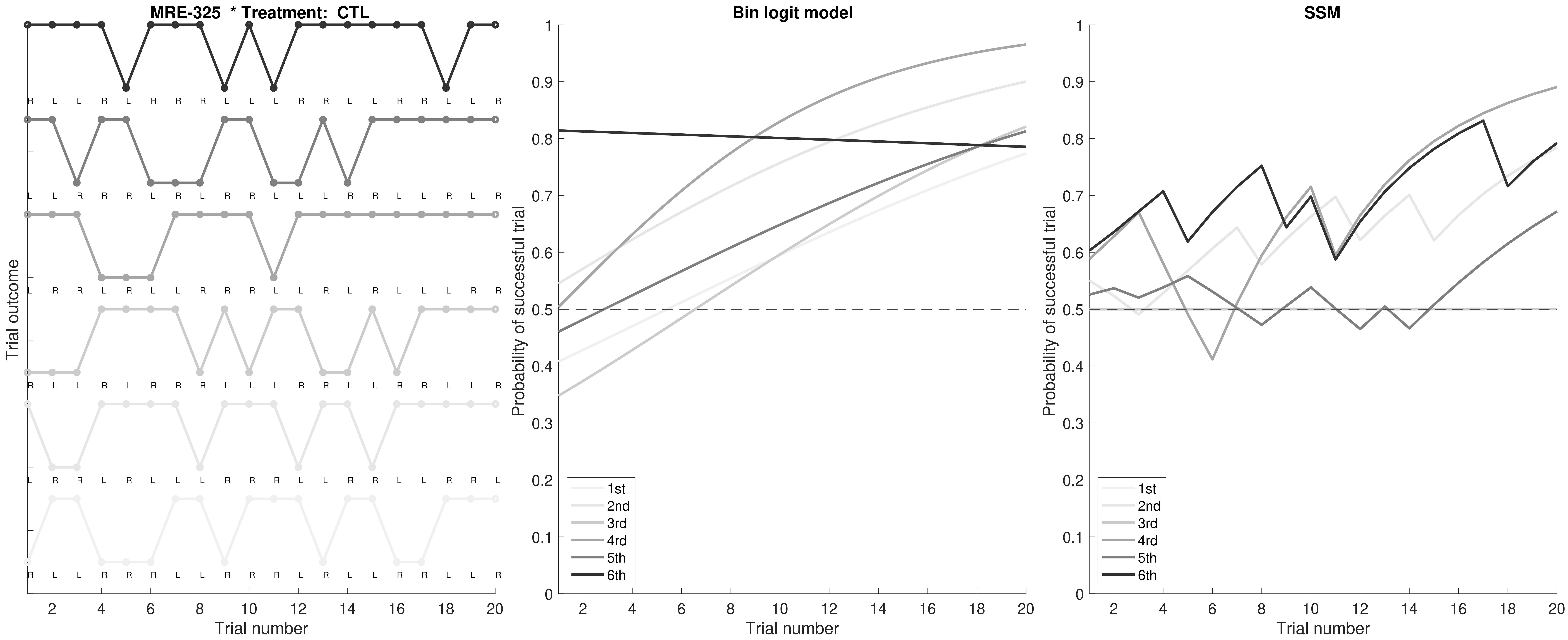

MRE-325 * Treatment: CTL
Trial outcome
Trial number
Bin logit model
Probability of successful trial
Trial number
SSM
Probability of successful trial
Trial number
1st
2nd
3rd
4rd
5th
6th

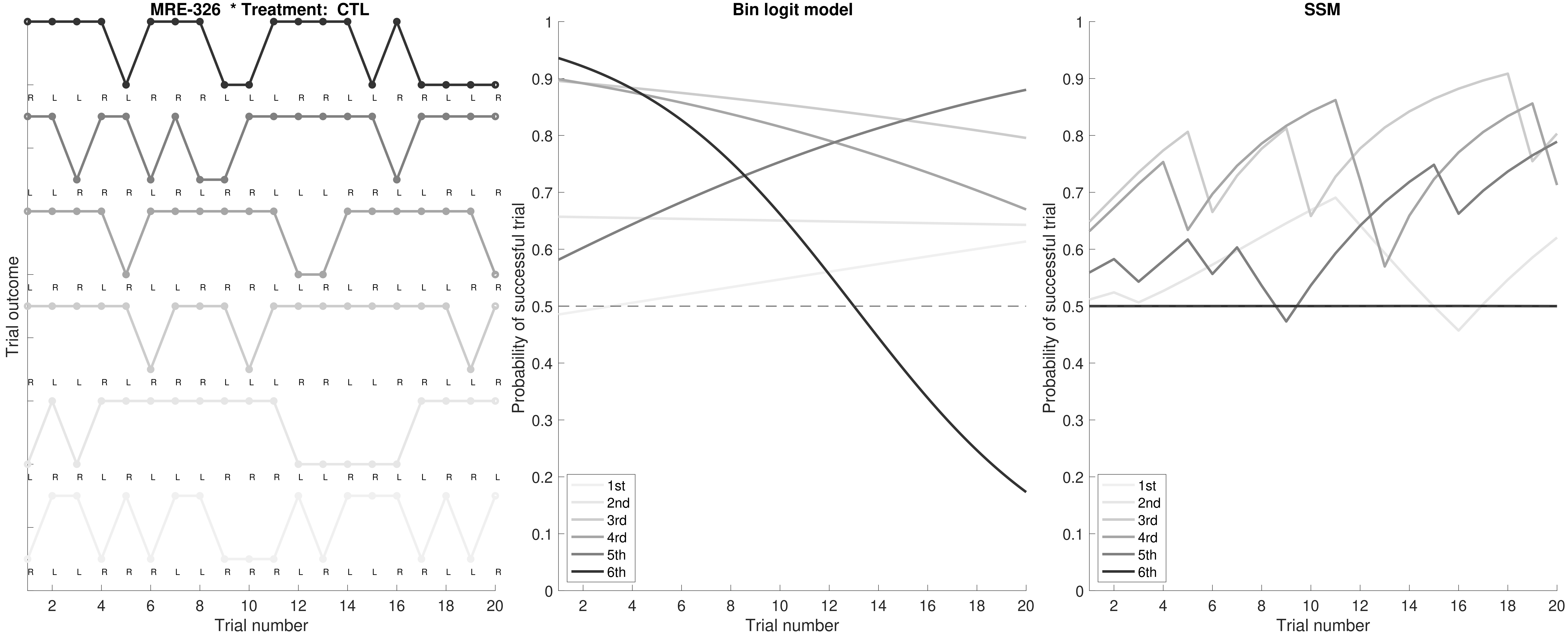

MRE-326 * Treatment: CTL
Trial outcome
Trial number
Bin logit model
Probability of successful trial
Trial number
SSM
Probability of successful trial
Trial number
1st
2nd
3rd
4rd
5th
6th

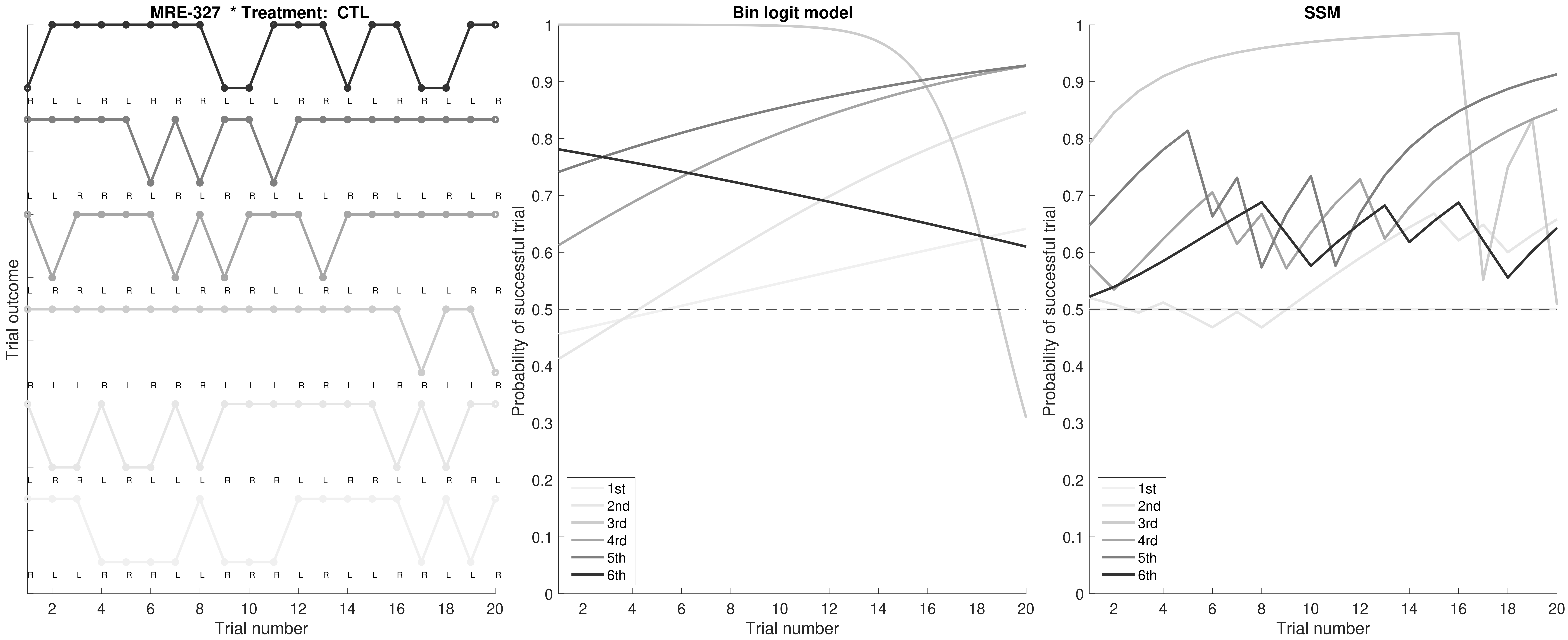
MRE-327 * Treatment: CTL
Trial outcome
Trial number
Bin logit model
Probability of successful trial
Trial number
1st
2nd
3rd
4rd
5th
6th
SSM
Probability of successful trial
Trial number
1st
2nd
3rd
4rd
5th
6th

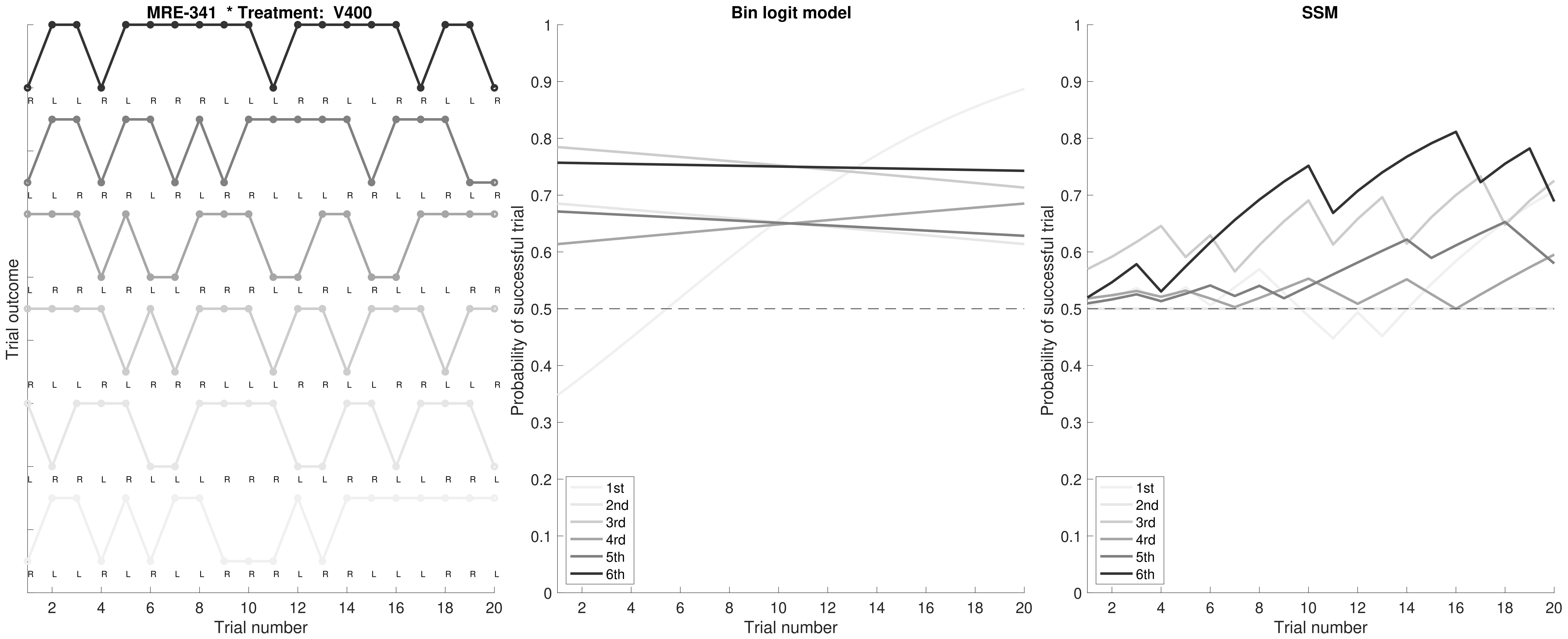

MRE-341 * Treatment: V400
Trial outcome
Trial number
Bin logit model
Probability of successful trial
Trial number
1st
2nd
3rd
4rd
5th
6th
SSM
Probability of successful trial
Trial number
1st
2nd
3rd
4rd
5th
6th

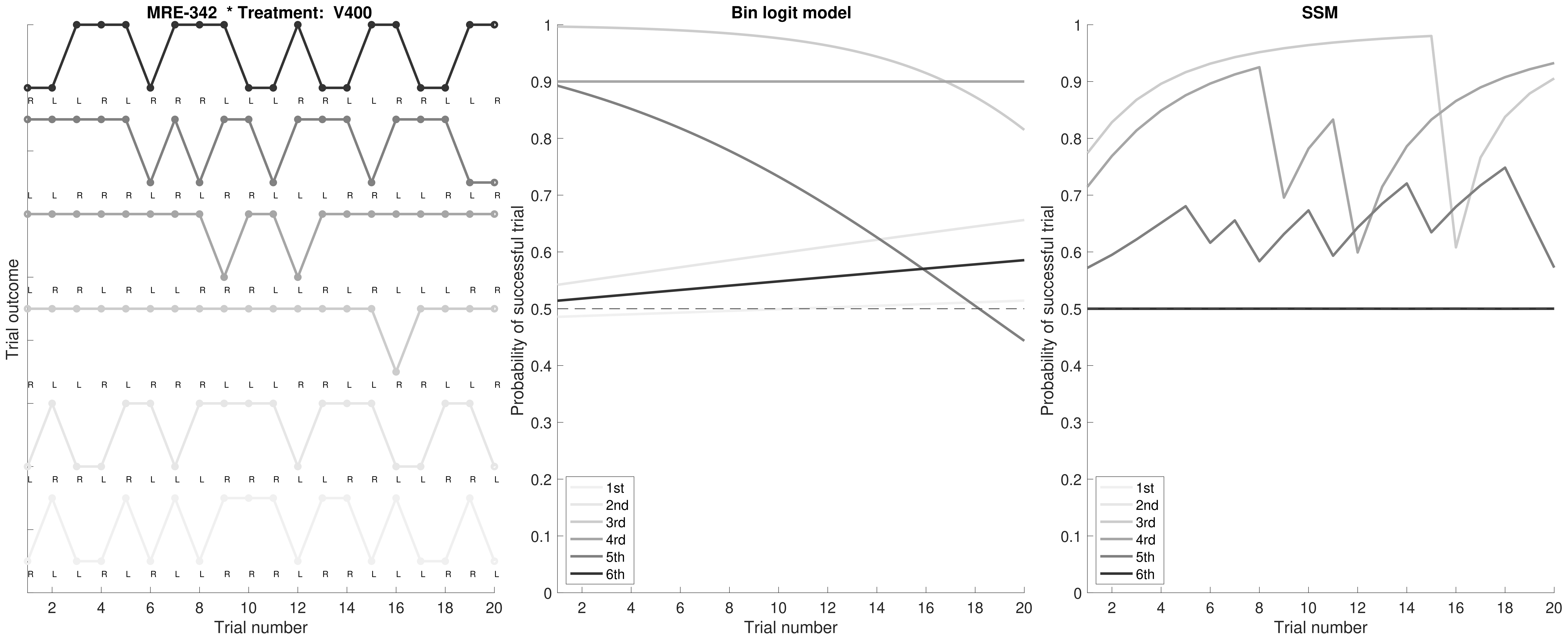

MRE-342 * Treatment: V400
Trial outcome
Trial number
Bin logit model
Probability of successful trial
Trial number
SSM
Probability of successful trial
Trial number
1st
2nd
3rd
4rd
5th
6th

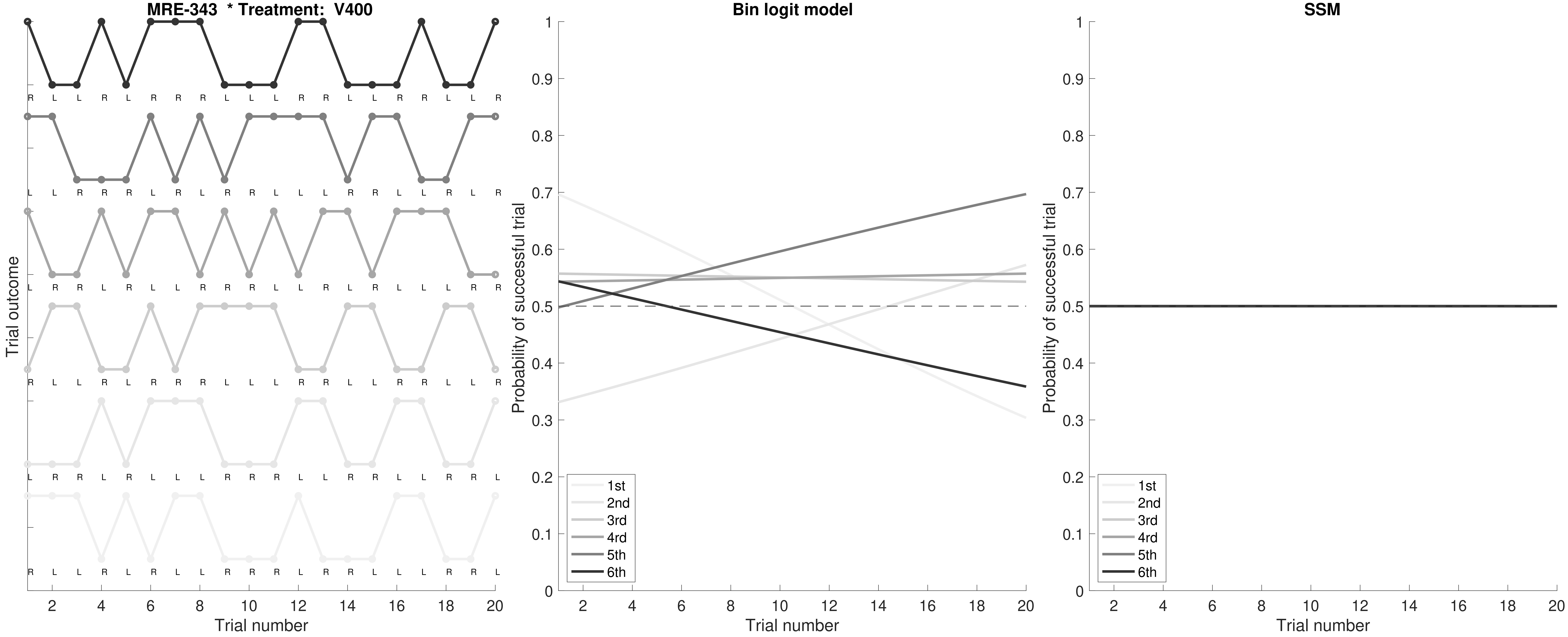

MRE-343 * Treatment: V400
Trial outcome
Trial number
Bin logit model
Probability of successful trial
Trial number
1st
2nd
3rd
4rd
5th
6th
SSM
Probability of successful trial
Trial number
1st
2nd
3rd
4rd
5th
6th

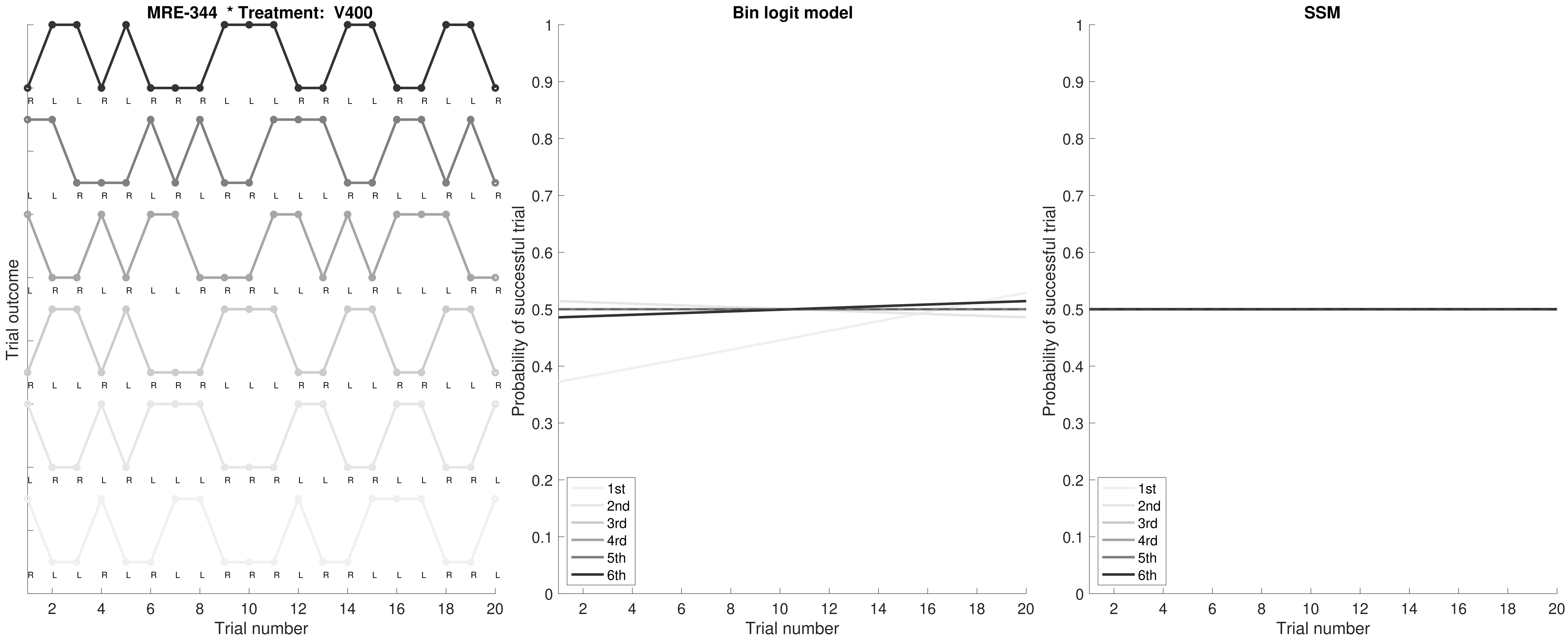

MRE-344 * Treatment: V400
Trial outcome
Trial number
Bin logit model
Probability of successful trial
Trial number
SSM
Probability of successful trial
Trial number
1st
2nd
3rd
4rd
5th
6th

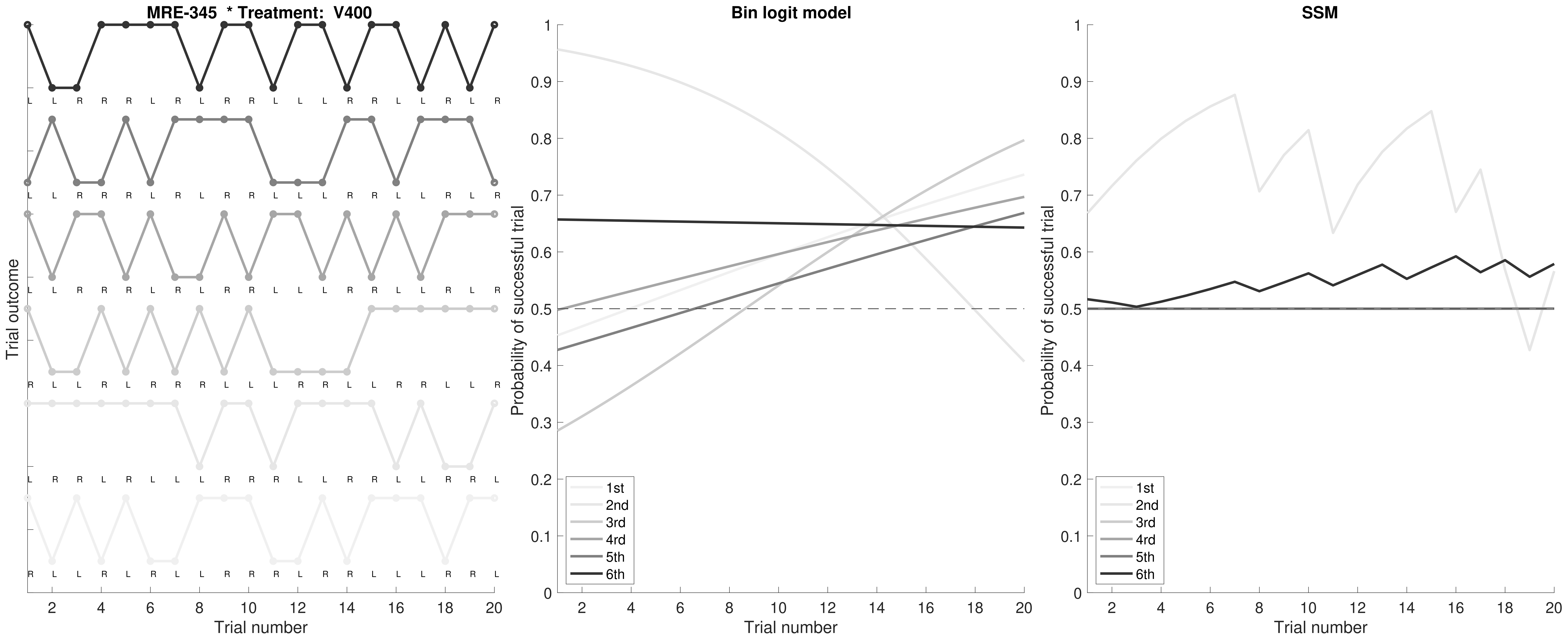
MRE-345 * Treatment: V400
Trial outcome
Trial number
Bin logit model
Probability of successful trial
Trial number
1st
2nd
3rd
4rd
5th
6th
SSM
Probability of successful trial
Trial number
1st
2nd
3rd
4rd
5th
6th

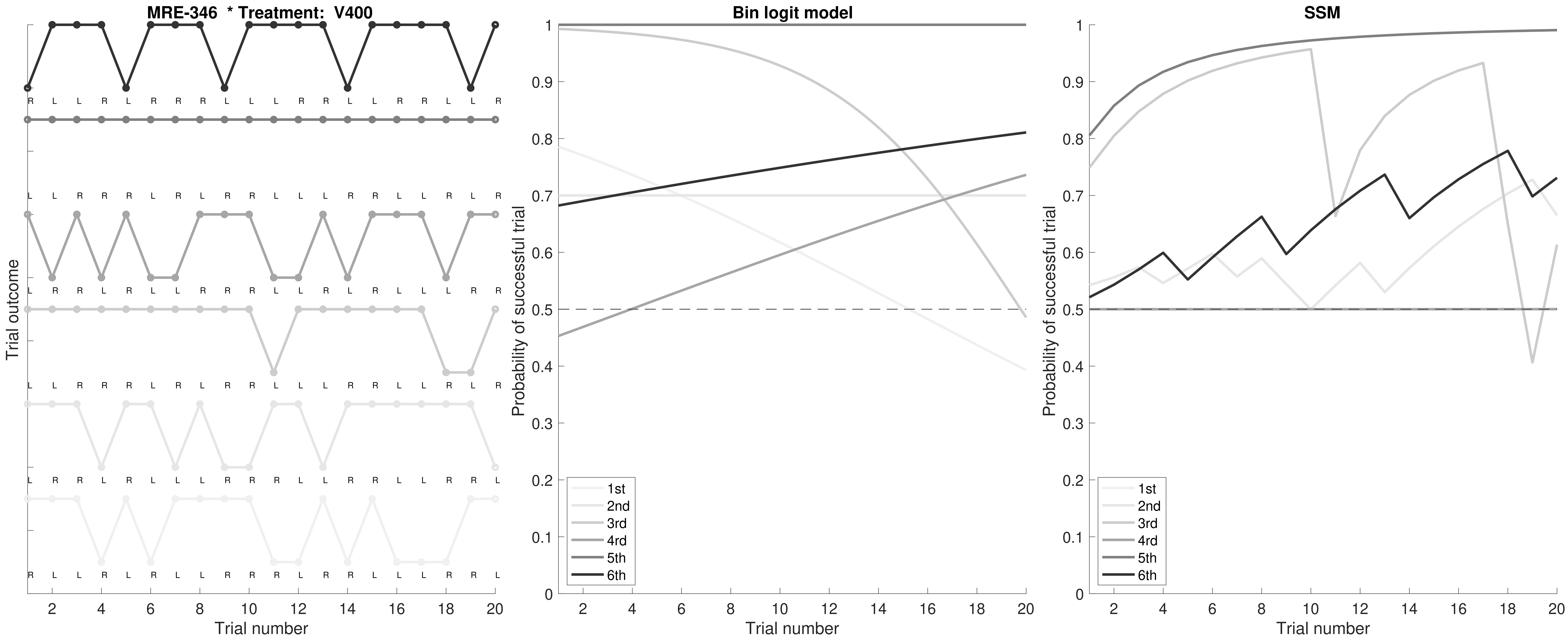
MRE-346 * Treatment: V400
Trial outcome
Trial number
Bin logit model
Probability of successful trial
Trial number
1st
2nd
3rd
4rd
5th
6th
SSM
Probability of successful trial
Trial number
1st
2nd
3rd
4rd
5th
6th

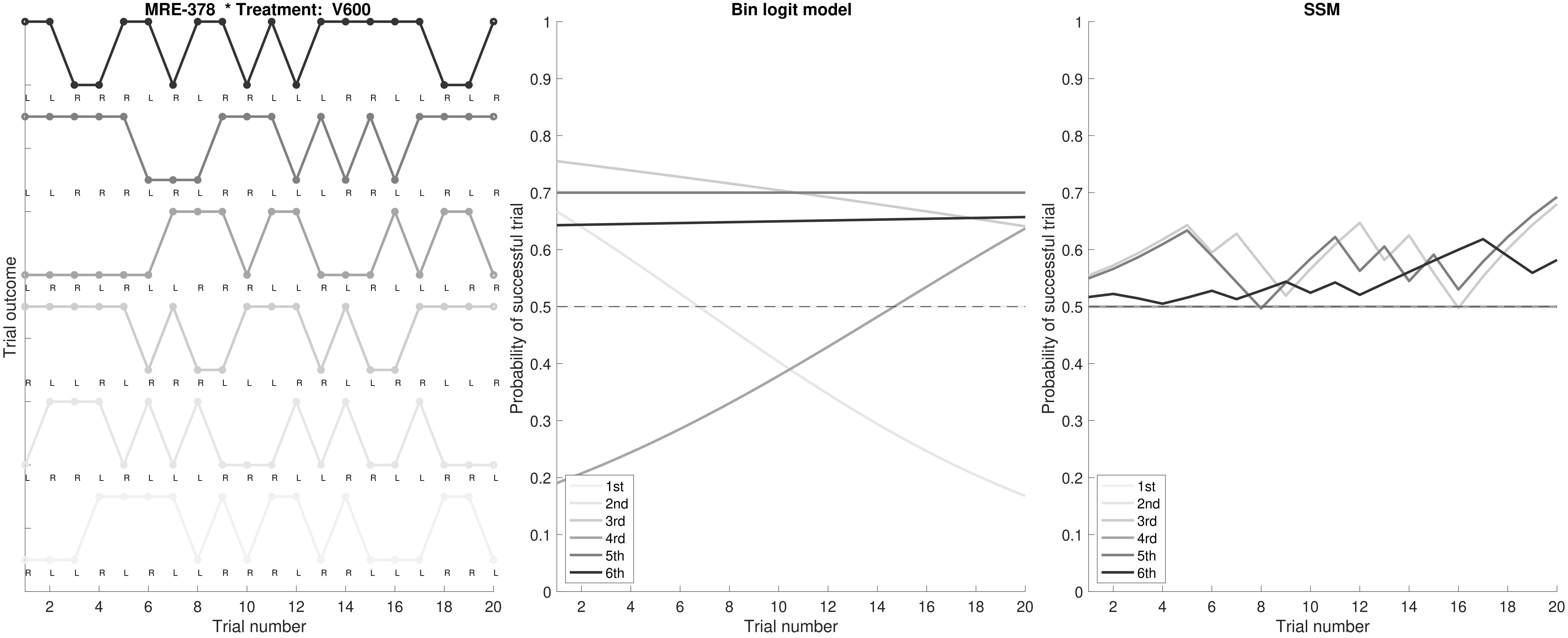

MRE-378 * Treatment: V600
Trial outcome
Trial number
Bin logit model
Probability of successful trial
Trial number
1st
2nd
3rd
4rd
5th
6th
SSM
Probability of successful trial
Trial number
1st
2nd
3rd
4rd
5th
6th

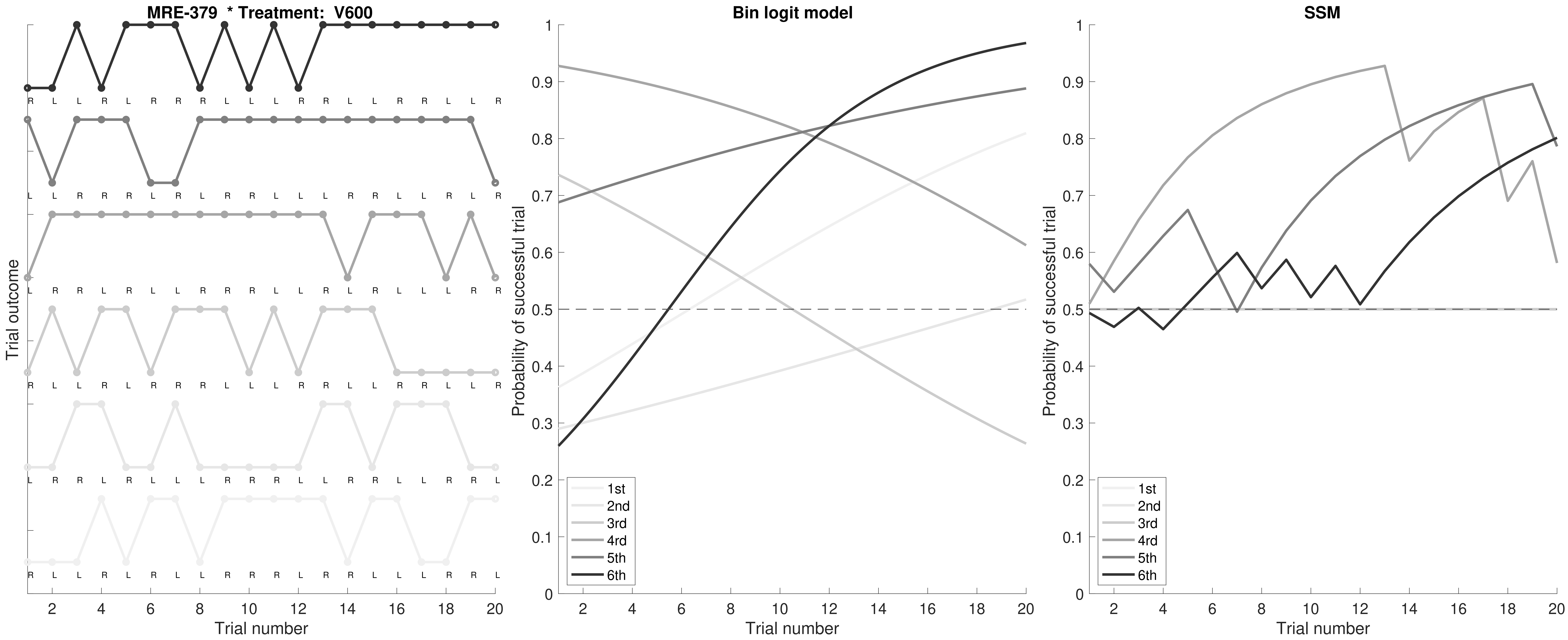
MRE-379 * Treatment: V600
R L L R L R R R L L L R R L L R R L L R
L L R R R L R L R R L L L R R L L R L R
L R R L R L L R R R L L R L R L L L R R
R L L R L R R R L L L R R L L R R L L R
L R R L R L L L R R R L L R R L L R R L
R L L R L R L L R R R L R R L L L R R L
Trial outcome
Trial number
Bin logit model
Probability of successful trial
Trial number
1st
2nd
3rd
4rd
5th
6th
SSM
Probability of successful trial
Trial number
1st
2nd
3rd
4rd
5th
6th

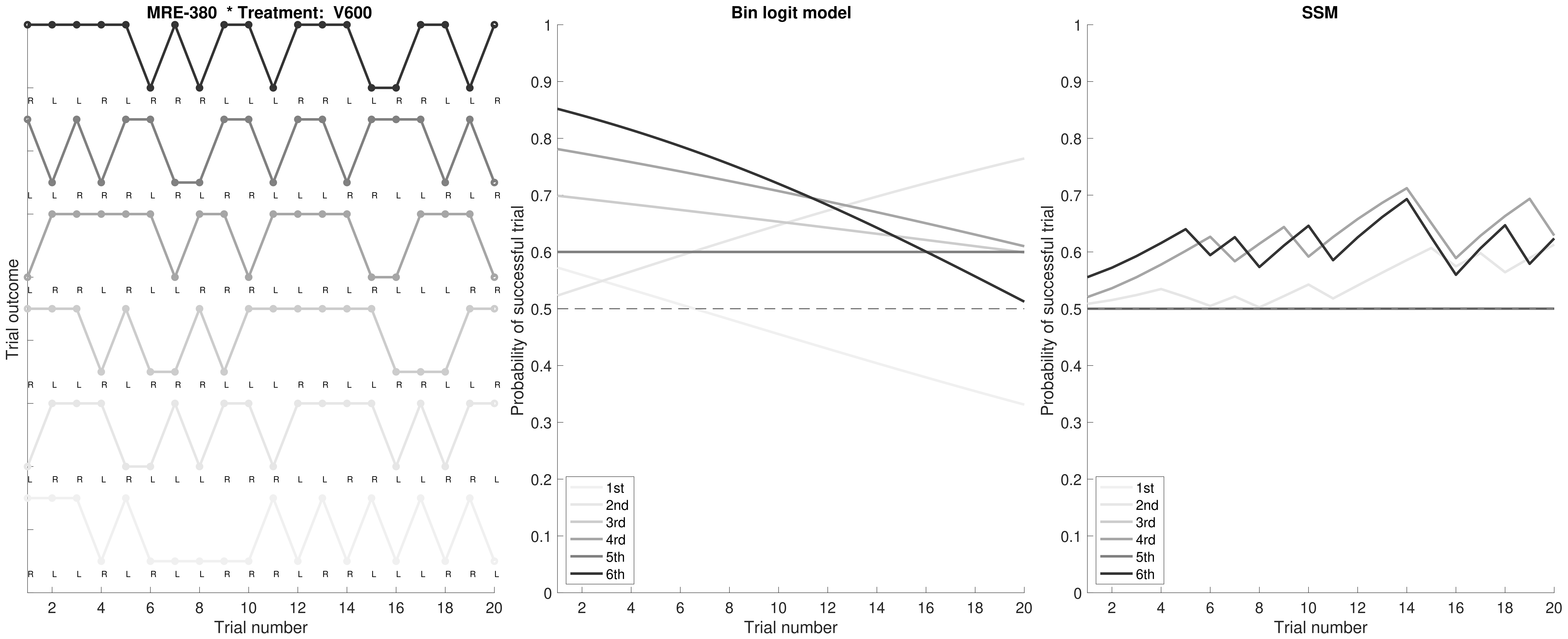

MRE-380 * Treatment: V600
Trial outcome
Trial number
R L L R L R R R L L L R R L L R R L L R
L L R R R L R L R R L L L R R L L R L R
L R R L R L L R R L R L R R L L L L R R
R L L R L R R R L L L R R L L R R L L R
L R R L R L L L R R R L L R R L L R R L
R L L R L R L L R R R L R R L L L R R L
Bin logit model
Probability of successful trial
Trial number
1st
2nd
3rd
4rd
5th
6th
SSM
Probability of successful trial
Trial number
1st
2nd
3rd
4rd
5th
6th

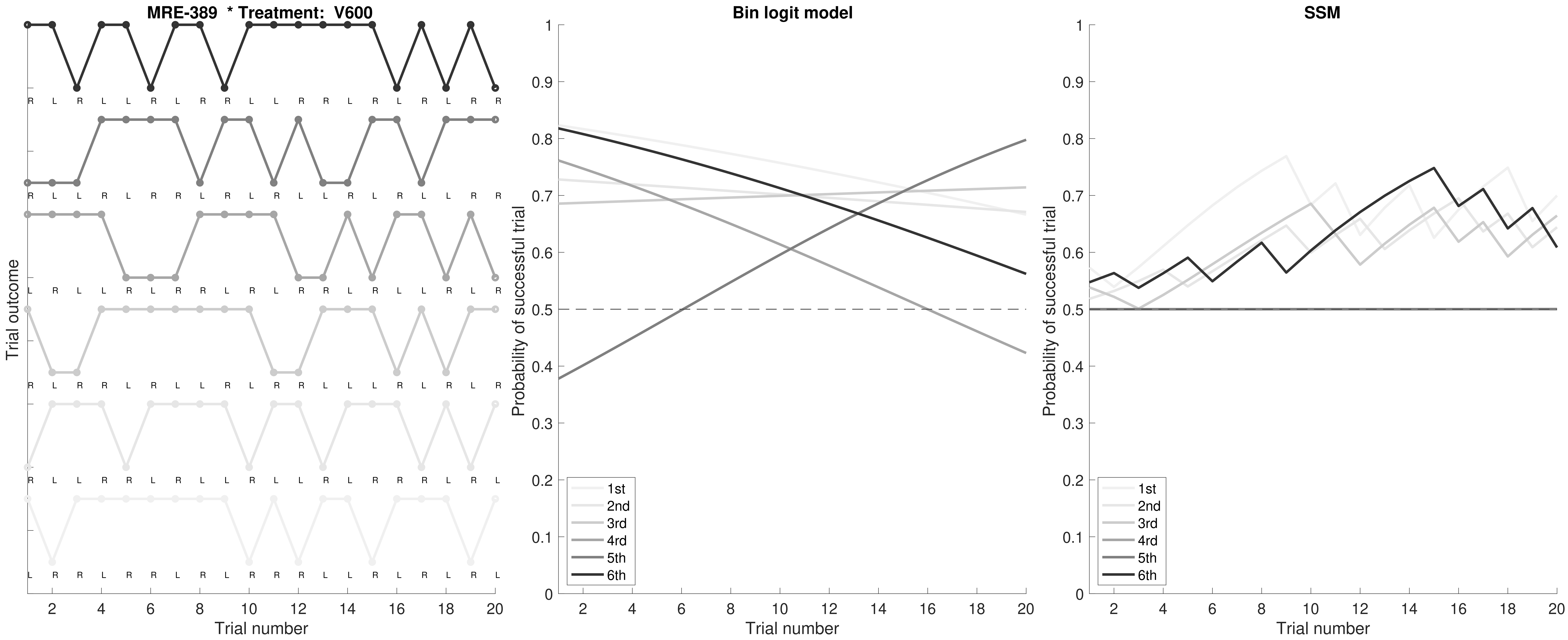
MRE-389 * Treatment: V600
Trial outcome
R L R L L R L R R L L R R L R L R L R R
R L L R L R R L R R R L L R L R L L R R
L R L L R L R R L R L R R L R L L R L R
R R R R L R L L R L R R L L L R L R R R
R L L R R L R L L R L L R L R R R L R L
L R R L R R L R L R R R L L R L R L R L
2 4 6 8 10 12 14 16 18 20
Trial number
Bin logit model
Probability of successful trial
0 0.1 0.2 0.3 0.4 0.5 0.6 0.7 0.8 0.9 1
1st
2nd
3rd
4rd
5th
6th
2 4 6 8 10 12 14 16 18 20
Trial number
SSM
Probability of successful trial
0 0.1 0.2 0.3 0.4 0.5 0.6 0.7 0.8 0.9 1
1st
2nd
3rd
4rd
5th
6th
2 4 6 8 10 12 14 16 18 20
Trial number

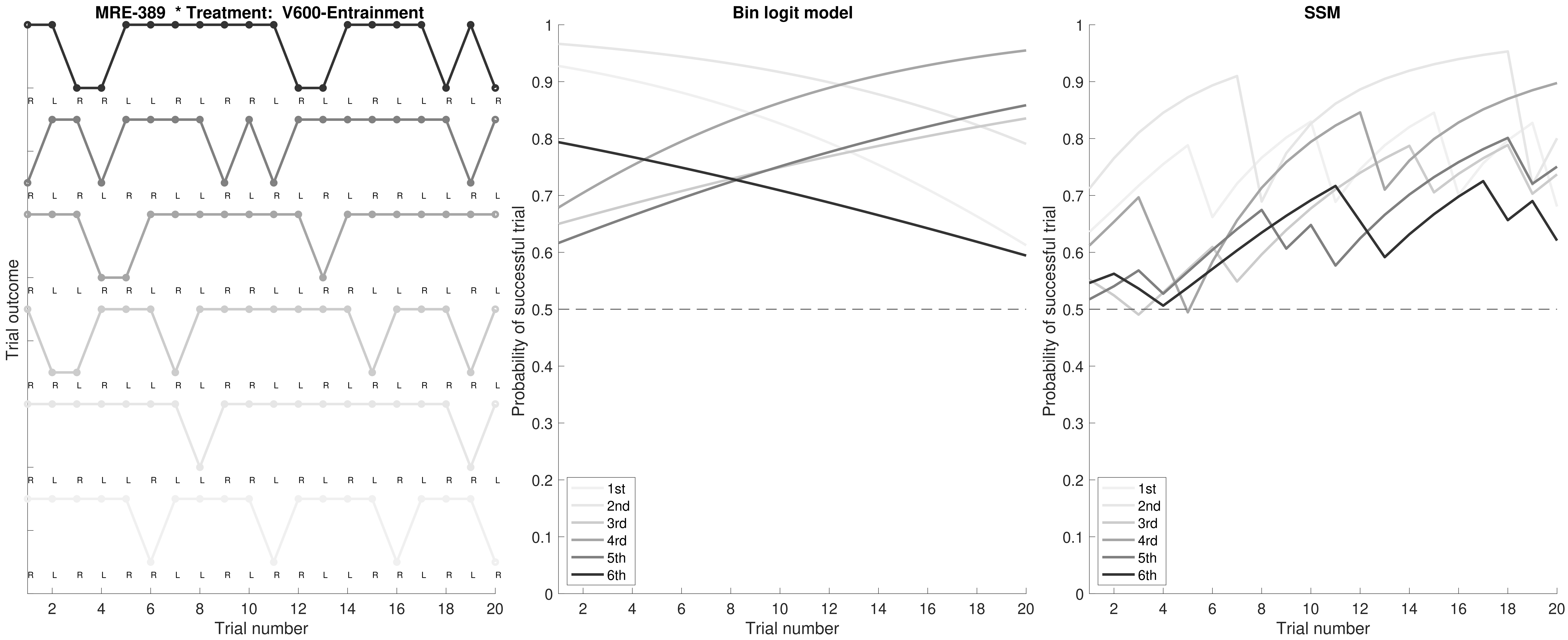

MRE-389 * Treatment: V600-Entrainment
Trial outcome
Trial number
Bin logit model
Probability of successful trial
Trial number
1st
2nd
3rd
4rd
5th
6th
SSM
Probability of successful trial
Trial number
1st
2nd
3rd
4rd
5th
6th

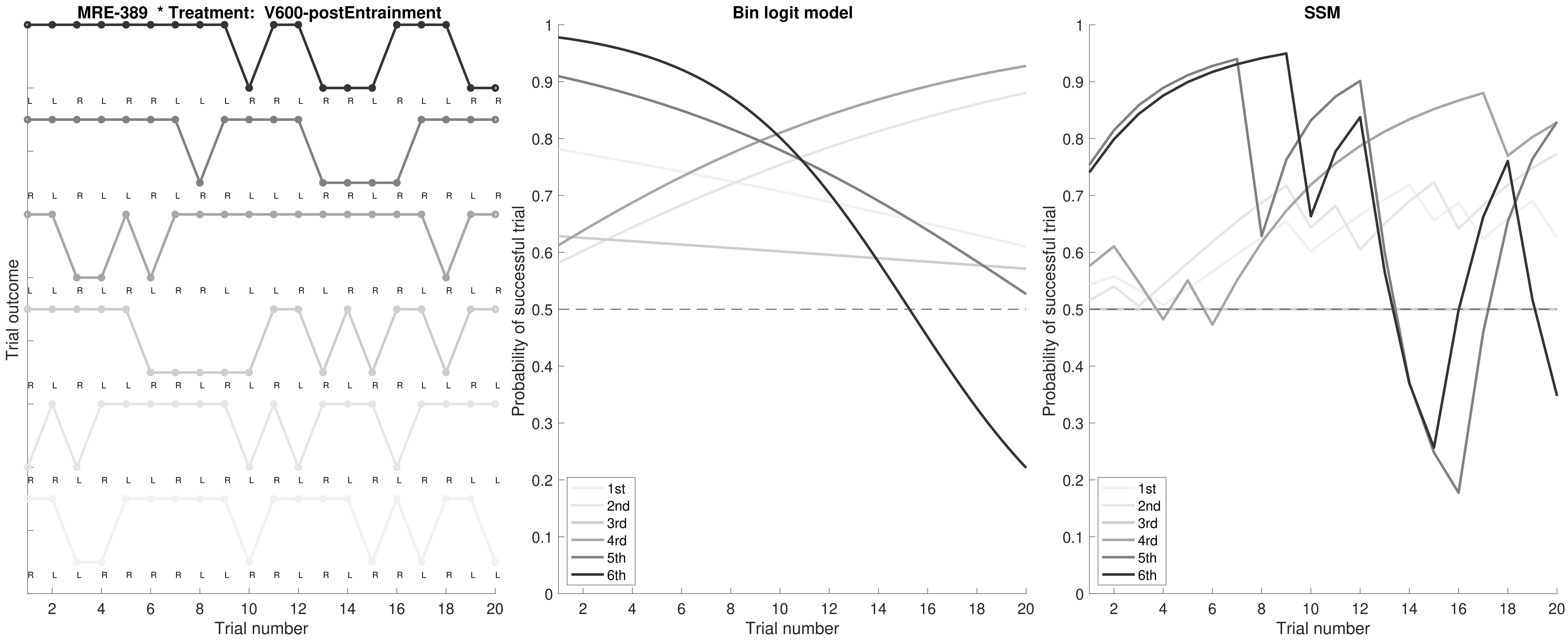
MRE-389 * Treatment: V600-postEntrainment
Trial outcome
Trial number
Bin logit model
Probability of successful trial
Trial number
1st
2nd
3rd
4rd
5th
6th
SSM
Probability of successful trial
Trial number
1st
2nd
3rd
4rd
5th
6th

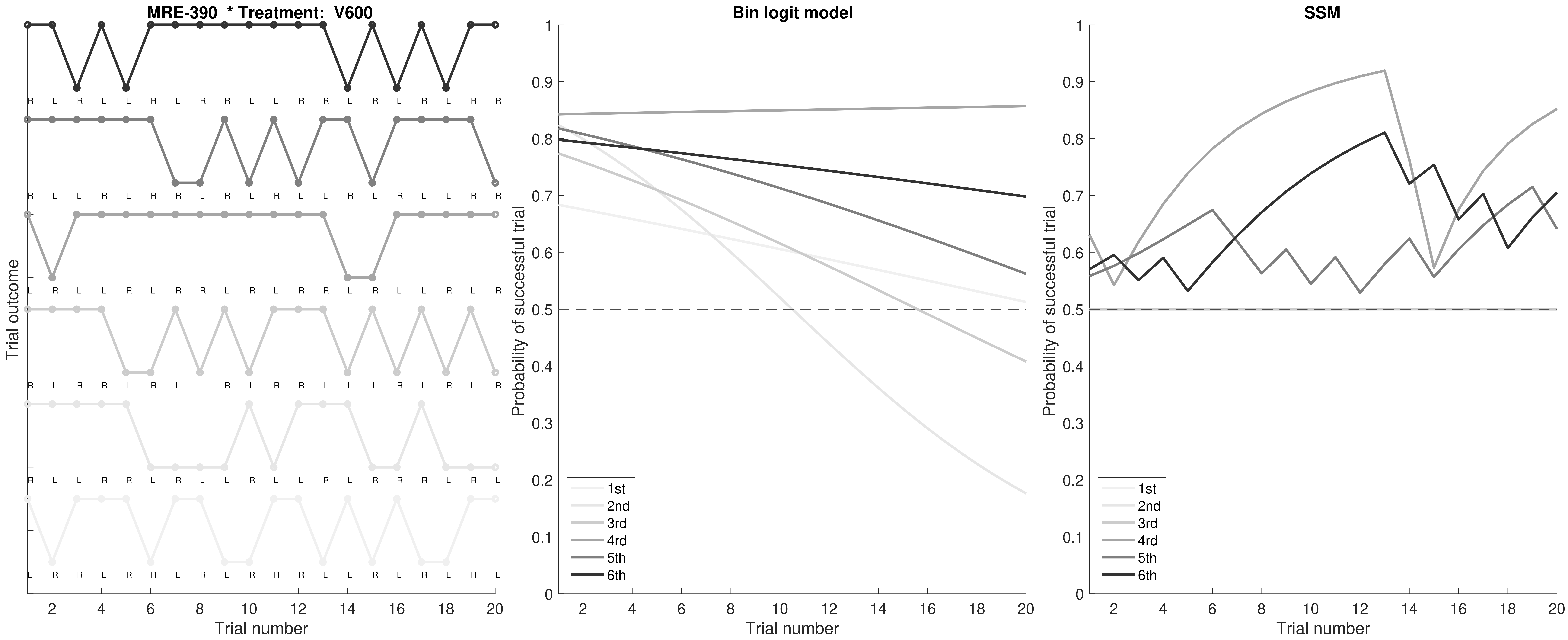

MRE-390 * Treatment: V600
Trial outcome
Trial number
Bin logit model
Probability of successful trial
Trial number
SSM
Probability of successful trial
Trial number
1st
2nd
3rd
4rd
5th
6th

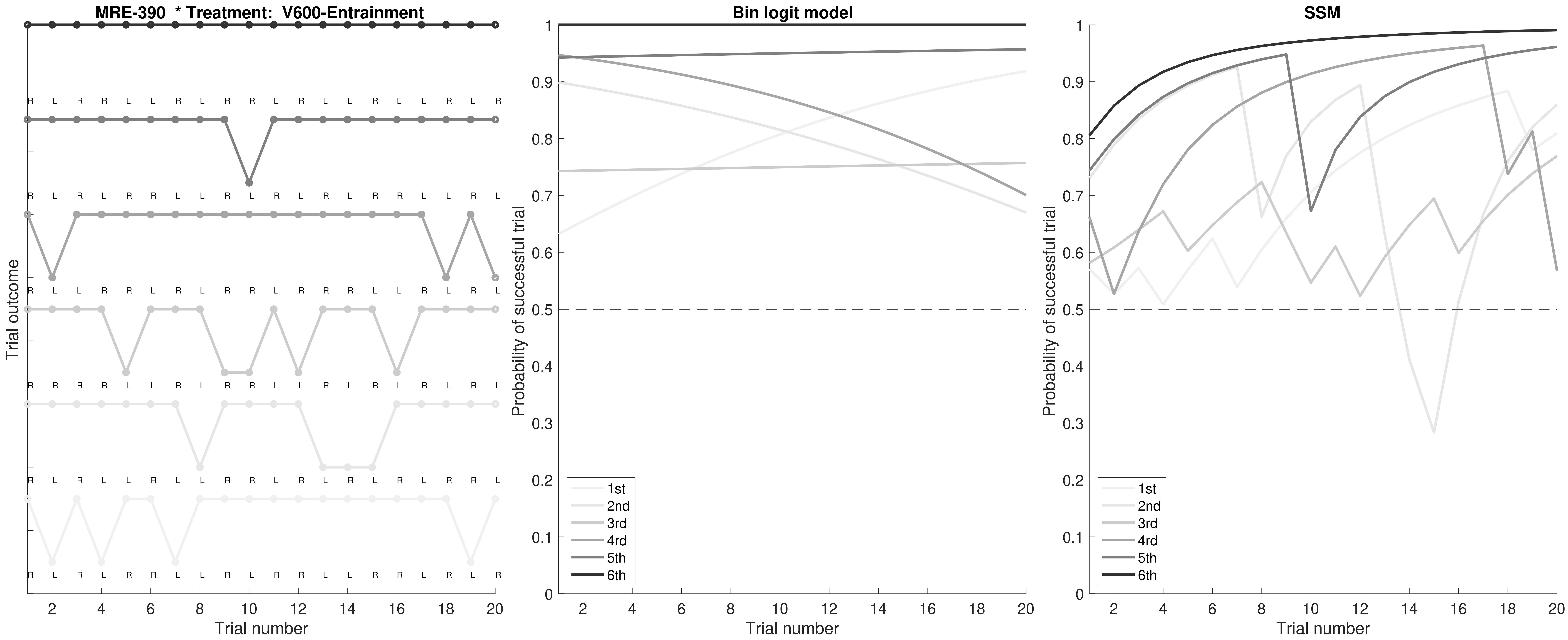
MRE-390 * Treatment: V600-Entrainment
Trial outcome
Trial number
Bin logit model
Probability of successful trial
Trial number
SSM
Probability of successful trial
Trial number
1st
2nd
3rd
4rd
5th
6th

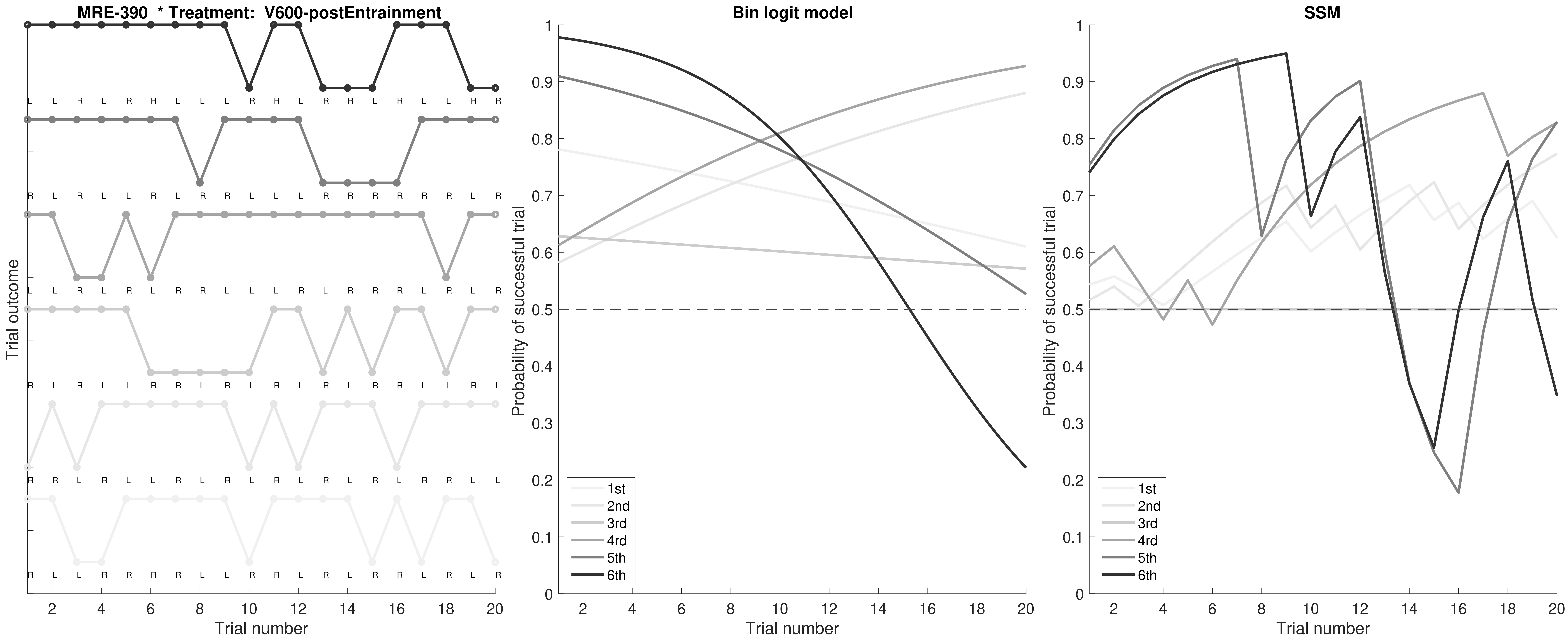
MRE-390 * Treatment: V600-postEntrainment
Trial outcome
Trial number
Bin logit model
Probability of successful trial
Trial number
1st
2nd
3rd
4rd
5th
6th
SSM
Probability of successful trial
Trial number
1st
2nd
3rd
4rd
5th
6th

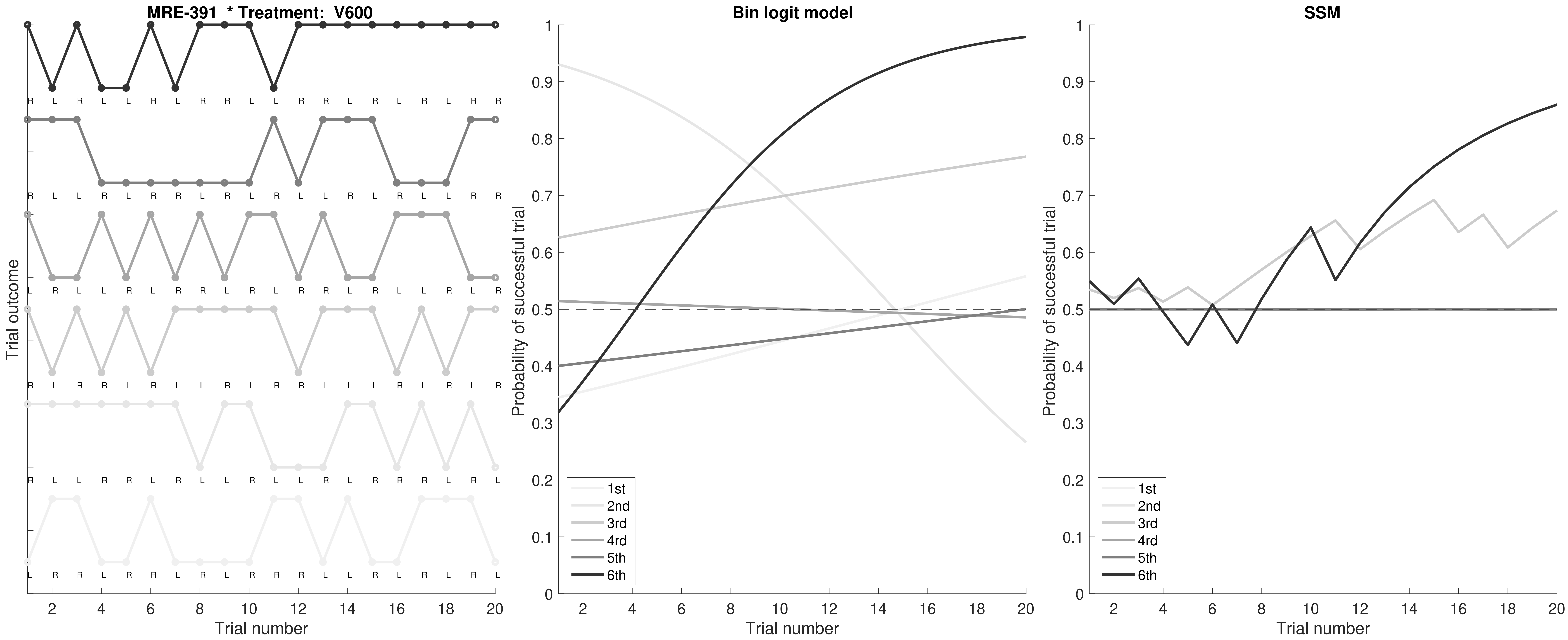

MRE-391 * Treatment: V600
Trial outcome
Trial number
Bin logit model
Probability of successful trial
Trial number
1st
2nd
3rd
4rd
5th
6th
SSM
Probability of successful trial
Trial number
1st
2nd
3rd
4rd
5th
6th

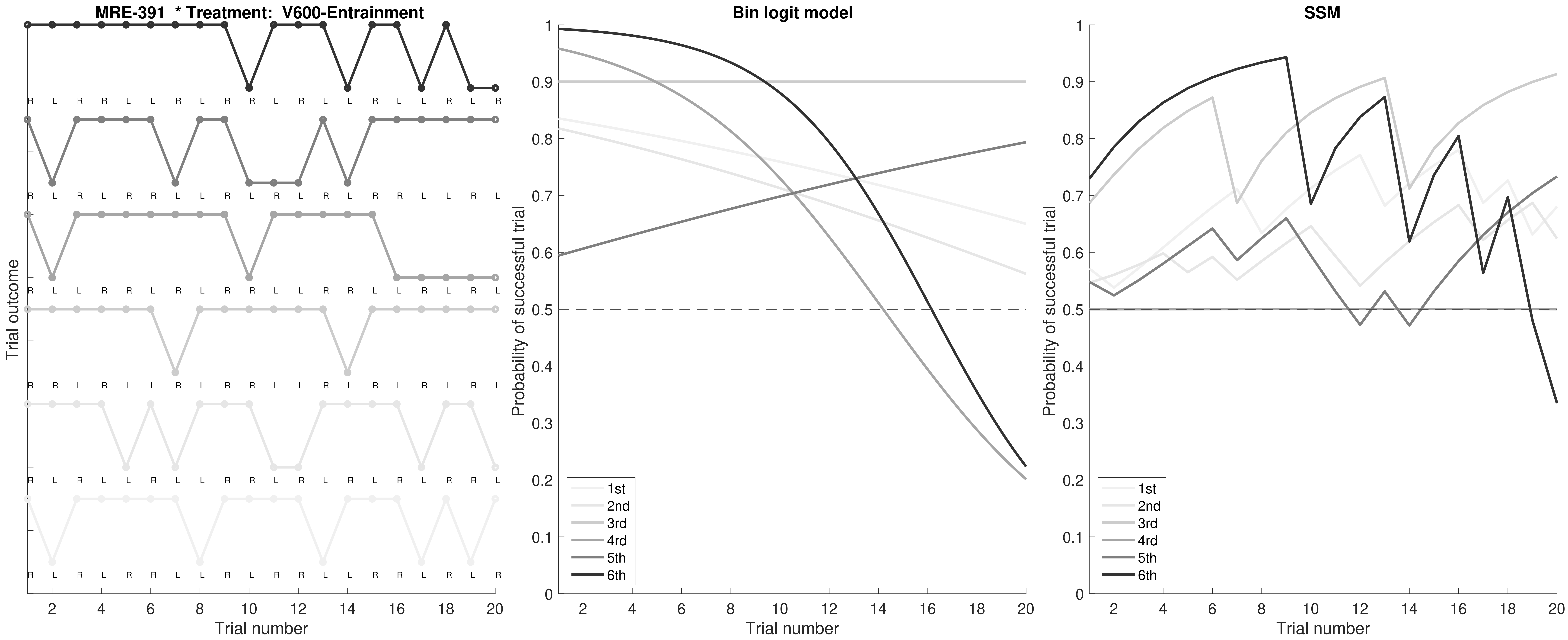

MRE-391 * Treatment: V600-Entrainment
Trial outcome
Trial number
Bin logit model
Probability of successful trial
Trial number
1st
2nd
3rd
4rd
5th
6th
SSM
Probability of successful trial
Trial number
1st
2nd
3rd
4rd
5th
6th

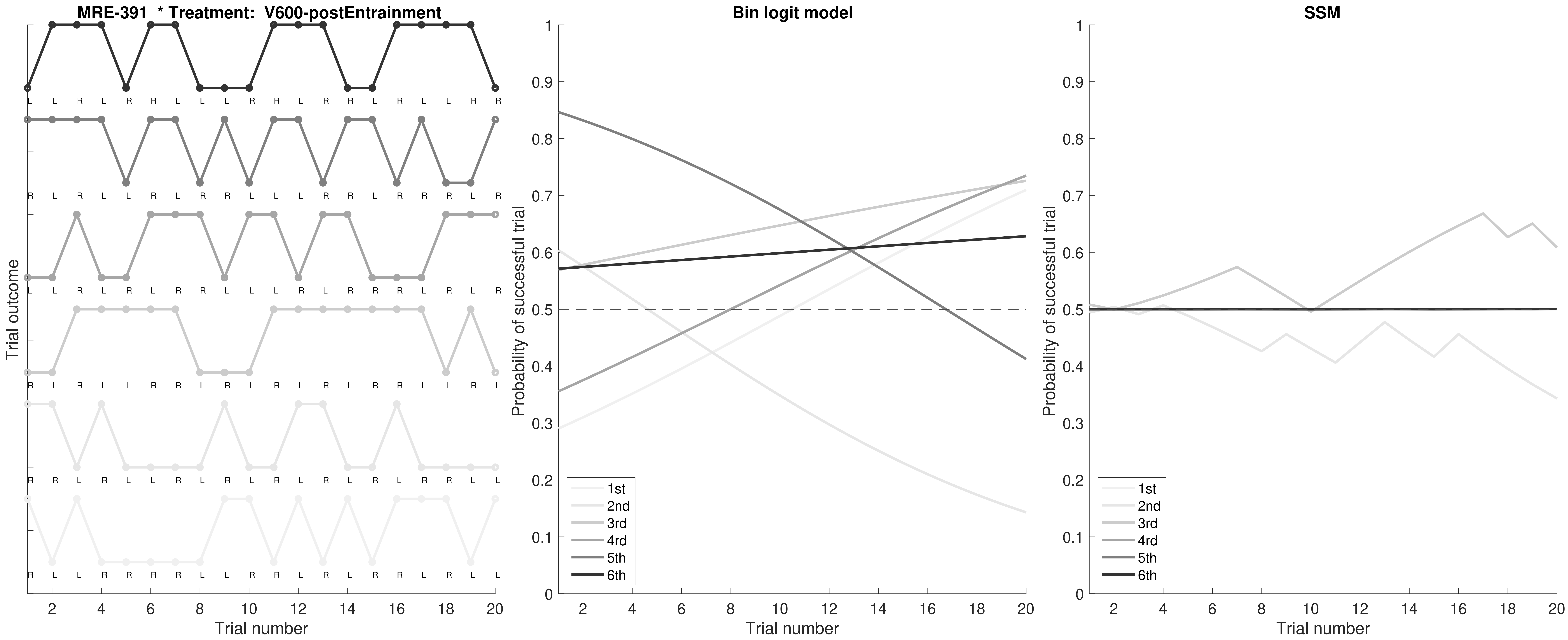

MRE-391 * Treatment: V600-postEntrainment
Trial outcome
Trial number
Bin logit model
Probability of successful trial
Trial number
1st
2nd
3rd
4rd
5th
6th
SSM
Probability of successful trial
Trial number
1st
2nd
3rd
4rd
5th
6th

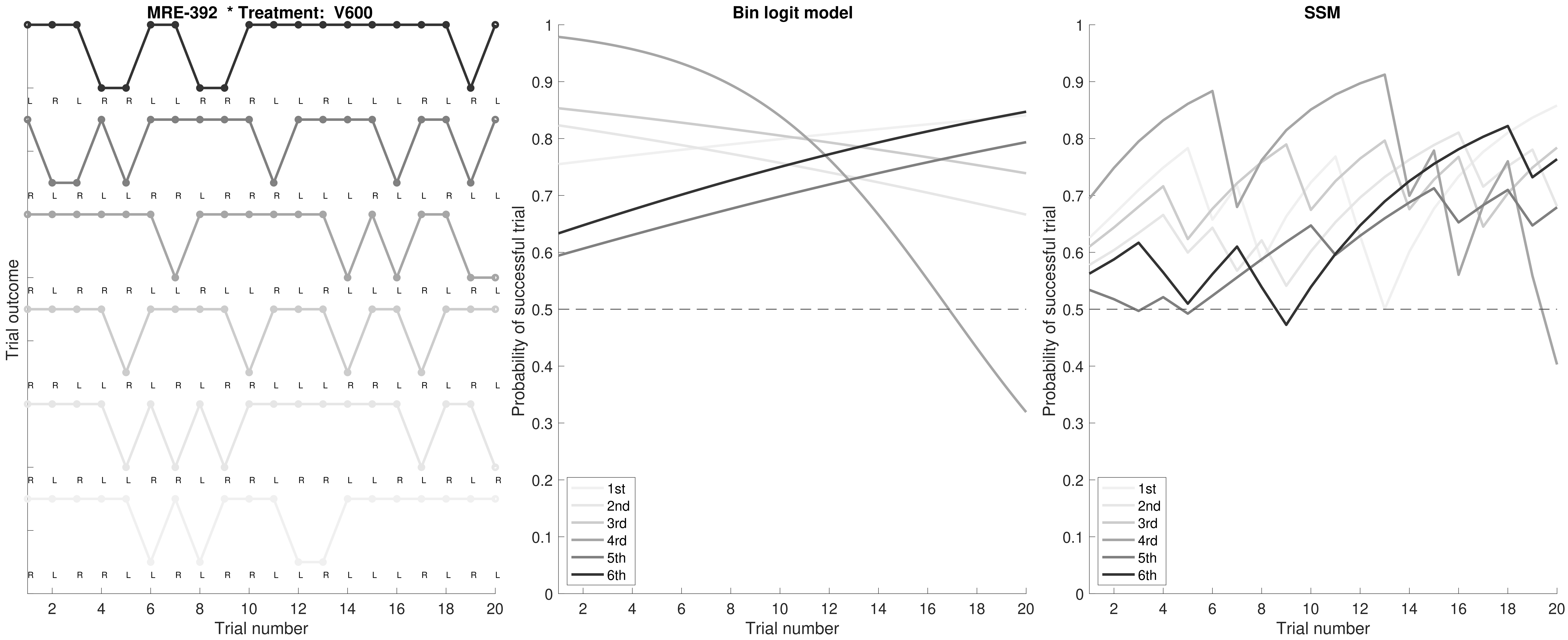

MRE-392 * Treatment: V600
Trial outcome
Trial number
Bin logit model
Probability of successful trial
SSM
1st
2nd
3rd
4rd
5th
6th

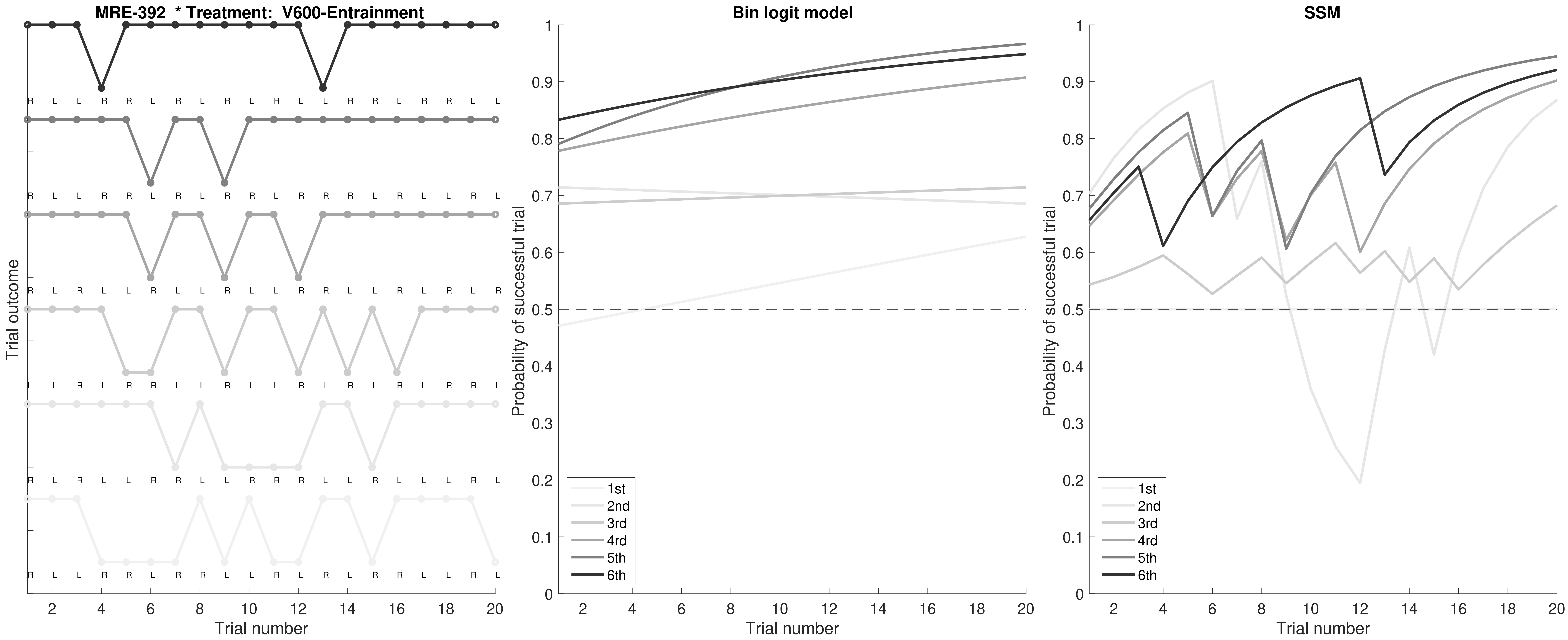

MRE-392 * Treatment: V600-Entrainment
Trial outcome
Trial number
Bin logit model
Probability of successful trial
Trial number
1st
2nd
3rd
4rd
5th
6th
SSM
Probability of successful trial
Trial number
1st
2nd
3rd
4rd
5th
6th

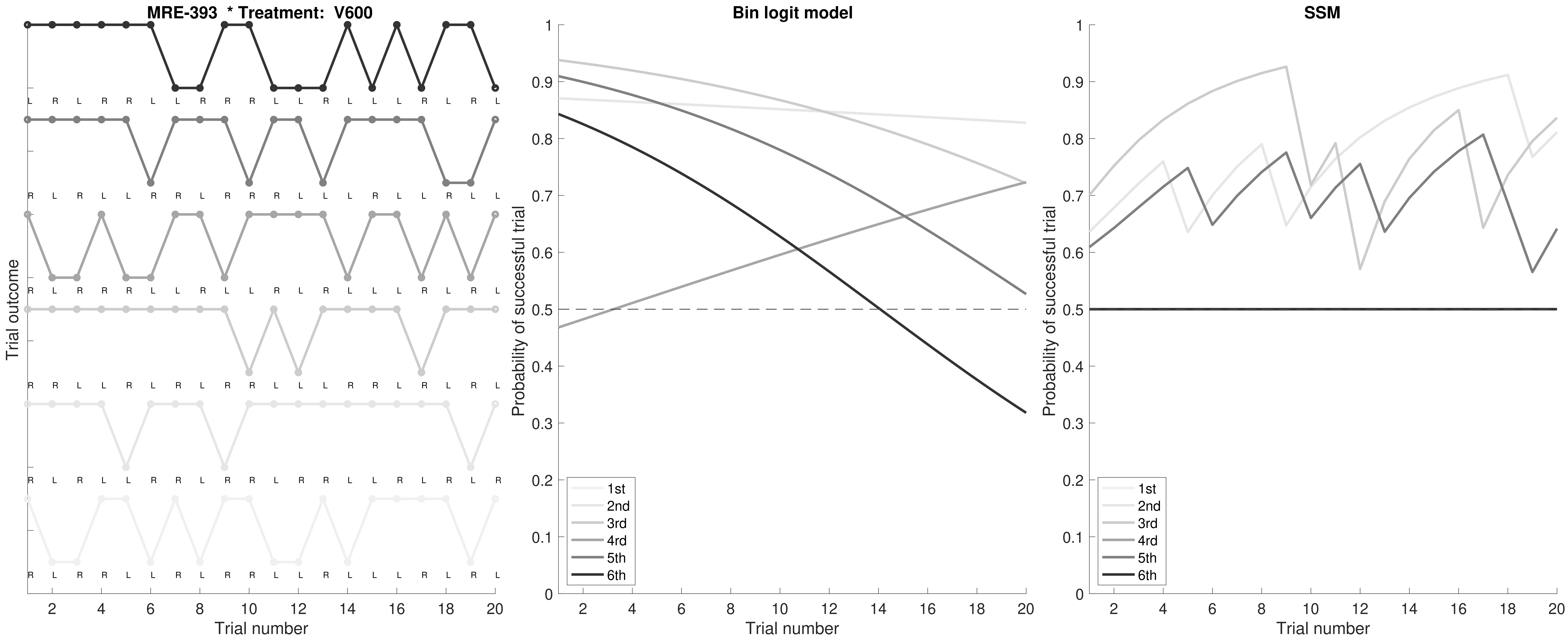
MRE-393 * Treatment: V600
Trial outcome
Trial number
Bin logit model
Probability of successful trial
Trial number
1st
2nd
3rd
4rd
5th
6th
SSM
Probability of successful trial
Trial number
1st
2nd
3rd
4rd
5th
6th

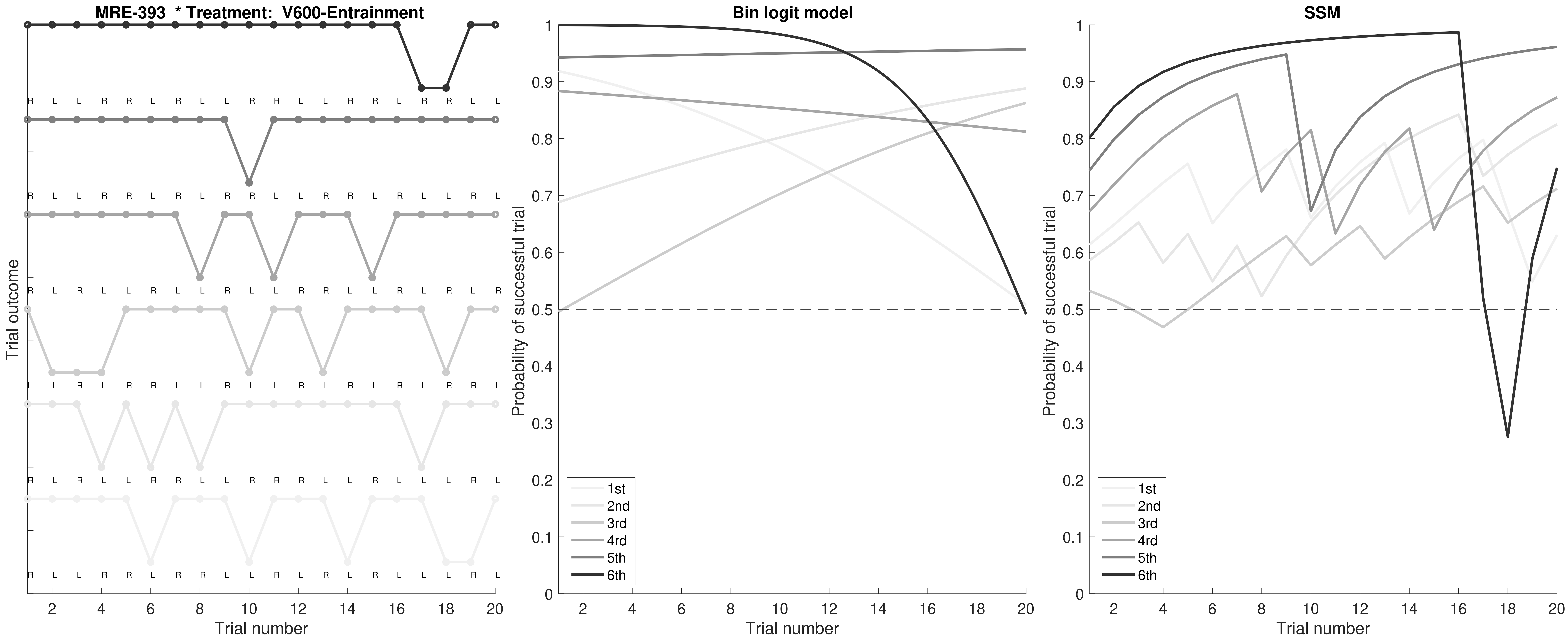

MRE-393 * Treatment: V600-Entrainment
Trial outcome
Trial number
Bin logit model
Probability of successful trial
SSM
1st
2nd
3rd
4rd
5th
6th

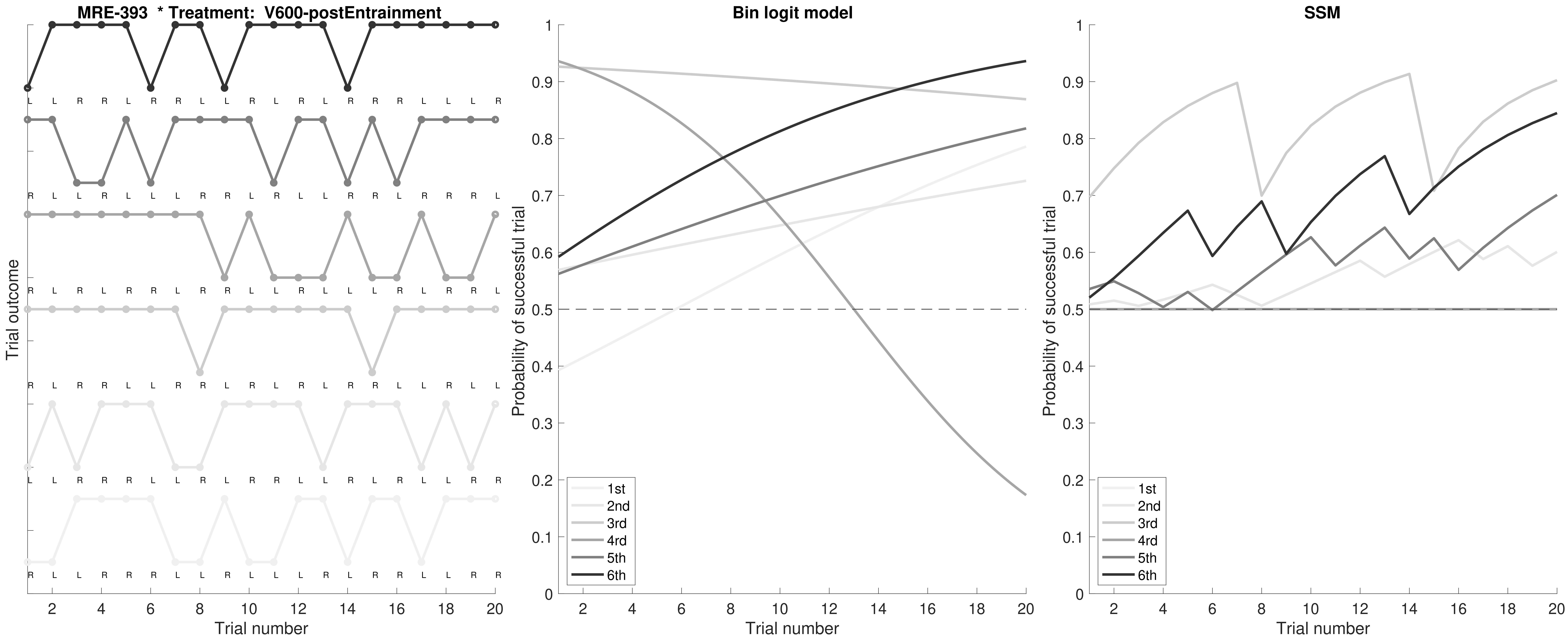
MRE-393 * Treatment: V600-postEntrainment
Trial outcome
Trial number
Bin logit model
Probability of successful trial
SSM
1st
2nd
3rd
4rd
5th
6th

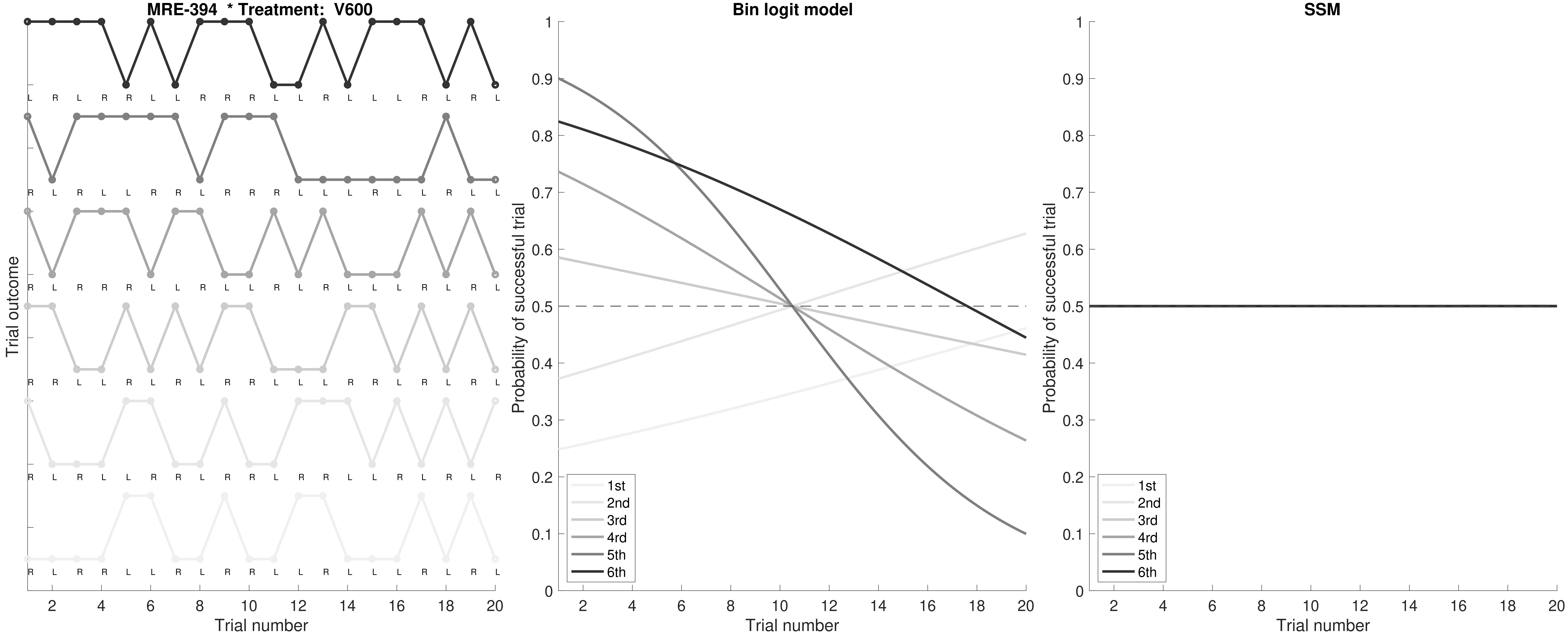

MRE-394 * Treatment: V600
L R L R R L L R R R L L R L L L R L R L
R L R L L R R L R R R L L L R L L R L L
R L R R R L L R L L R L R L L L R L R L
R R L L R L R L R R L L L R R L R L R L
R L R L L R R L R L L R R L L R L R L R
R L R R L L R L R R L L R L L L R L R L
Trial outcome
Trial number
Bin logit model
Probability of successful trial
Trial number
1st
2nd
3rd
4rd
5th
6th
SSM
Probability of successful trial
Trial number
1st
2nd
3rd
4rd
5th
6th

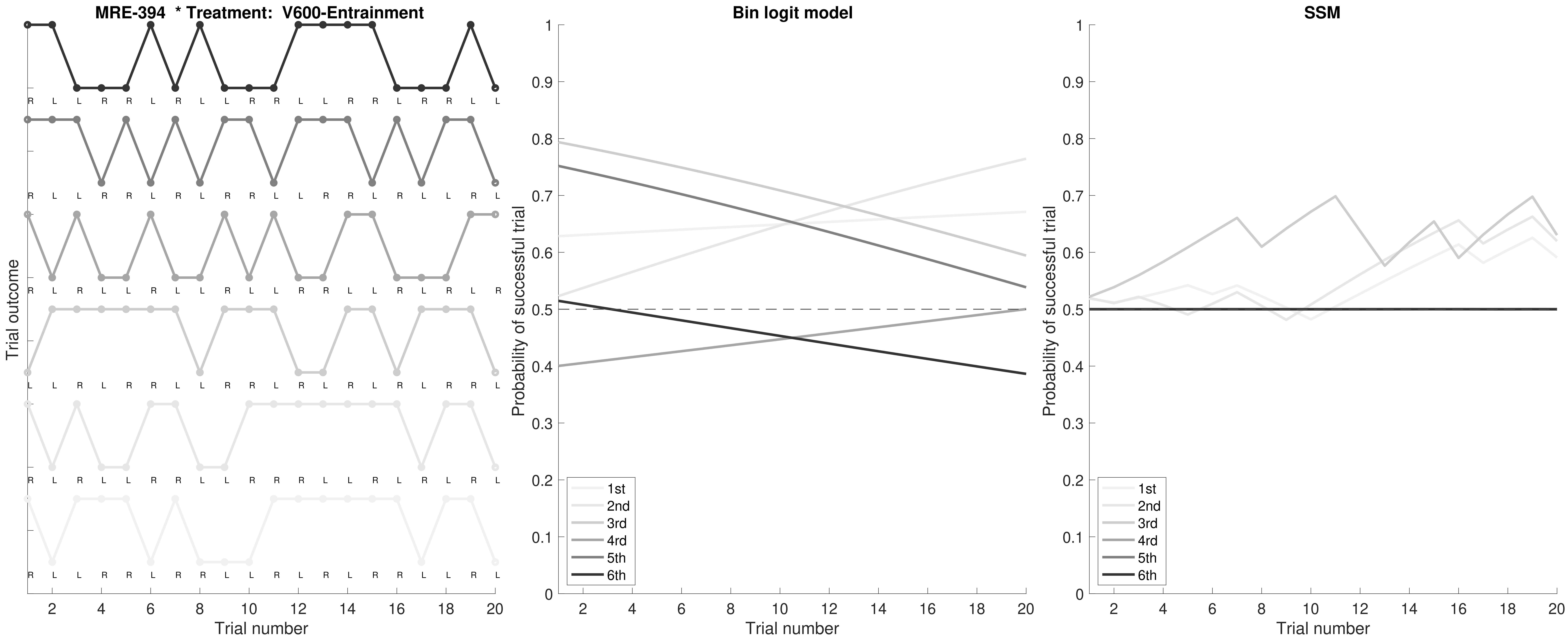

MRE-394 * Treatment: V600-Entrainment
Trial outcome
Trial number
Bin logit model
Probability of successful trial
SSM
1st
2nd
3rd
4rd
5th
6th

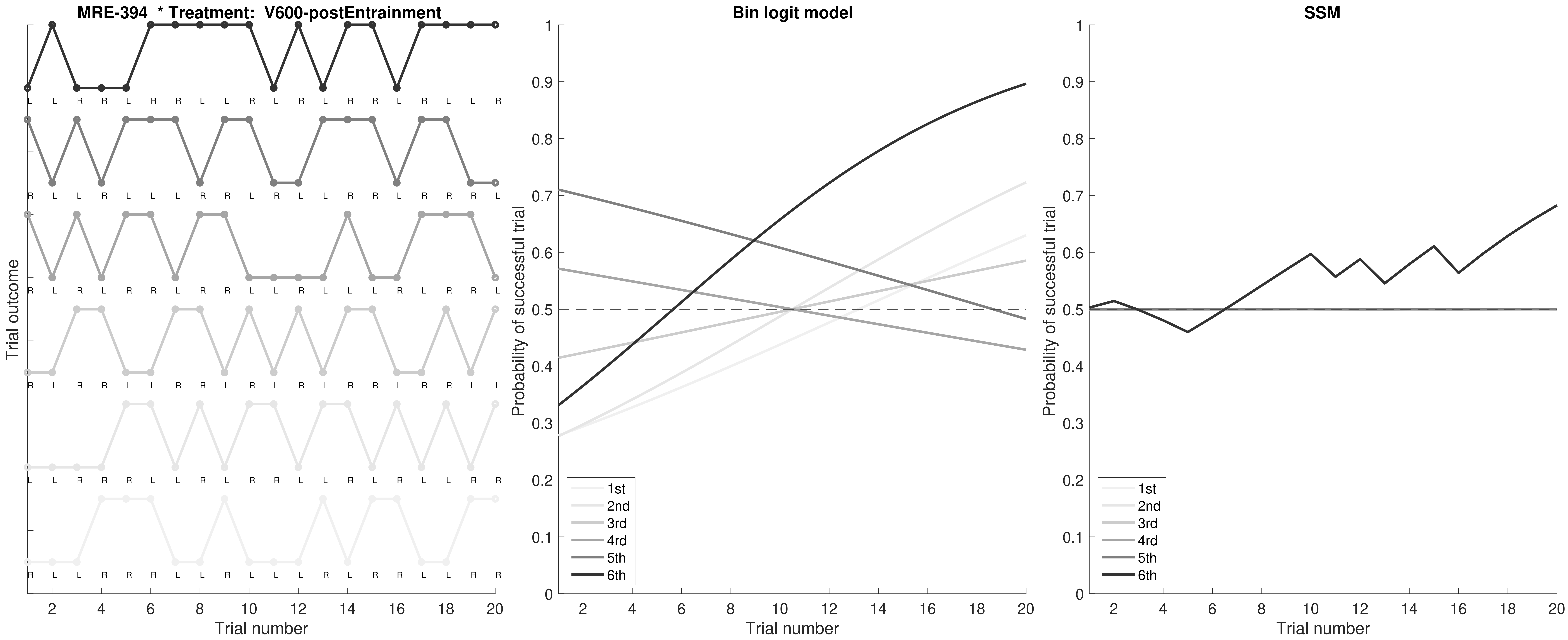
MRE-394 * Treatment: V600-postEntrainment
Trial outcome
Trial number
Bin logit model
Probability of successful trial
Trial number
SSM
Probability of successful trial
Trial number
1st
2nd
3rd
4rd
5th
6th

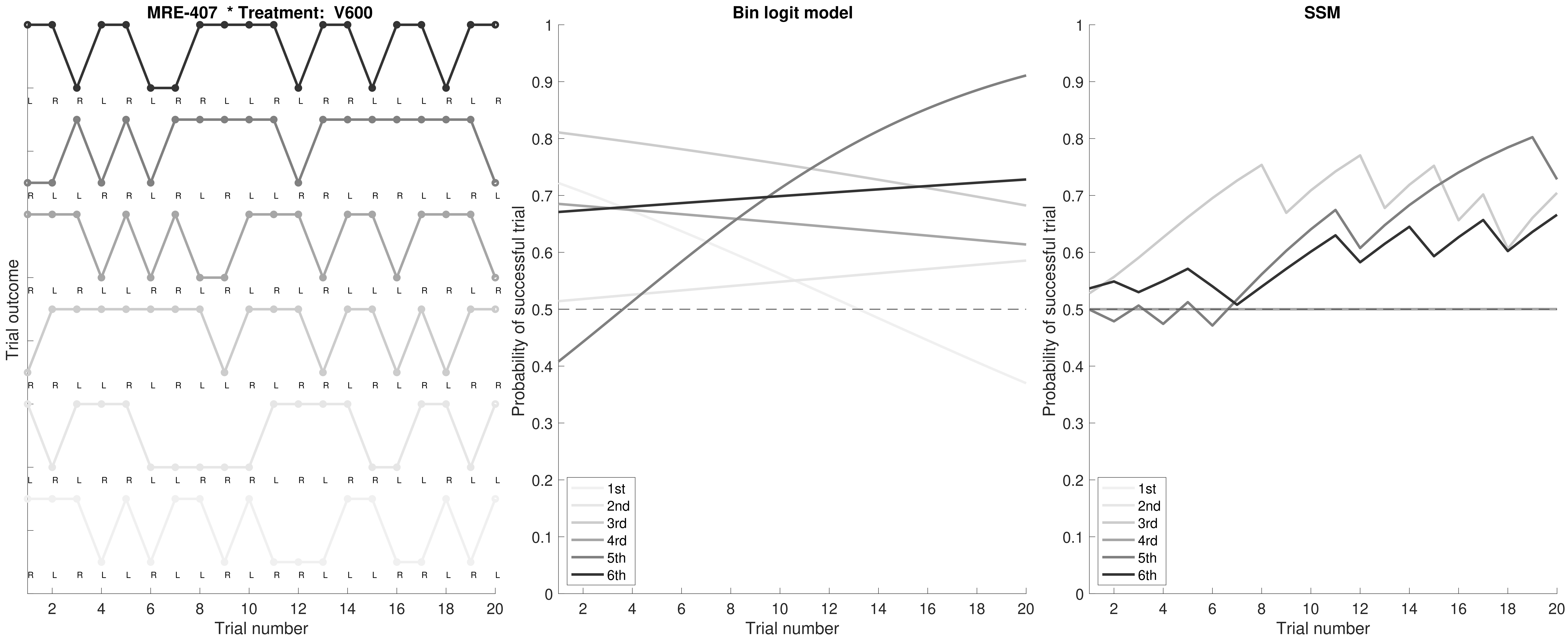

MRE-407 * Treatment: V600
Trial outcome
Trial number
Bin logit model
Probability of successful trial
Trial number
SSM
Probability of successful trial
Trial number
1st
2nd
3rd
4rd
5th
6th

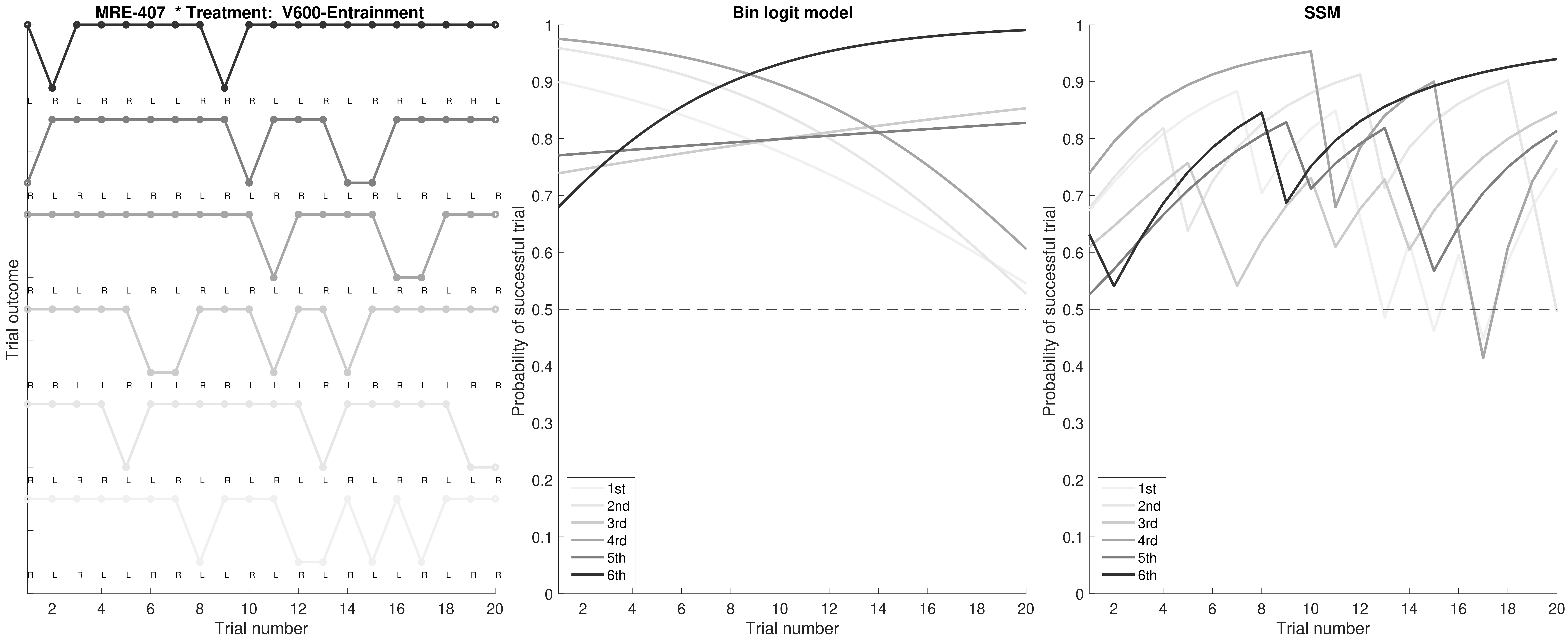

MRE-407 * Treatment: V600-Entrainment
Trial outcome
Trial number
Bin logit model
Probability of successful trial
Trial number
SSM
Probability of successful trial
Trial number
1st
2nd
3rd
4rd
5th
6th

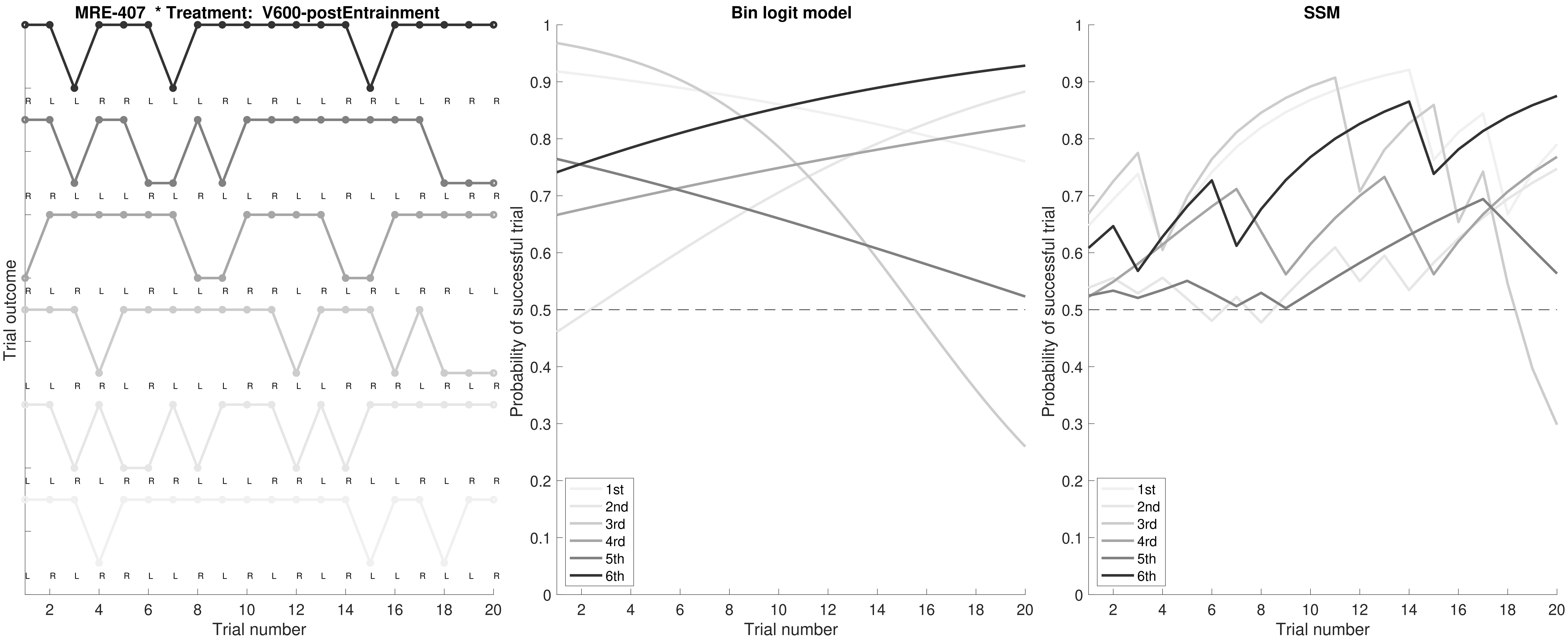

MRE-407 * Treatment: V600-postEntrainment
Trial outcome
Trial number
Bin logit model
Probability of successful trial
SSM
1st
2nd
3rd
4rd
5th
6th

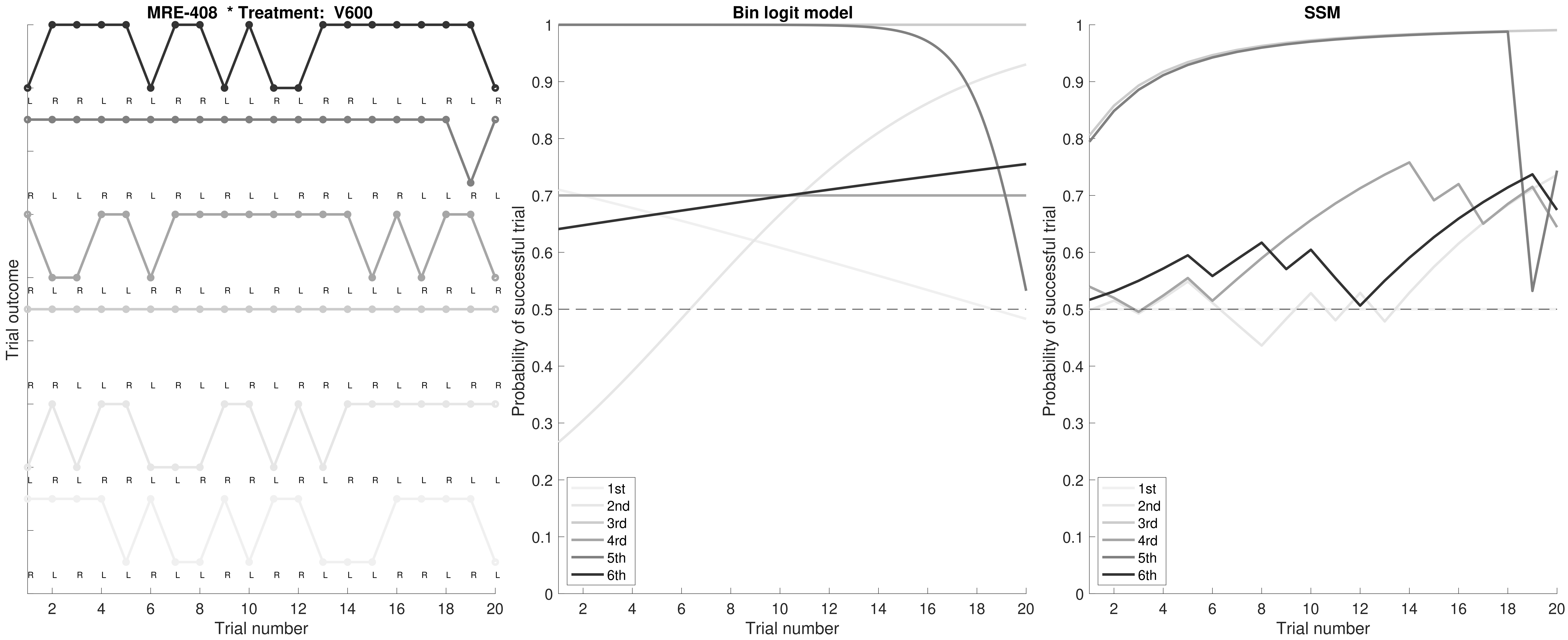

MRE-408 * Treatment: V600
Trial outcome
Trial number
Bin logit model
Probability of successful trial
Trial number
SSM
Probability of successful trial
Trial number
1st
2nd
3rd
4rd
5th
6th

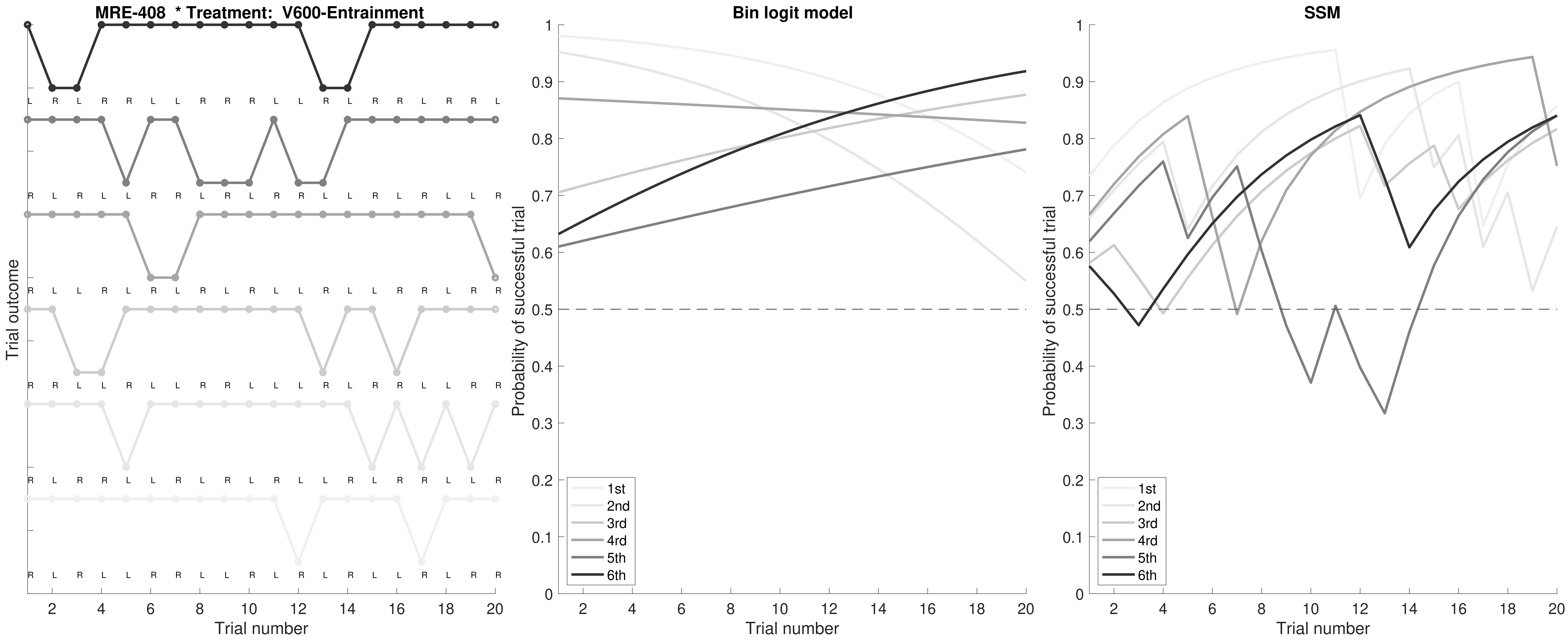

MRE-408 * Treatment: V600-Entrainment
L R L R R L L R R R L L R L R R L R R L
R L R L L R R L R L R R L L R L R L L R
R L L R L R L R L R L L R L L R R L R R
R R L L R L L R R L L L R L R R L R R R
R L R R L L R L R L R L L R L R R L L R
R L R L L R R L L R L R L R L L R L R R
Trial outcome
Trial number
Bin logit model
Probability of successful trial
Trial number
1st
2nd
3rd
4rd
5th
6th
SSM
Probability of successful trial
Trial number
1st
2nd
3rd
4rd
5th
6th

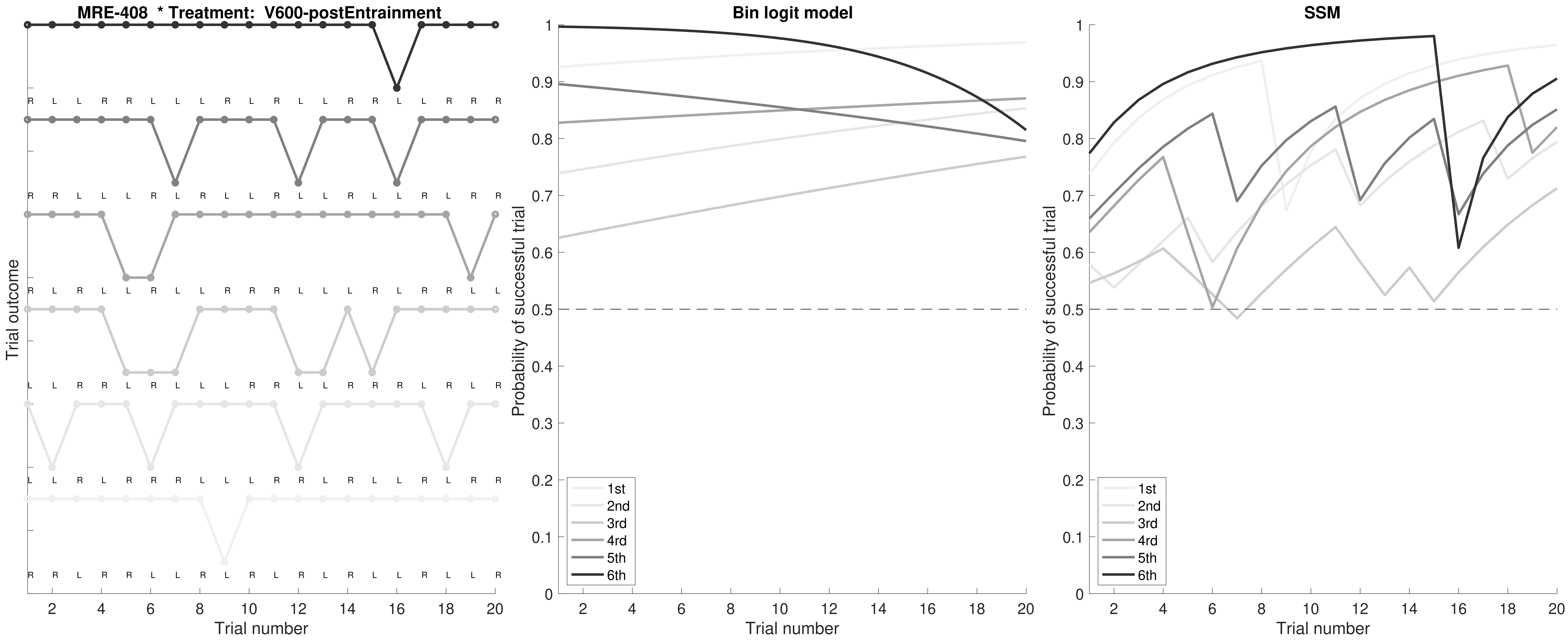

MRE-408 * Treatment: V600-postEntrainment
Bin logit model
SSM
Trial outcome
Trial number
Probability of successful trial
1st
2nd
3rd
4rd
5th
6th

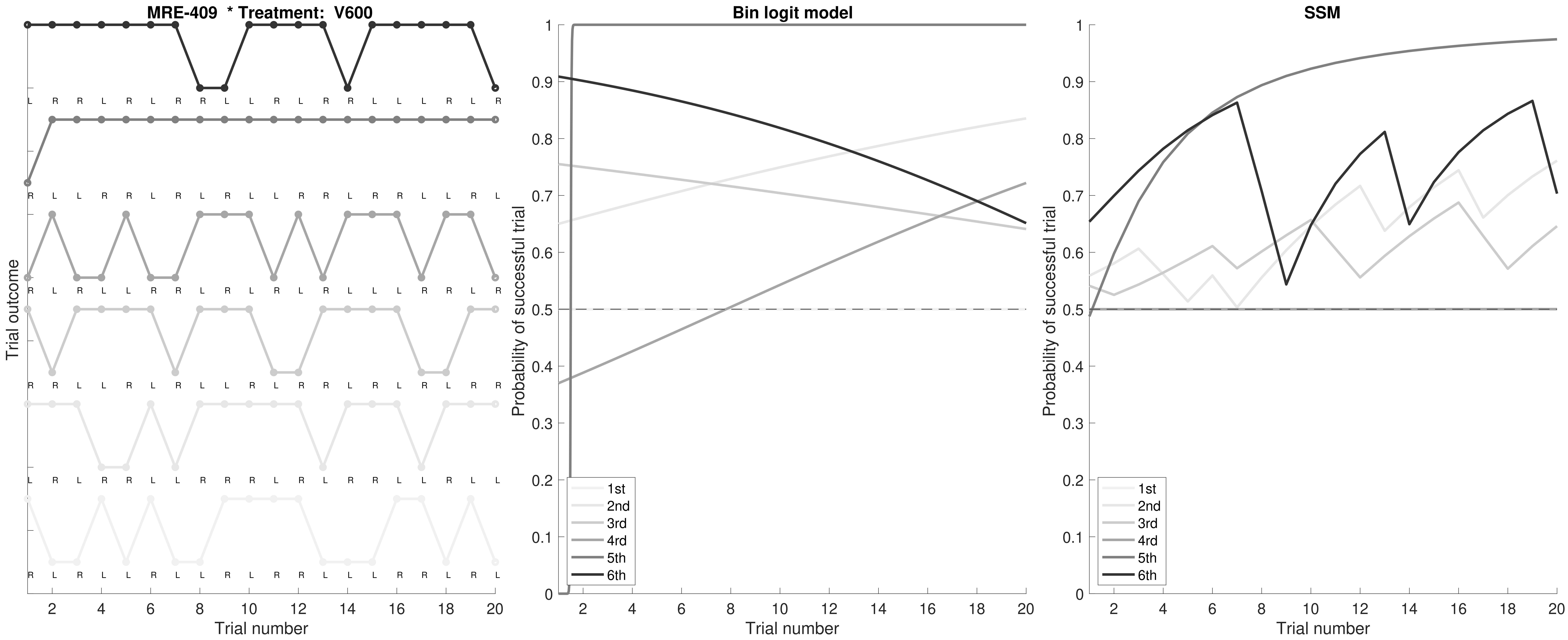

MRE-409 * Treatment: V600
Trial outcome
Trial number
Bin logit model
Probability of successful trial
1st
2nd
3rd
4rd
5th
6th
SSM

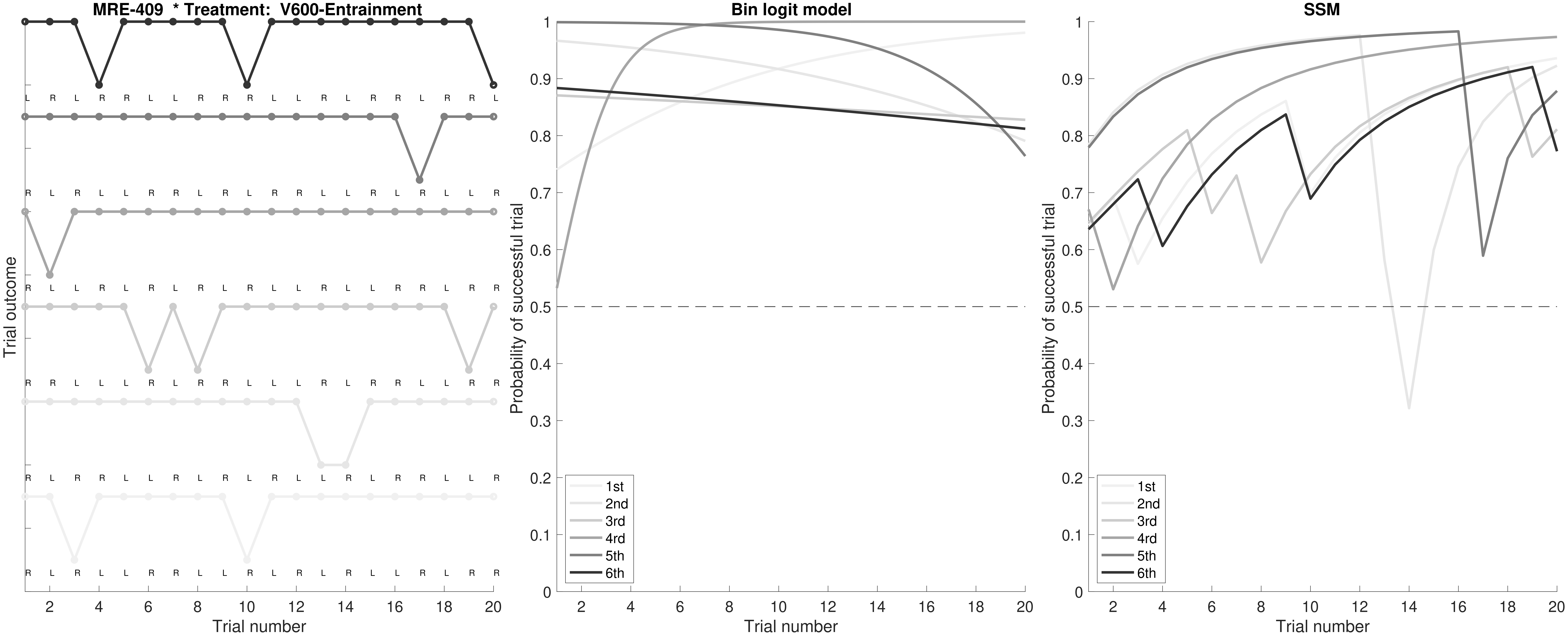
MRE-409 * Treatment: V600-Entrainment
Trial outcome
Trial number
Bin logit model
Probability of successful trial
Trial number
SSM
Probability of successful trial
Trial number
1st
2nd
3rd
4rd
5th
6th

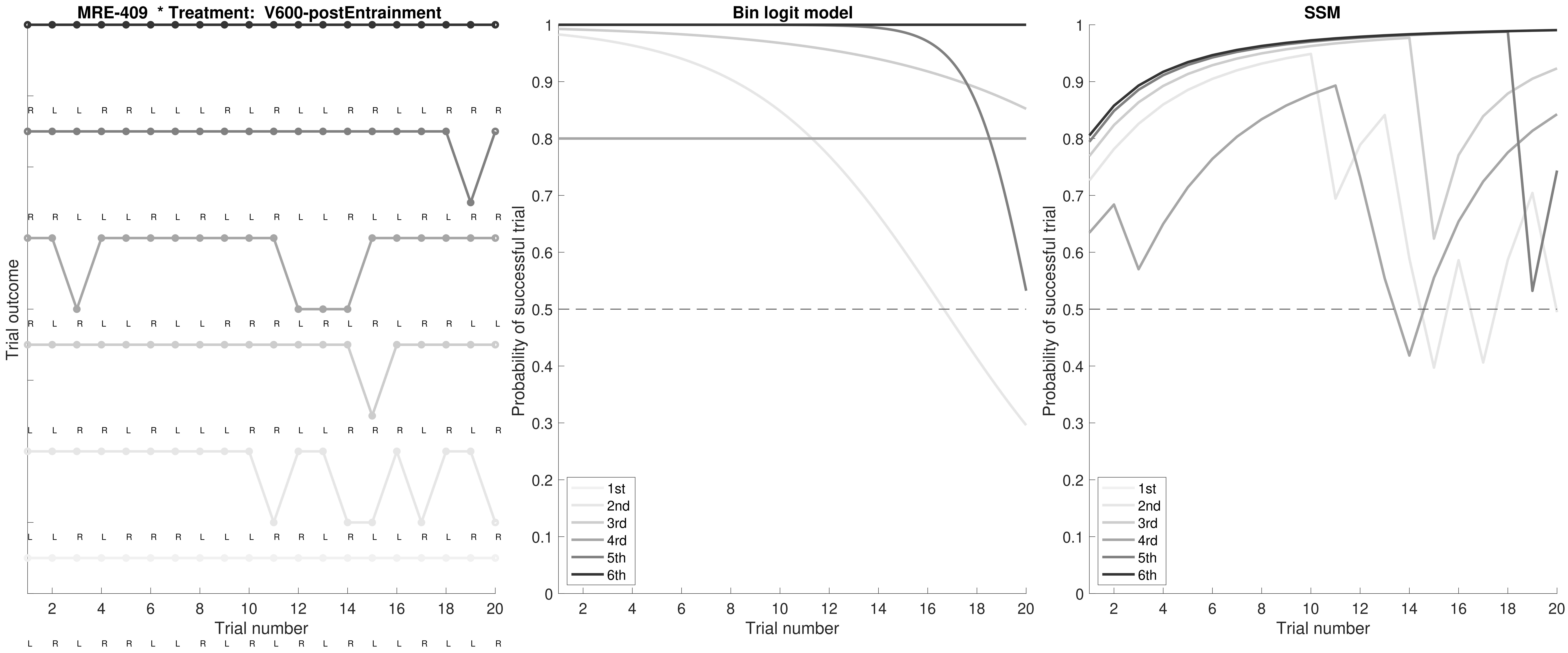
MRE-409 * Treatment: V600-postEntrainment
Bin logit model
SSM
Trial outcome
Trial number
Probability of successful trial
1st
2nd
3rd
4rd
5th
6th

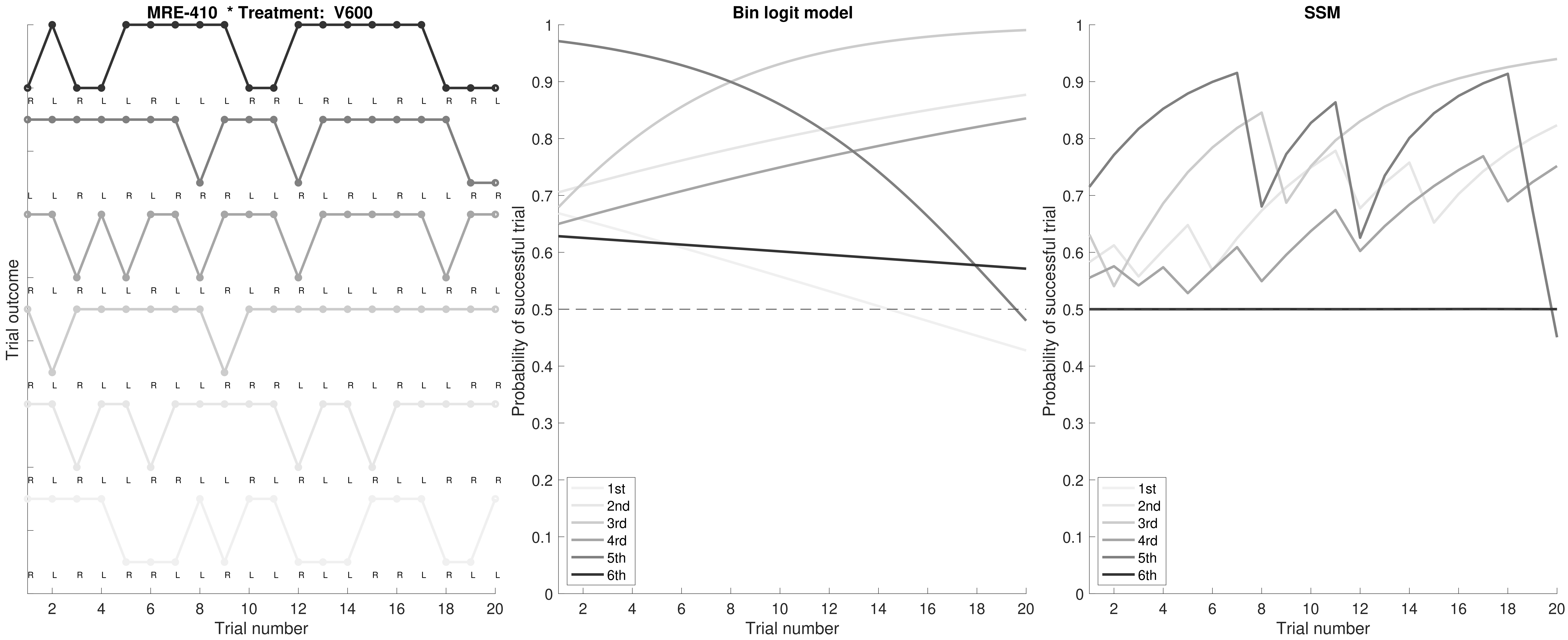

MRE-410 * Treatment: V600
Trial outcome
Trial number
Bin logit model
Probability of successful trial
Trial number
1st
2nd
3rd
4rd
5th
6th
SSM
Probability of successful trial
Trial number
1st
2nd
3rd
4rd
5th
6th

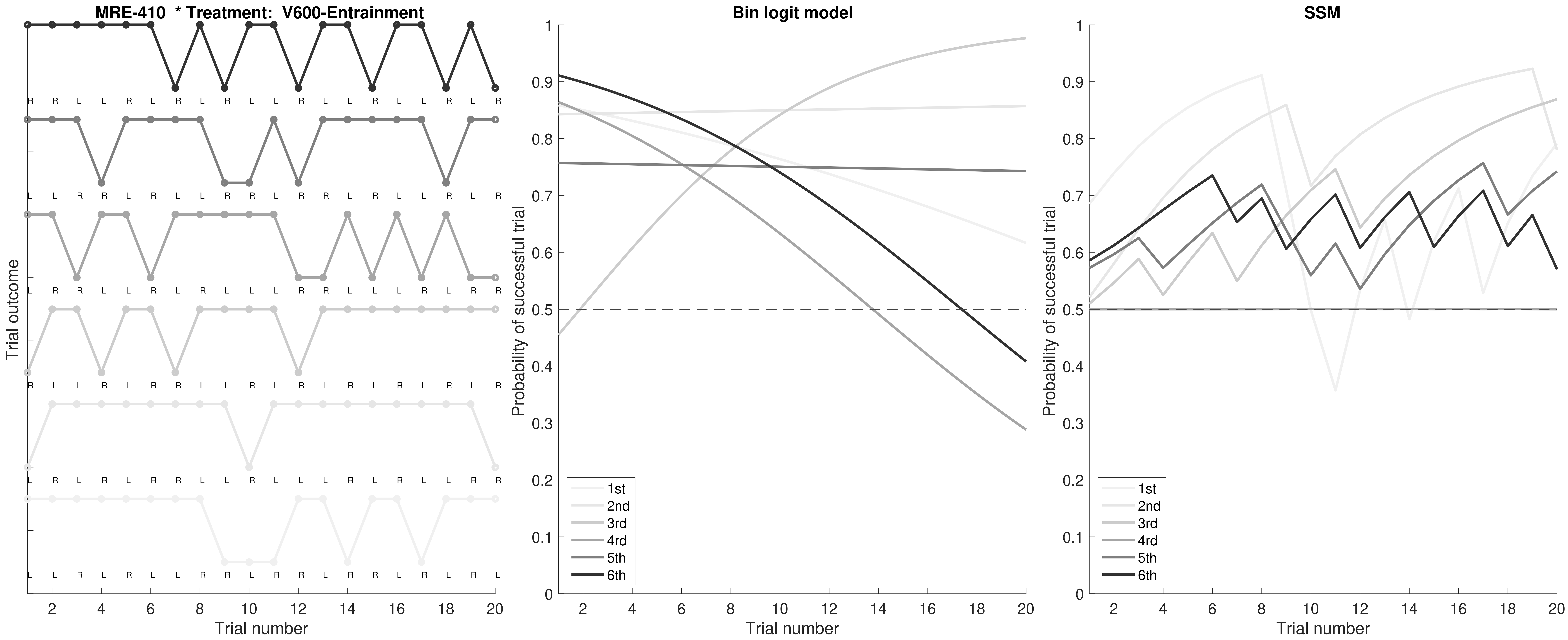

MRE-410 * Treatment: V600-Entrainment
Trial outcome
Trial number
Bin logit model
Probability of successful trial
Trial number
1st
2nd
3rd
4rd
5th
6th
SSM
Probability of successful trial
Trial number
1st
2nd
3rd
4rd
5th
6th

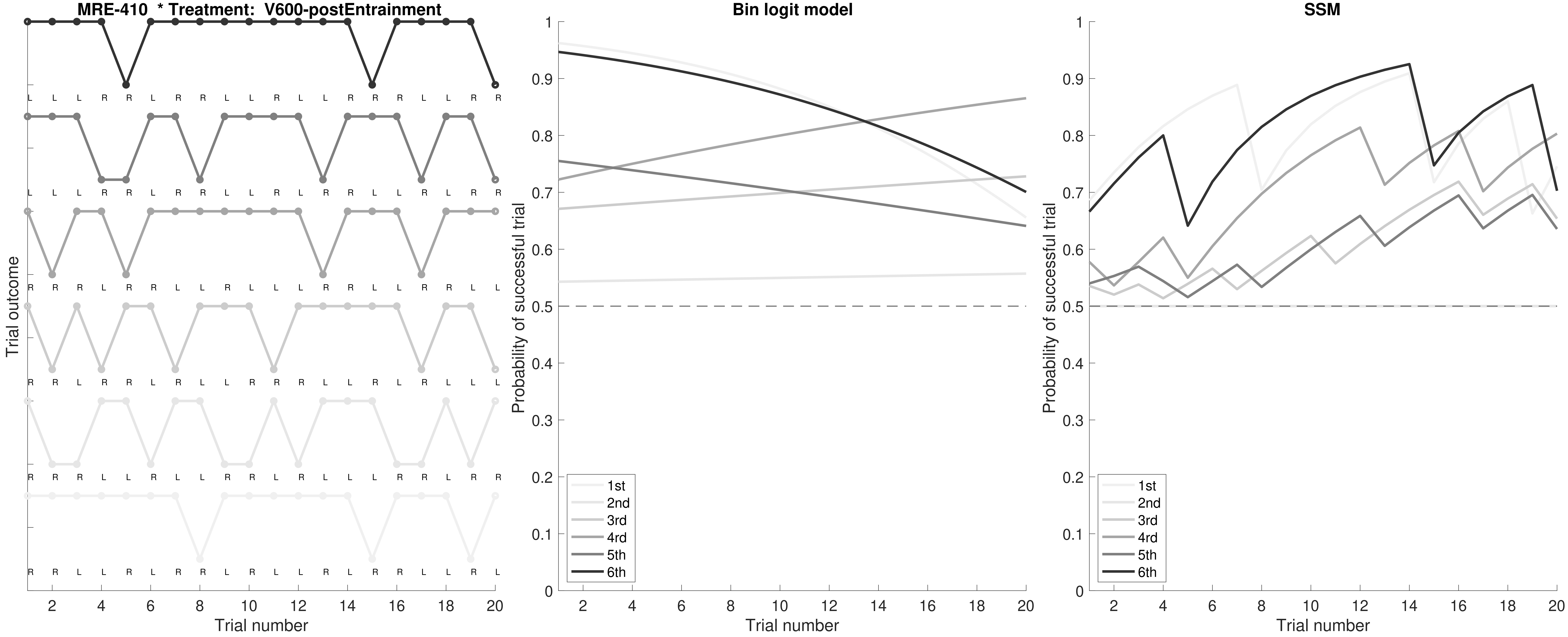

MRE-410 * Treatment: V600-postEntrainment
Trial outcome
L L L R R L R R L L R L L R R R L L R R
L L L R R L R R L L R L R R L L R L R R
R R R L R R L L R L L L R R L L R R L L
R R L R R L R L L R R R L L R L R L L L
R R R L L R L L R R L R L L L R R L R R
R R L L R L R R L R L L R L R R L L R L
Trial number
2 4 6 8 10 12 14 16 18 20
Bin logit model
Probability of successful trial
0 0.1 0.2 0.3 0.4 0.5 0.6 0.7 0.8 0.9 1
1st
2nd
3rd
4rd
5th
6th
SSM

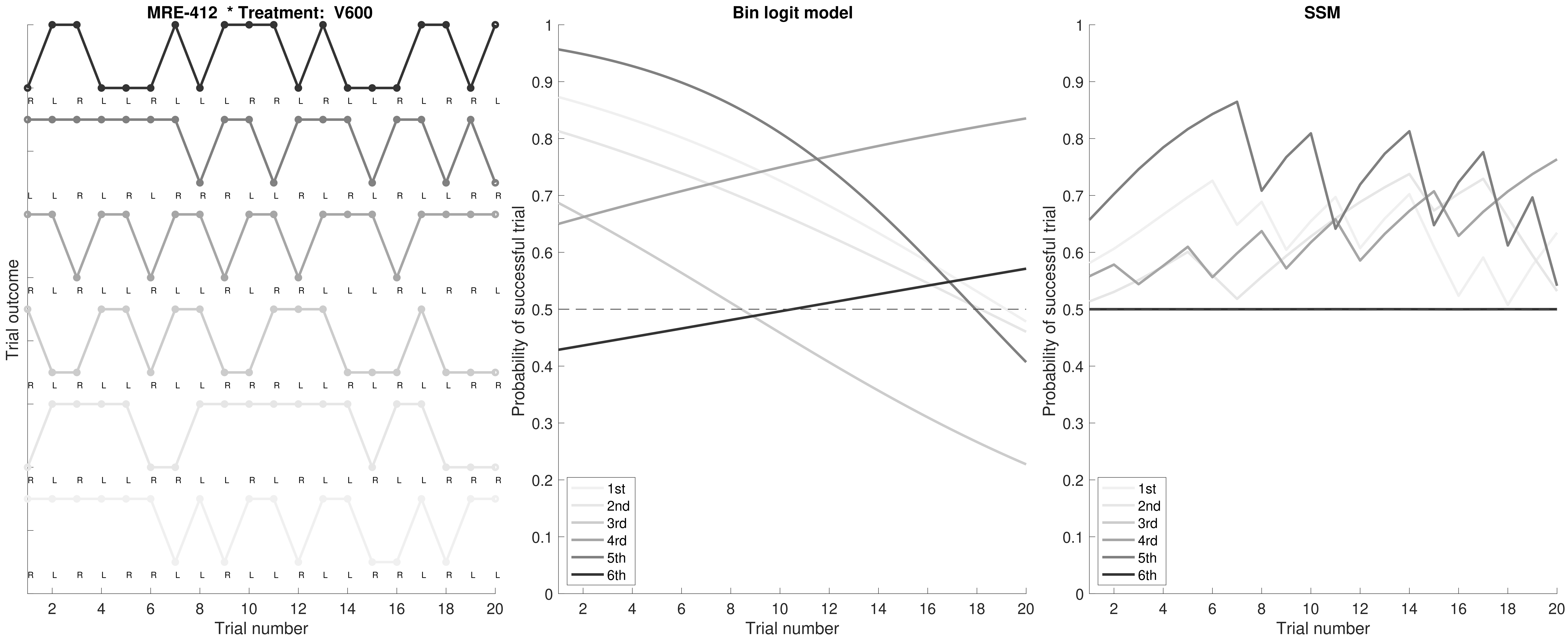

MRE-412 * Treatment: V600
Trial outcome
Trial number
Bin logit model
Probability of successful trial
SSM
1st
2nd
3rd
4rd
5th
6th

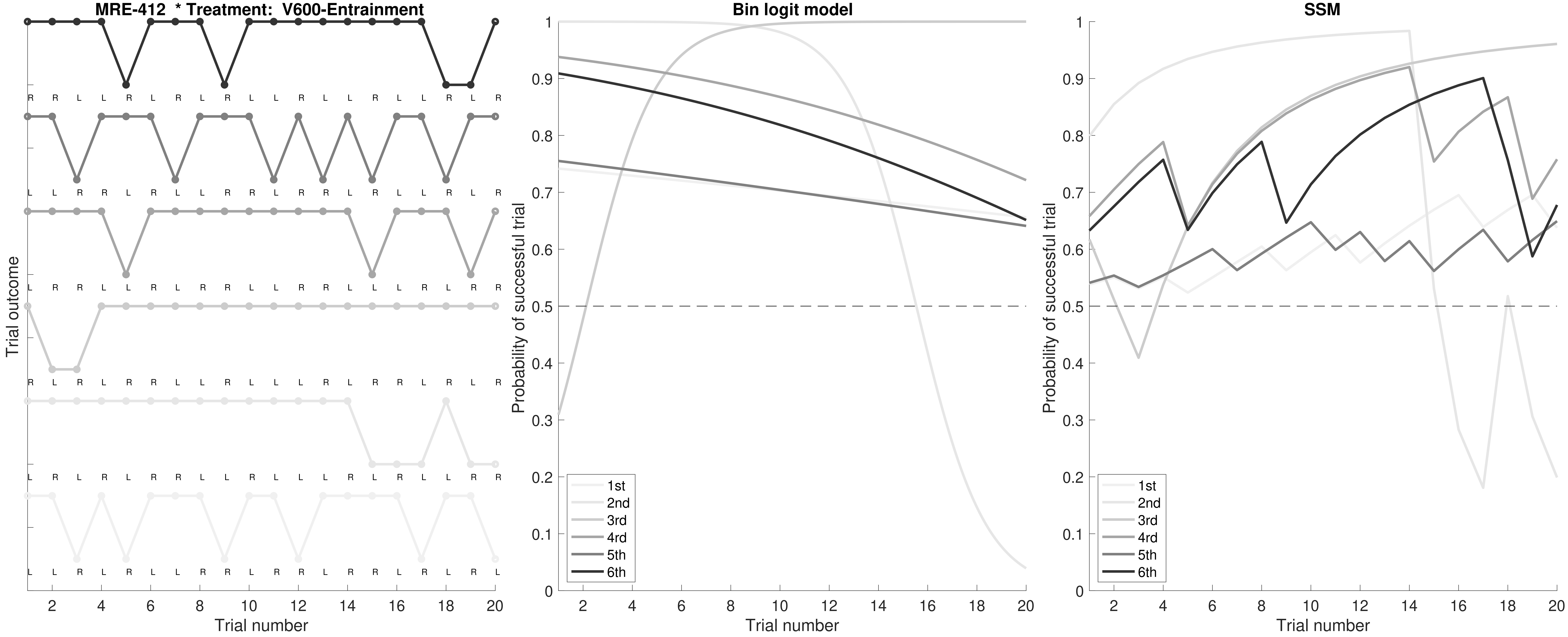

MRE-412 * Treatment: V600-Entrainment
Trial outcome
Trial number
Bin logit model
Probability of successful trial
Trial number
1st
2nd
3rd
4rd
5th
6th
SSM
Probability of successful trial
Trial number
1st
2nd
3rd
4rd
5th
6th

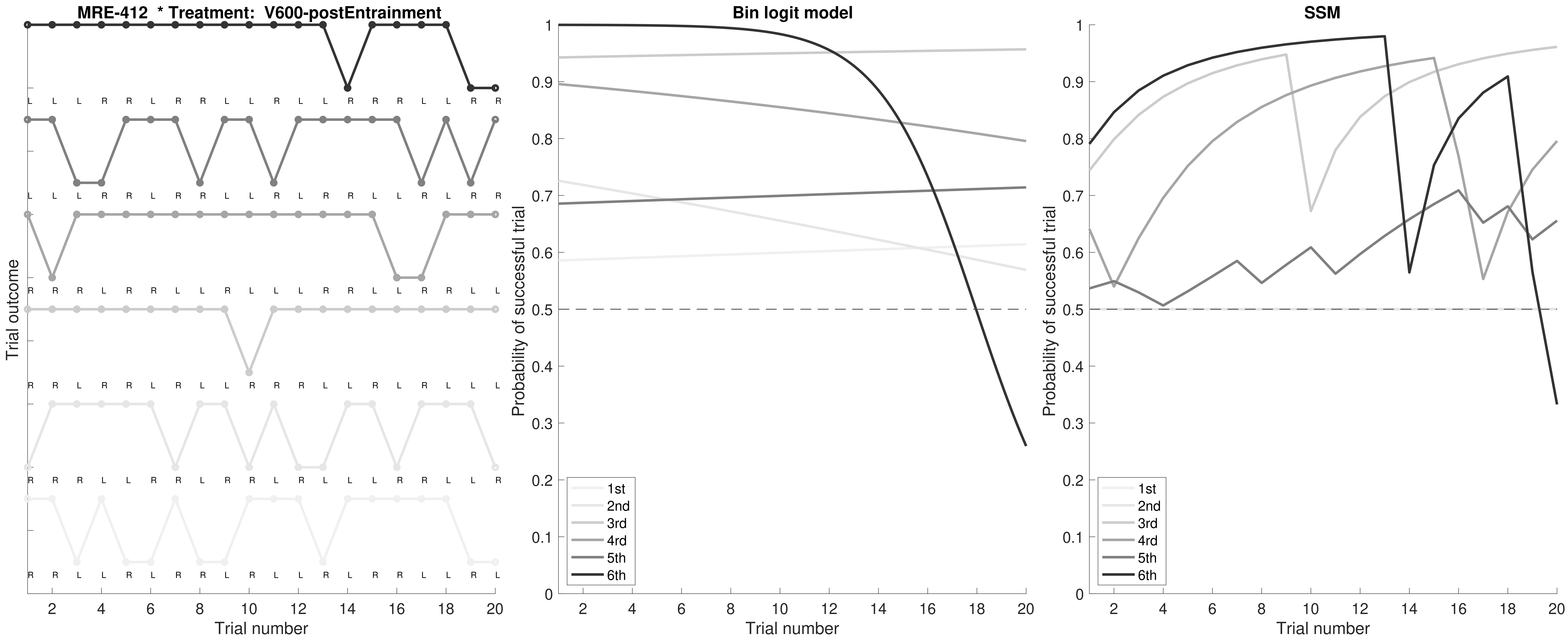

MRE-412 * Treatment: V600-postEntrainment
Trial outcome
L L L R R L R R L L R L L R R R L L R R
L L L R R L R R L L R L R R L L R L R R
R R R L R R L L R L L L R R L L R R L L
R R L R R L R L L R R R L L R L R L L L
R R R L L R R L R R L R L L L R R L L R
R R L L R L R R L R R L R L R R L L R L
Trial number
Bin logit model
Probability of successful trial
1st
2nd
3rd
4rd
5th
6th
Trial number
SSM
Probability of successful trial
1st
2nd
3rd
4rd
5th
6th
Trial number

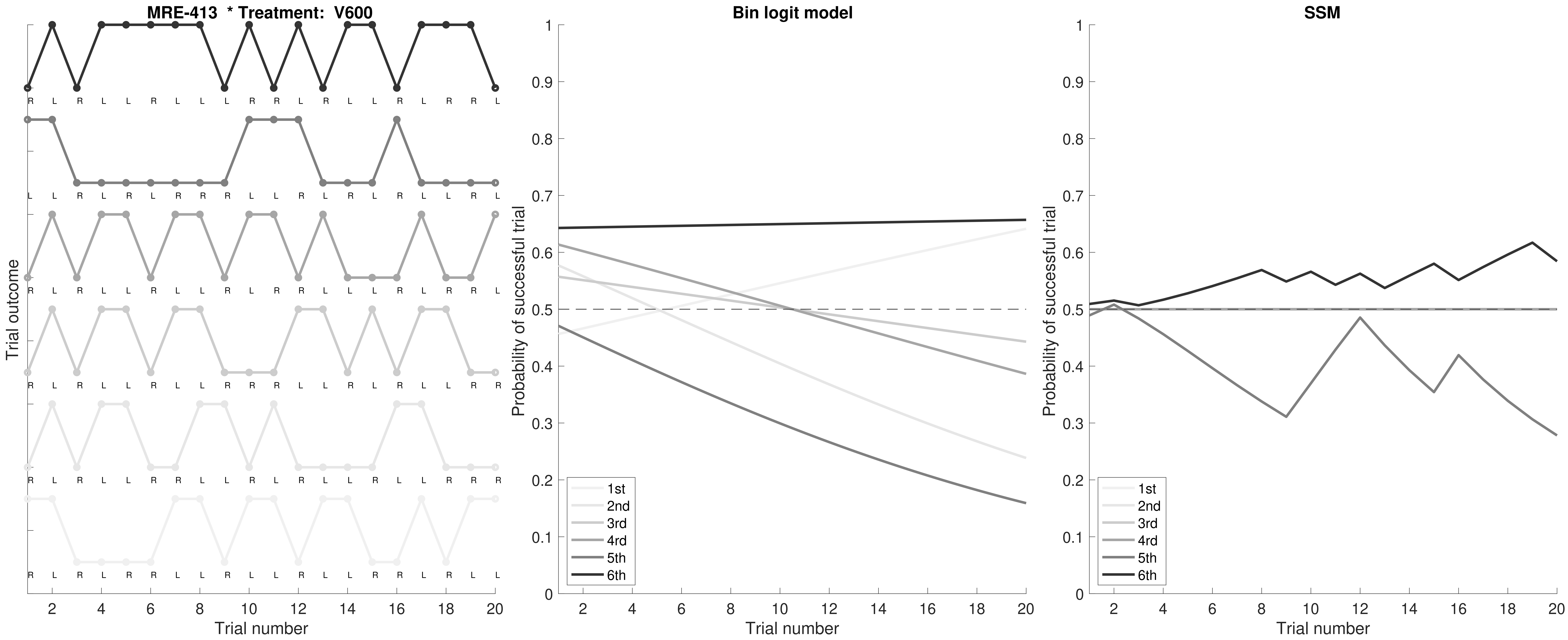

MRE-413 * Treatment: V600
Trial outcome
Trial number
Bin logit model
Probability of successful trial
Trial number
1st
2nd
3rd
4rd
5th
6th
SSM
Probability of successful trial
Trial number
1st
2nd
3rd
4rd
5th
6th

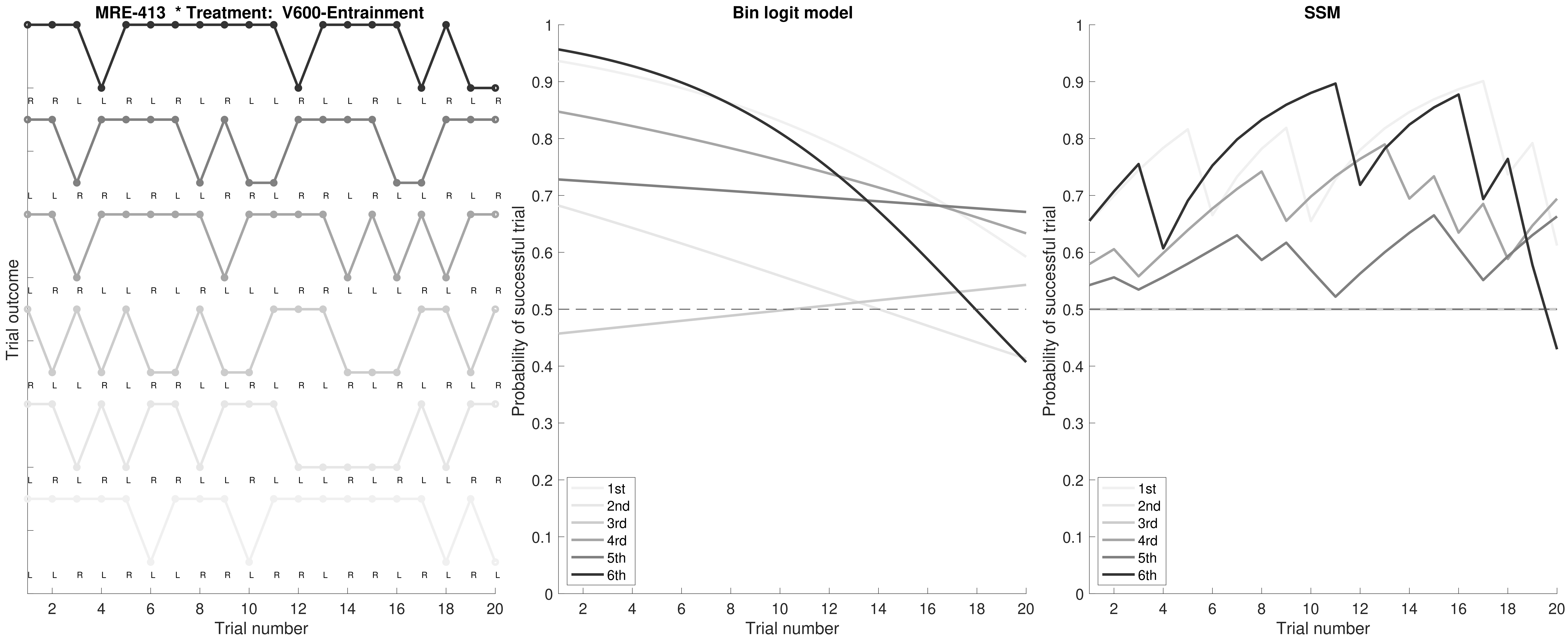

MRE-413 * Treatment: V600-Entrainment
Trial outcome
Trial number
Bin logit model
Probability of successful trial
Trial number
1st
2nd
3rd
4rd
5th
6th
SSM
Probability of successful trial
Trial number
1st
2nd
3rd
4rd
5th
6th

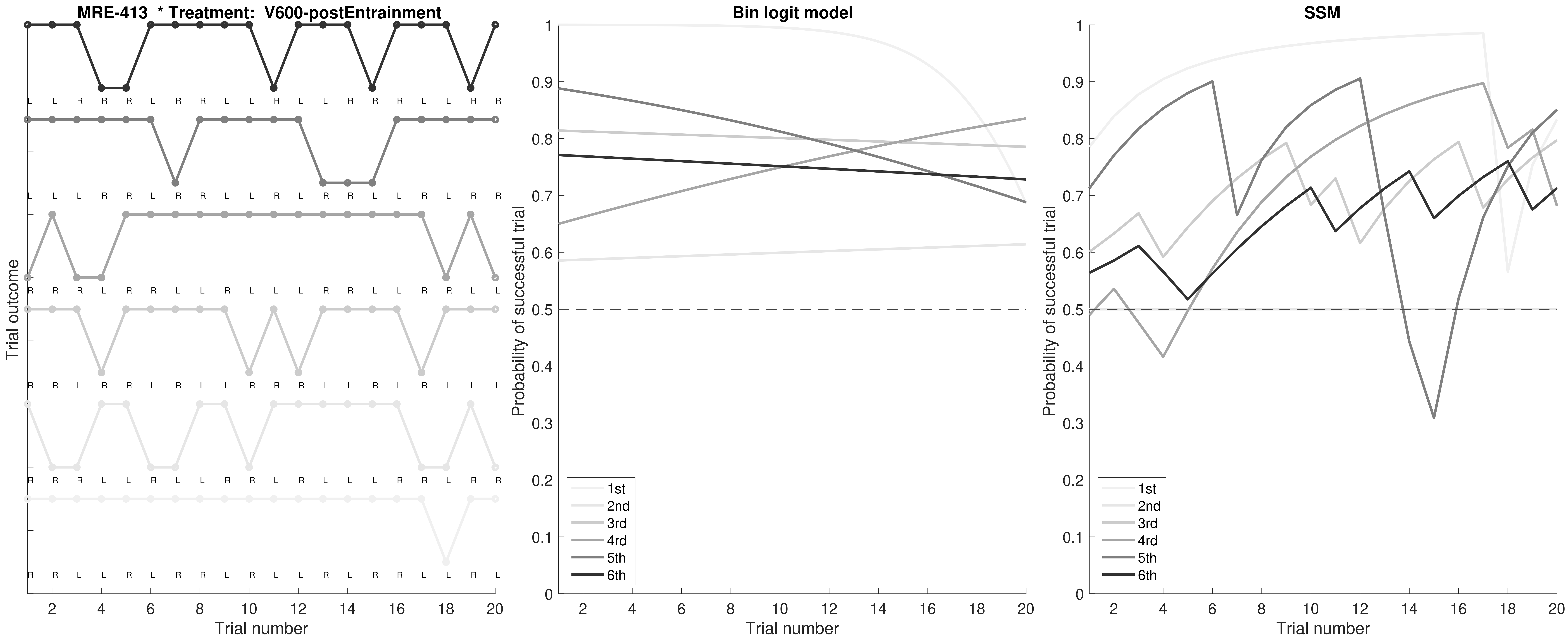

MRE-413 * Treatment: V600-postEntrainment
Trial outcome
Trial number
Bin logit model
Probability of successful trial
Trial number
1st
2nd
3rd
4rd
5th
6th
SSM
Probability of successful trial
Trial number
1st
2nd
3rd
4rd
5th
6th

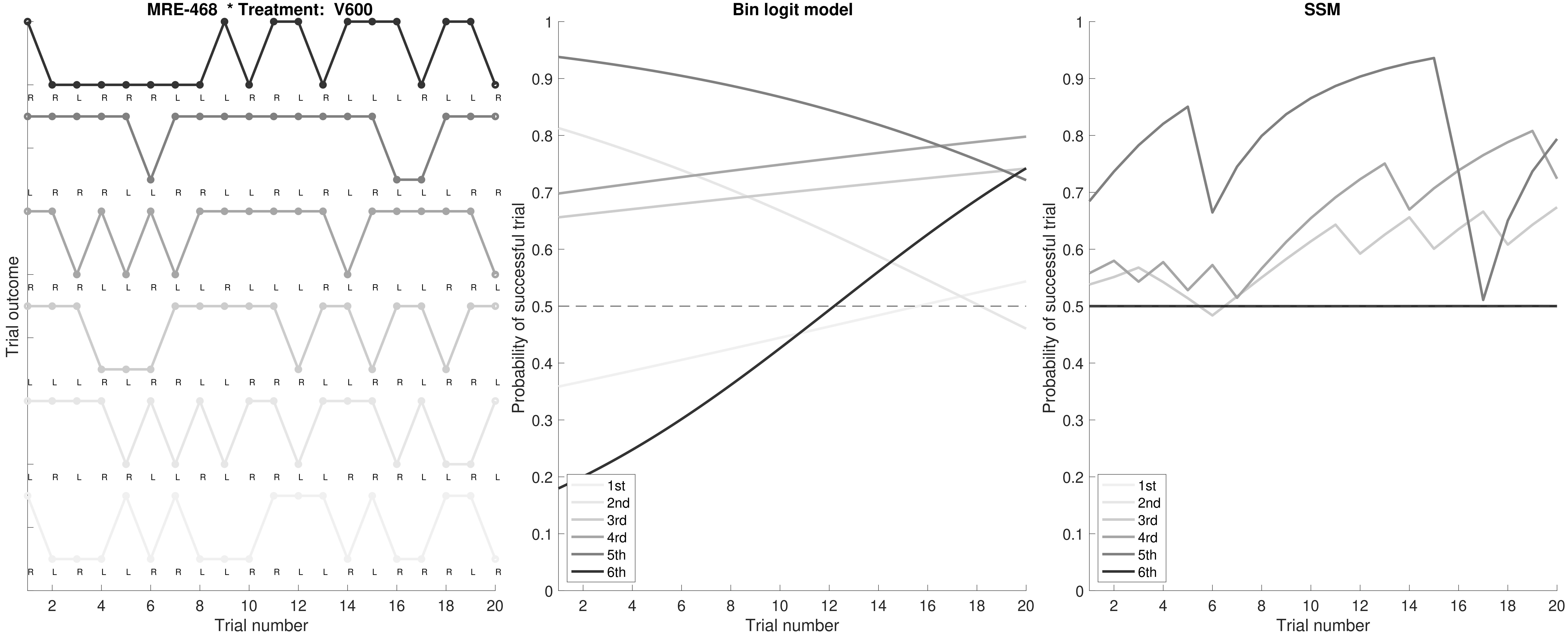
MRE-468 * Treatment: V600
Trial outcome
Trial number
Bin logit model
Probability of successful trial
Trial number
1st
2nd
3rd
4rd
5th
6th
SSM
Probability of successful trial
Trial number
1st
2nd
3rd
4rd
5th
6th

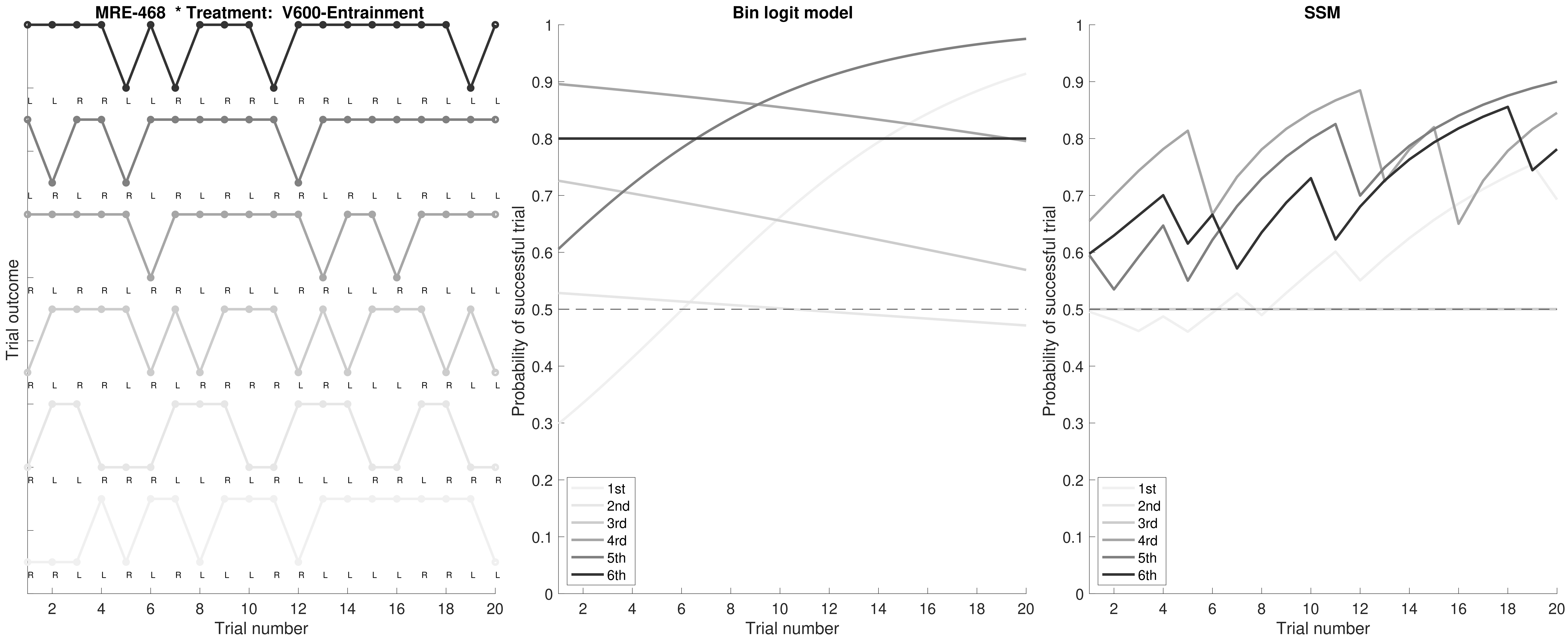
MRE-468 * Treatment: V600-Entrainment
L L R R L L R L R R L R R L R L R L L L
L R L R R L R L R R R R L R L L R L L L
R L R R L R R L R L R L R L L R R L L L
R L R R L R L R R R R L R L L R L R L L
R L L R R R L L R L R L L L R R L R R R
R R L L R L R L L L R R L L L L R R L L
Trial outcome
Trial number
Bin logit model
Probability of successful trial
Trial number
1st
2nd
3rd
4rd
5th
6th
SSM
Probability of successful trial
Trial number
1st
2nd
3rd
4rd
5th
6th

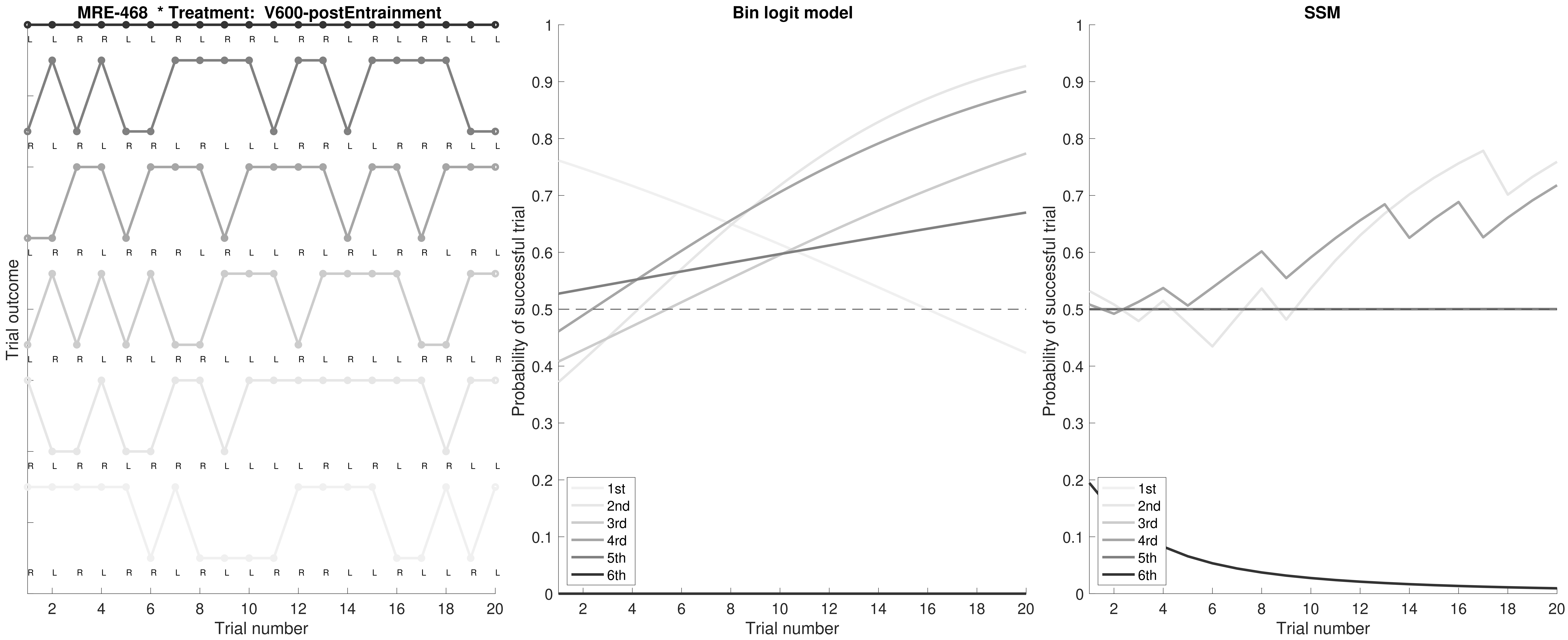
MRE-468 * Treatment: V600-postEntrainment
Trial outcome
Trial number
Bin logit model
Probability of successful trial
Trial number
SSM
Probability of successful trial
Trial number
1st
2nd
3rd
4rd
5th
6th
