## Supplemental Figure 2 for "Auditory Gamma-Frequency Entrainment Abolishes Working Memory Deficits in a Rodent Model of Autism"

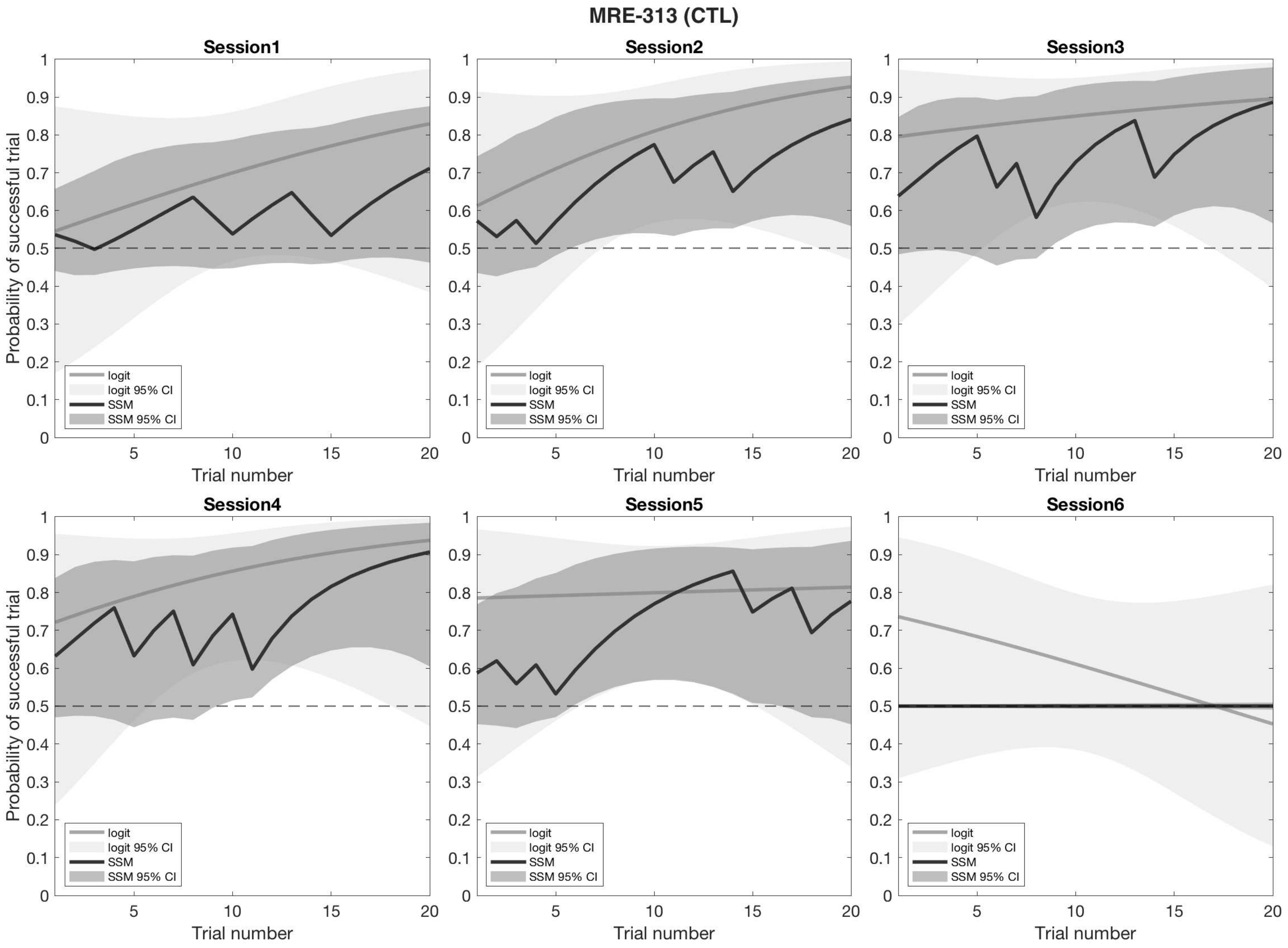
MRE-313 (CTL)
Session1
Session2
Session3
Session4
Session5
Session6
Probability of successful trial
Trial number
logit
logit 95% CI
SSM
SSM 95% CI

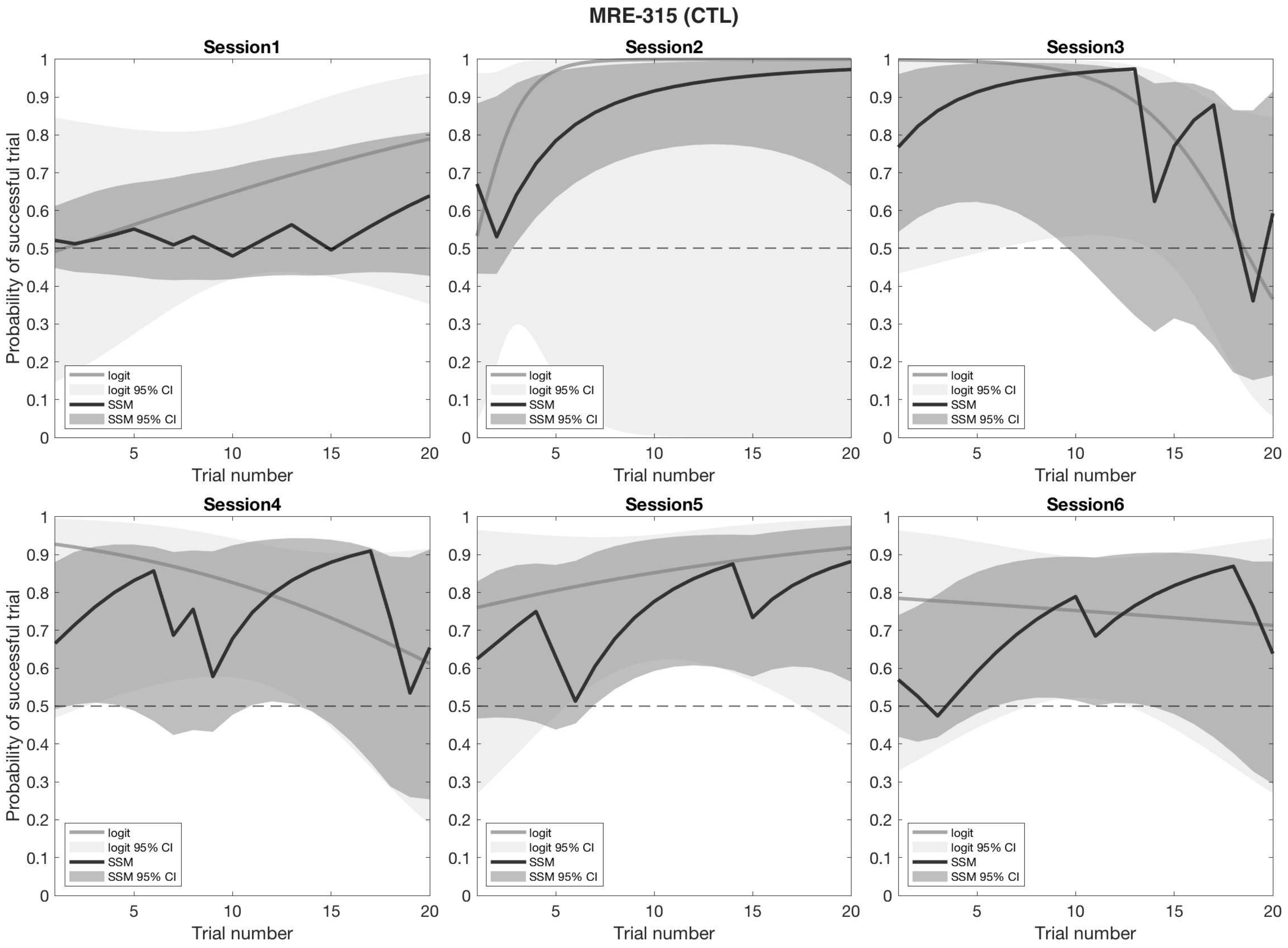

MRE-315 (CTL)
Session1
Session2
Session3
Session4
Session5
Session6
Probability of successful trial
Trial number
logit
logit 95% CI
SSM
SSM 95% CI

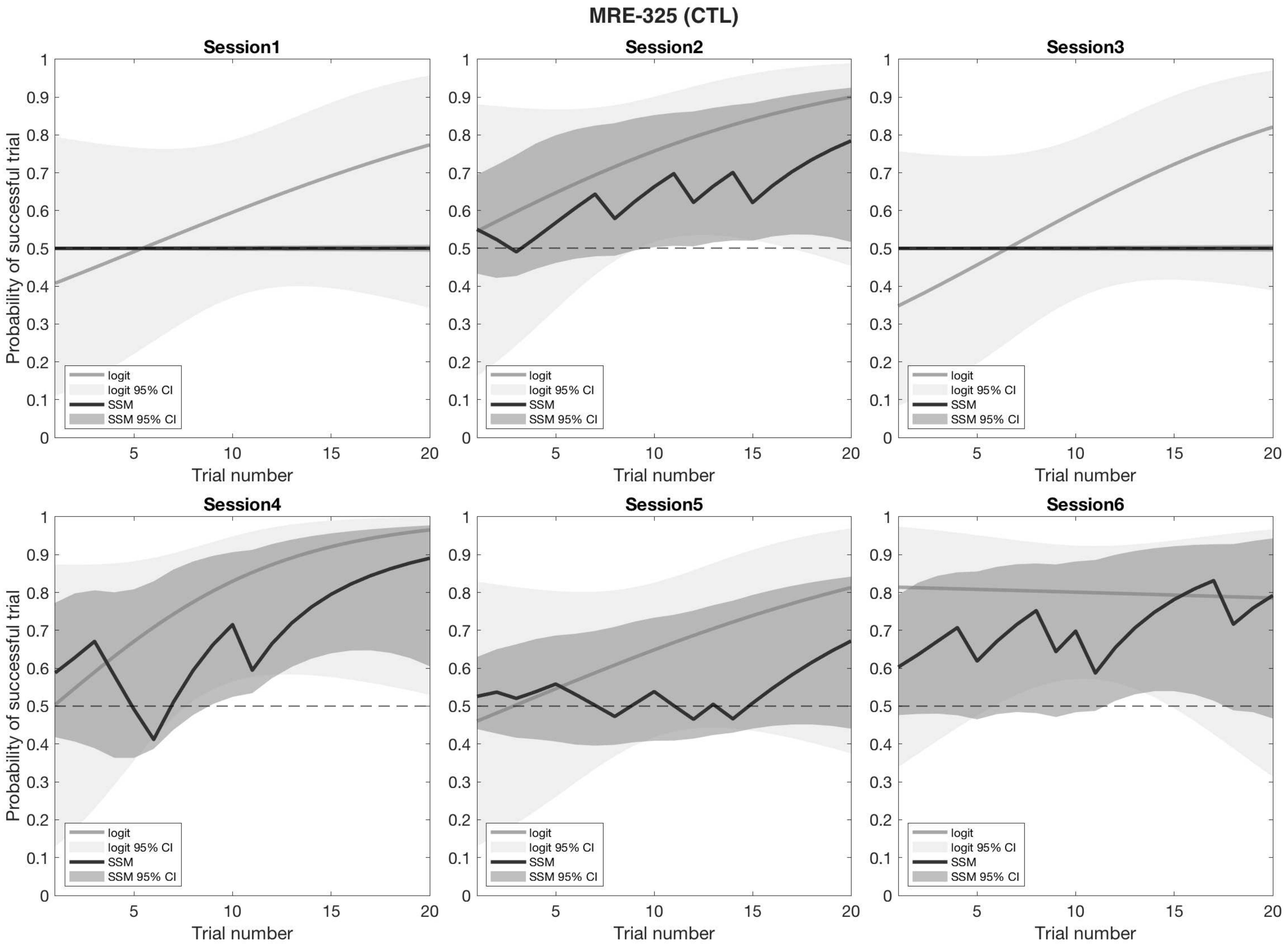

MRE-325 (CTL)
Session1
Session2
Session3
Session4
Session5
Session6
Probability of successful trial
Trial number
logit
logit 95% CI
SSM
SSM 95% CI

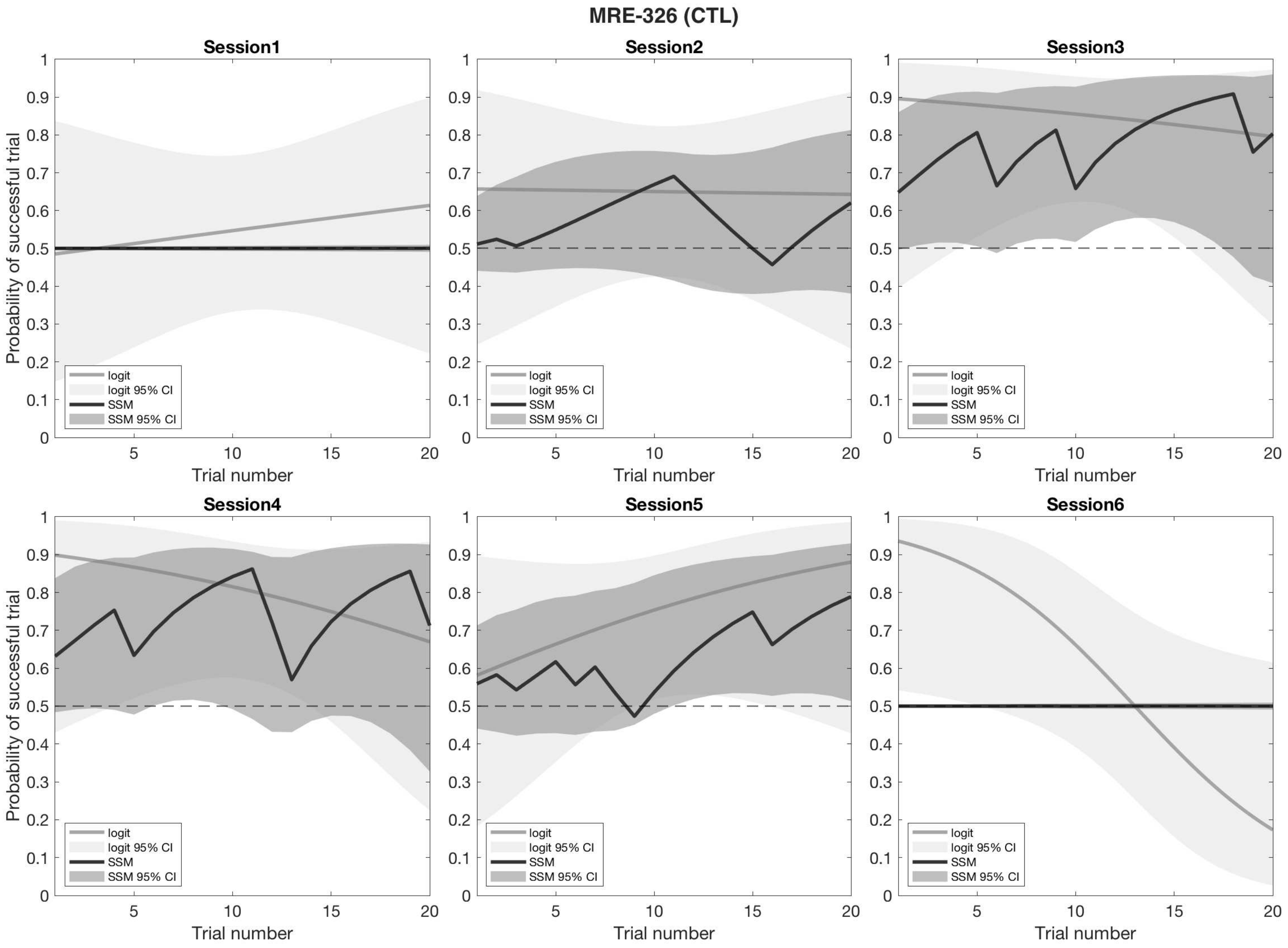
MRE-326 (CTL)
Session1
Session2
Session3
Session4
Session5
Session6
Probability of successful trial
Trial number
logit
logit 95% CI
SSM
SSM 95% CI

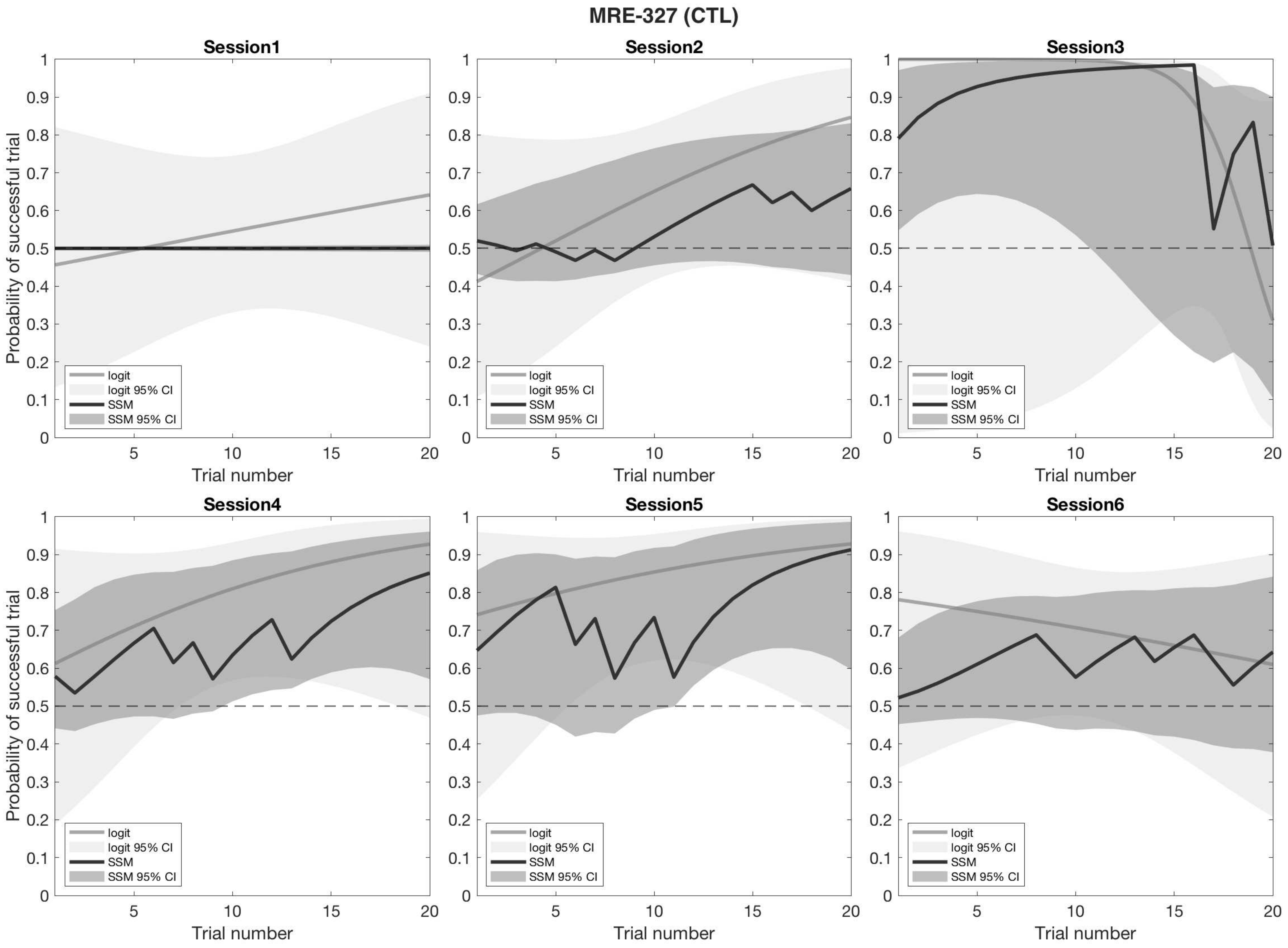

MRE-327 (CTL)
Session1
Session2
Session3
Session4
Session5
Session6
Probability of successful trial
Trial number
logit
logit 95% CI
SSM
SSM 95% CI

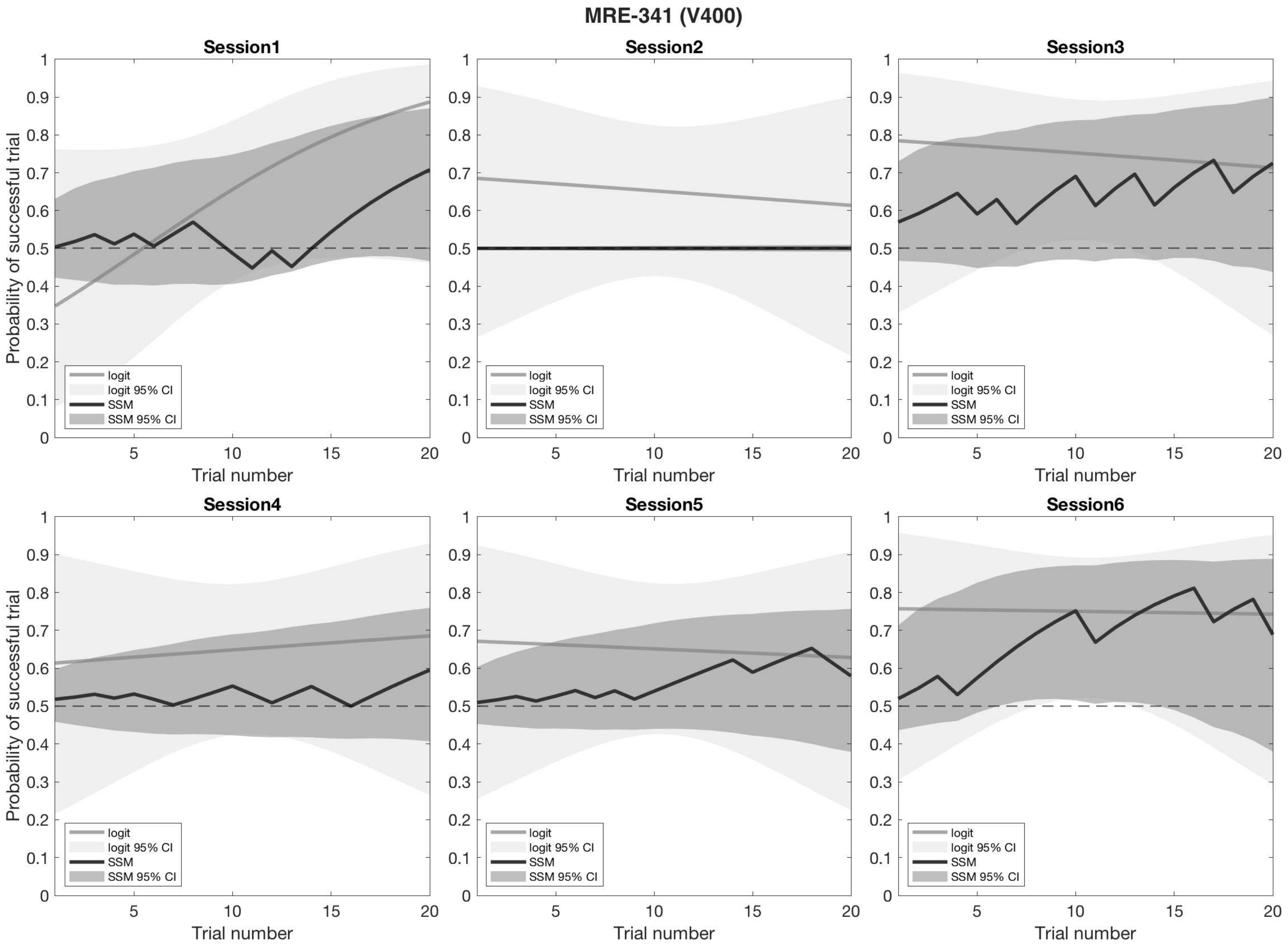
MRE-341 (V400)
Session1
Session2
Session3
Session4
Session5
Session6
Probability of successful trial
Trial number
logit
logit 95% CI
SSM
SSM 95% CI
0
0.1
0.2
0.3
0.4
0.5
0.6
0.7
0.8
0.9
1
5
10
15
20

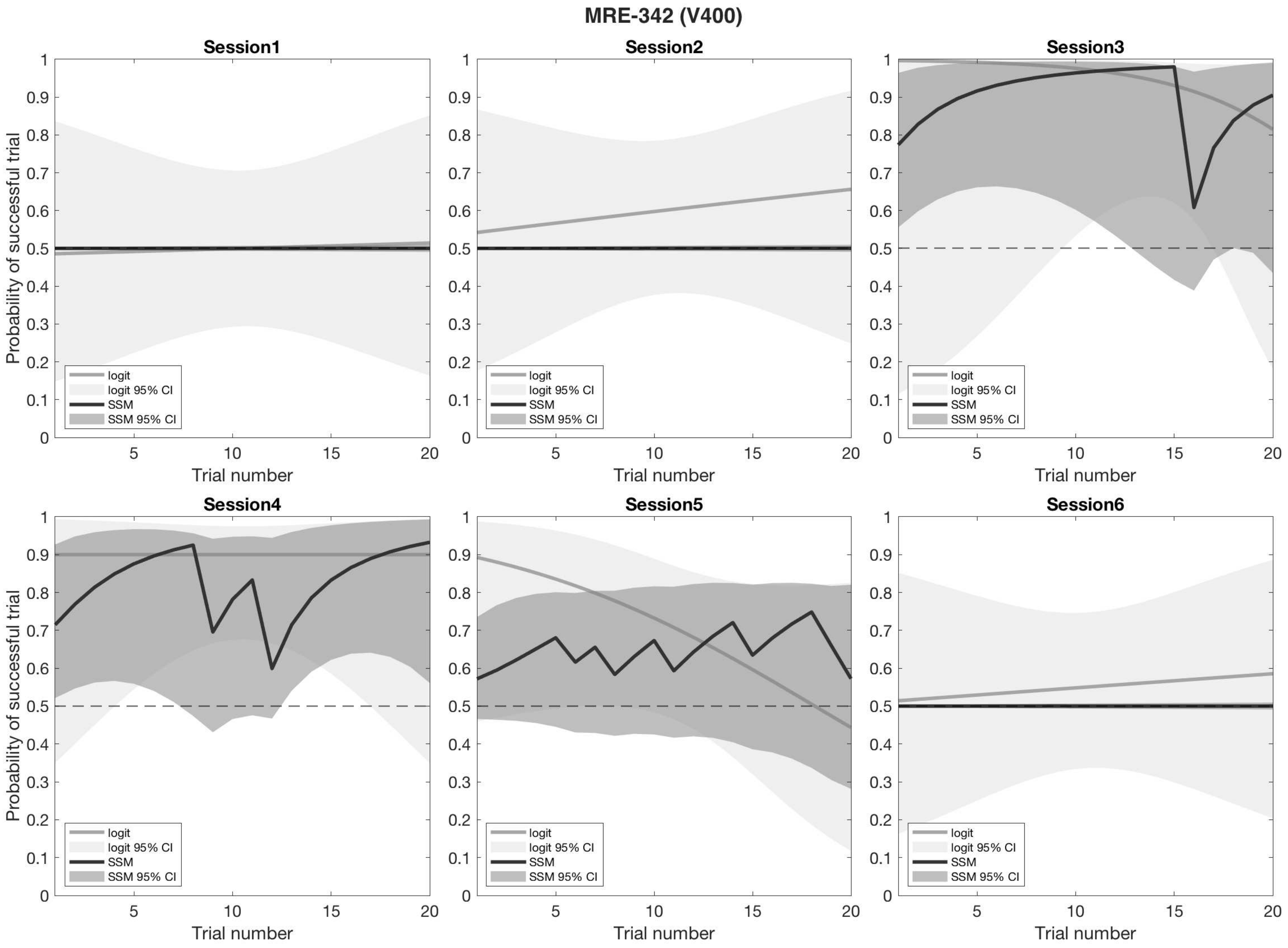
MRE-342 (V400)
Session1
Session2
Session3
Session4
Session5
Session6
Probability of successful trial
Trial number
logit
logit 95% CI
SSM
SSM 95% CI

### MRE-343 (V400)

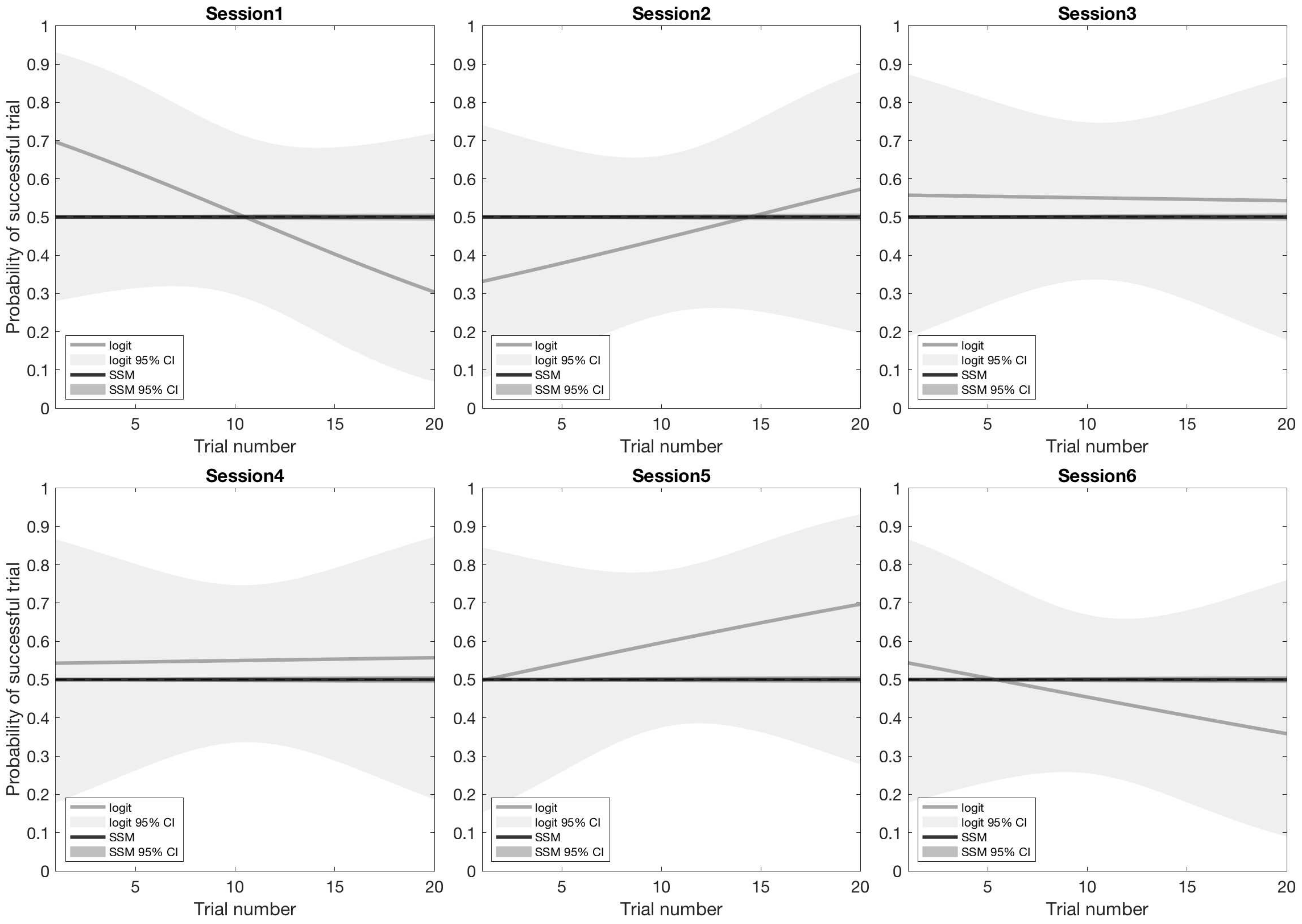

Session1
Session2
Session3
Session4
Session5
Session6
Probability of successful trial
Trial number
logit
logit 95% CI
SSM
SSM 95% CI

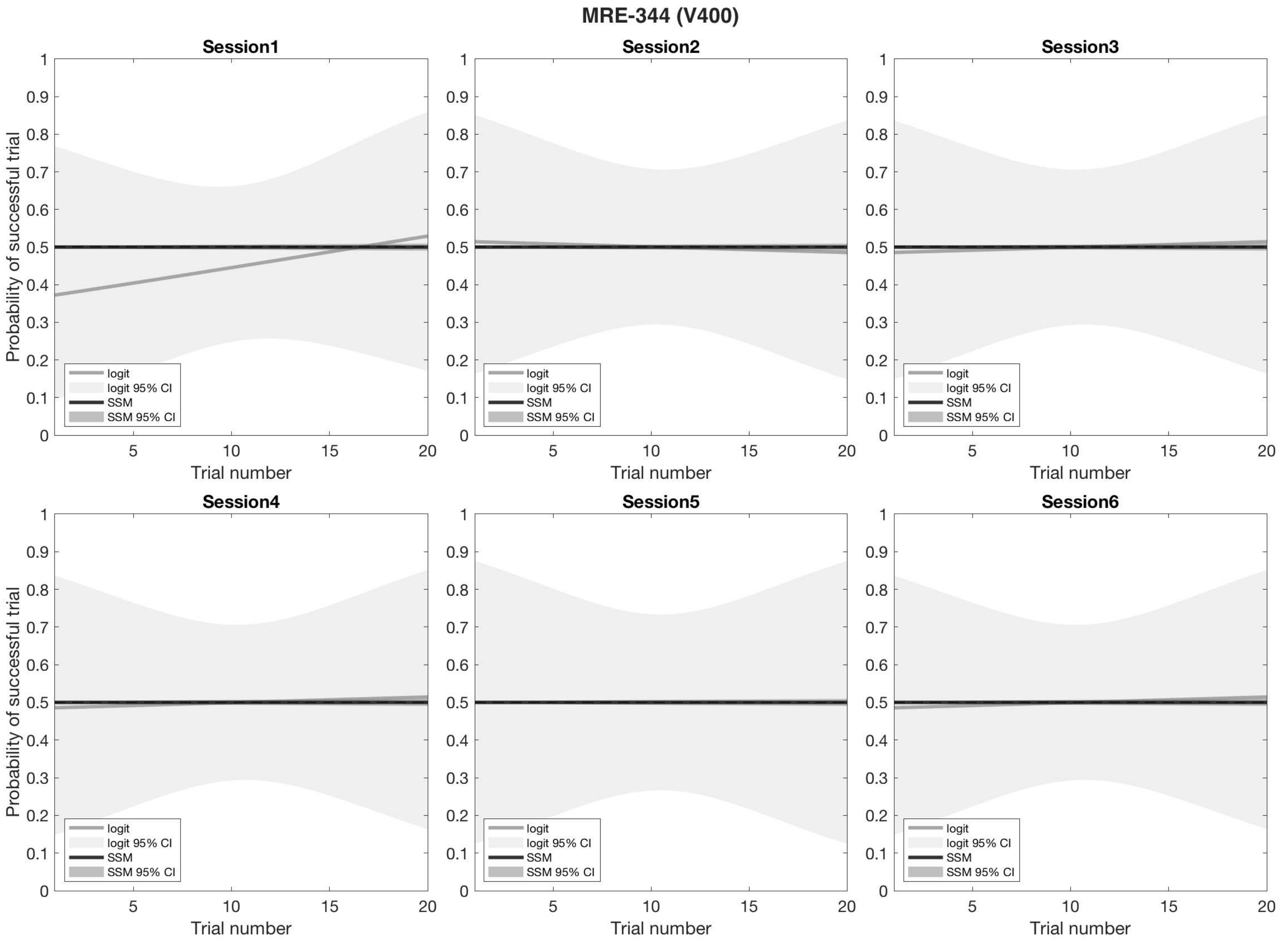
MRE-344 (V400)
Session1
Session2
Session3
Session4
Session5
Session6
Probability of successful trial
Trial number
logit
logit 95% CI
SSM
SSM 95% CI

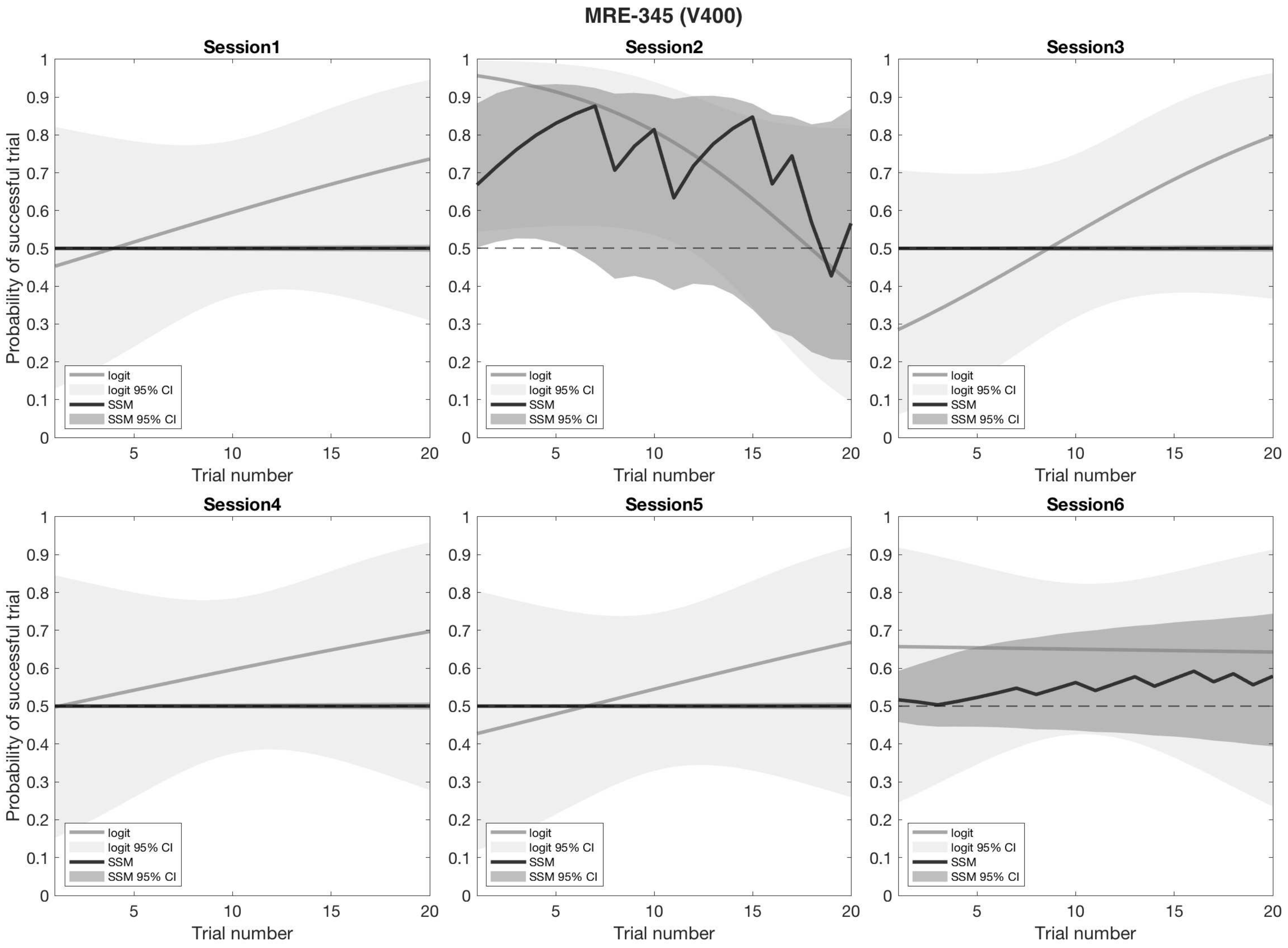

MRE-345 (V400)
Session1
Session2
Session3
Session4
Session5
Session6
Probability of successful trial
Trial number
logit
logit 95% CI
SSM
SSM 95% CI

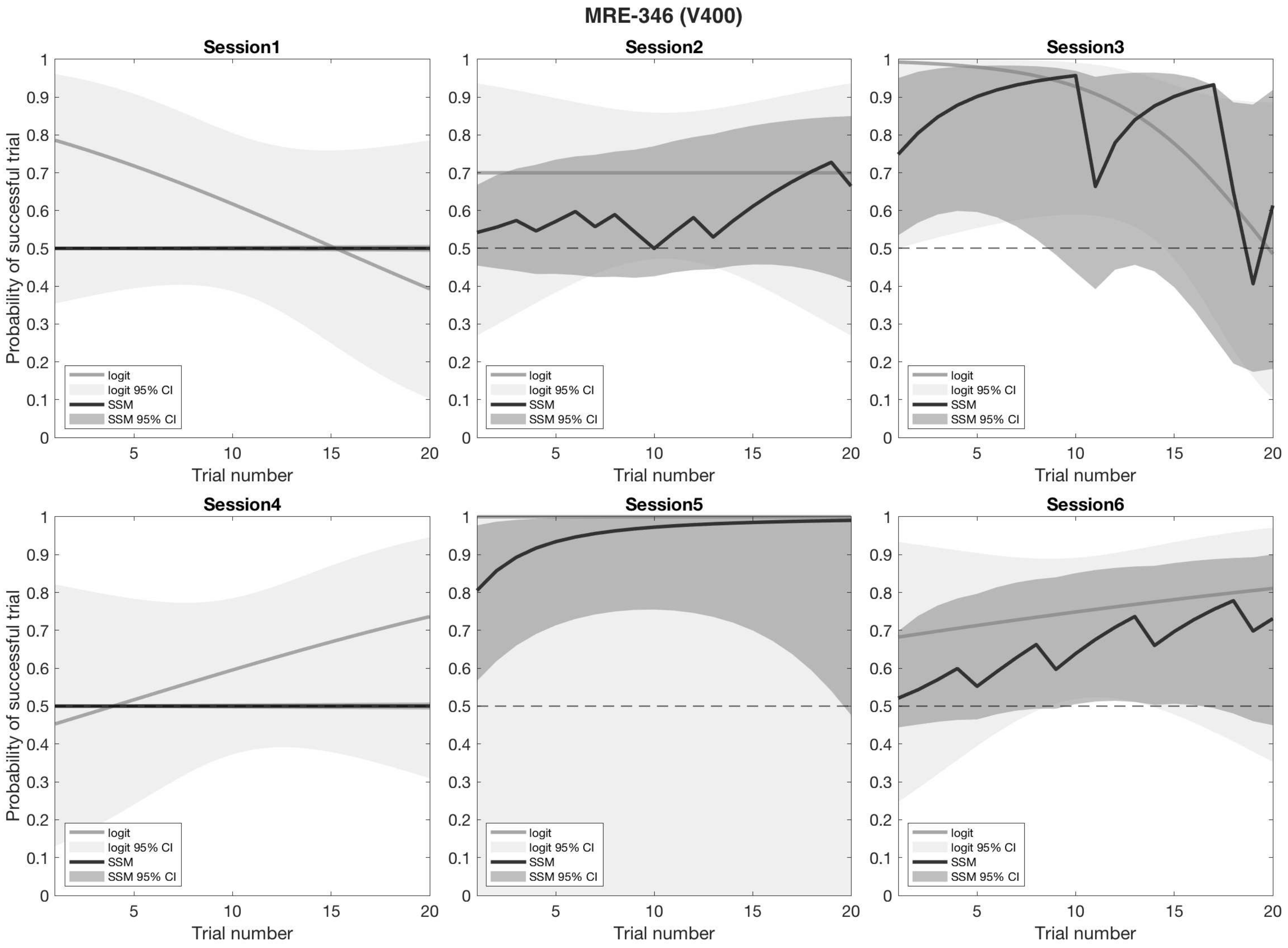
MRE-346 (V400)
Session1
Session2
Session3
Session4
Session5
Session6
Probability of successful trial
Trial number
logit
logit 95% CI
SSM
SSM 95% CI

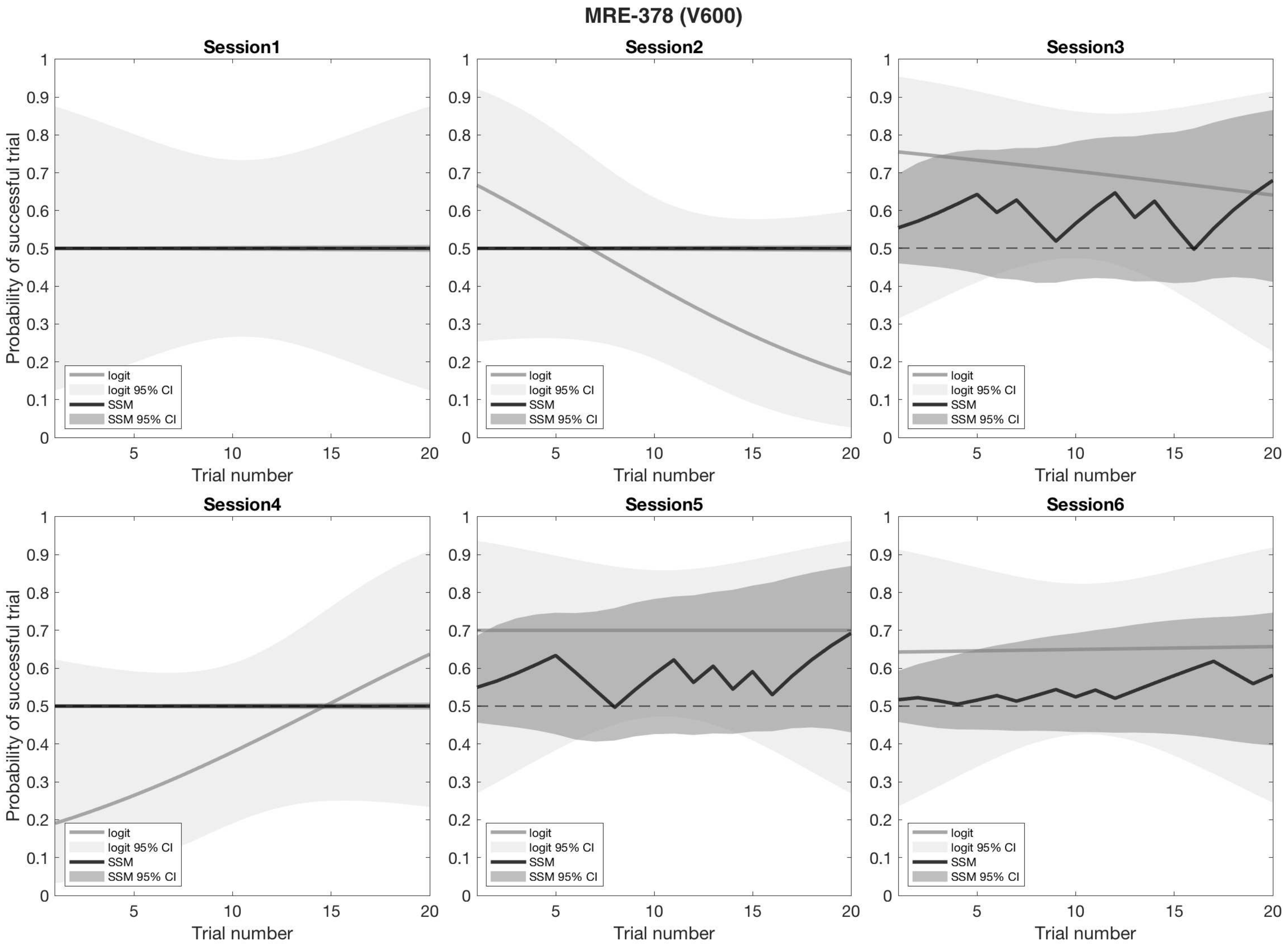
MRE-378 (V600)
Session1
Session2
Session3
Session4
Session5
Session6
Probability of successful trial
Trial number
logit
logit 95% CI
SSM
SSM 95% CI
0
0.1
0.2
0.3
0.4
0.5
0.6
0.7
0.8
0.9
1
5
10
15
20

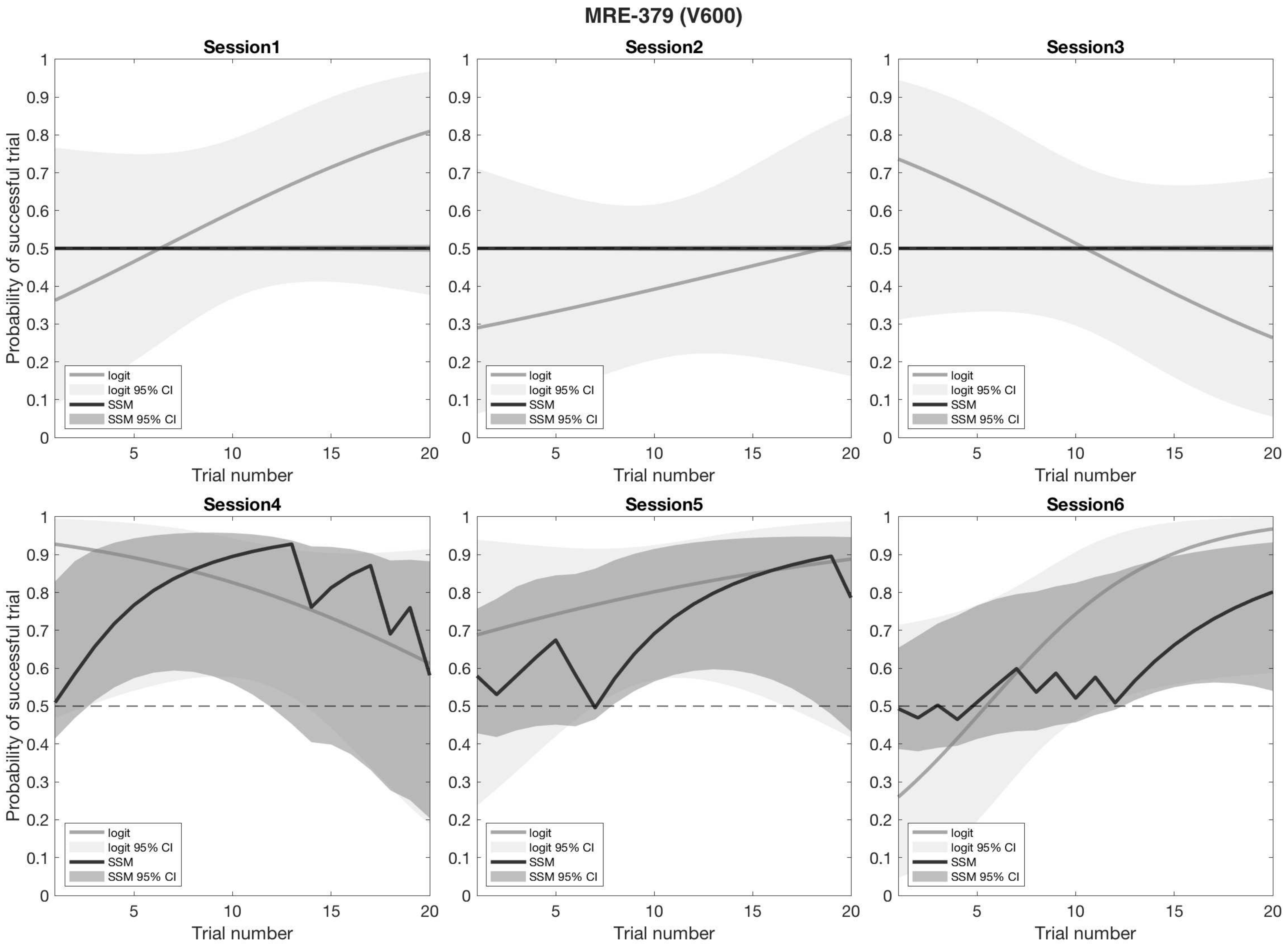

MRE-379 (V600)
Session1
Session2
Session3
Session4
Session5
Session6
Probability of successful trial
Trial number
logit
logit 95% CI
SSM
SSM 95% CI

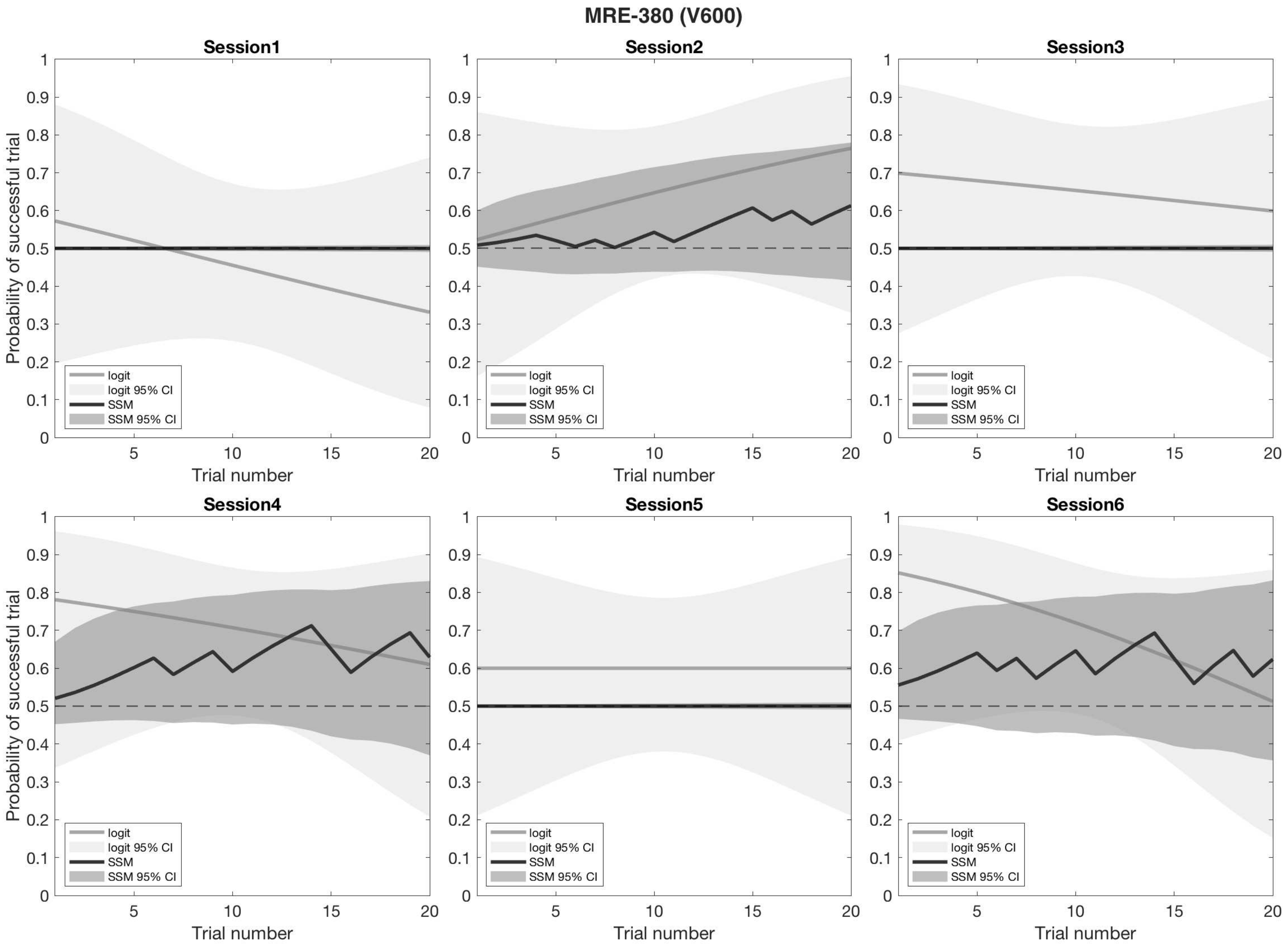
MRE-380 (V600)
Session1
Session2
Session3
Session4
Session5
Session6
Probability of successful trial
Trial number
logit
logit 95% CI
SSM
SSM 95% CI

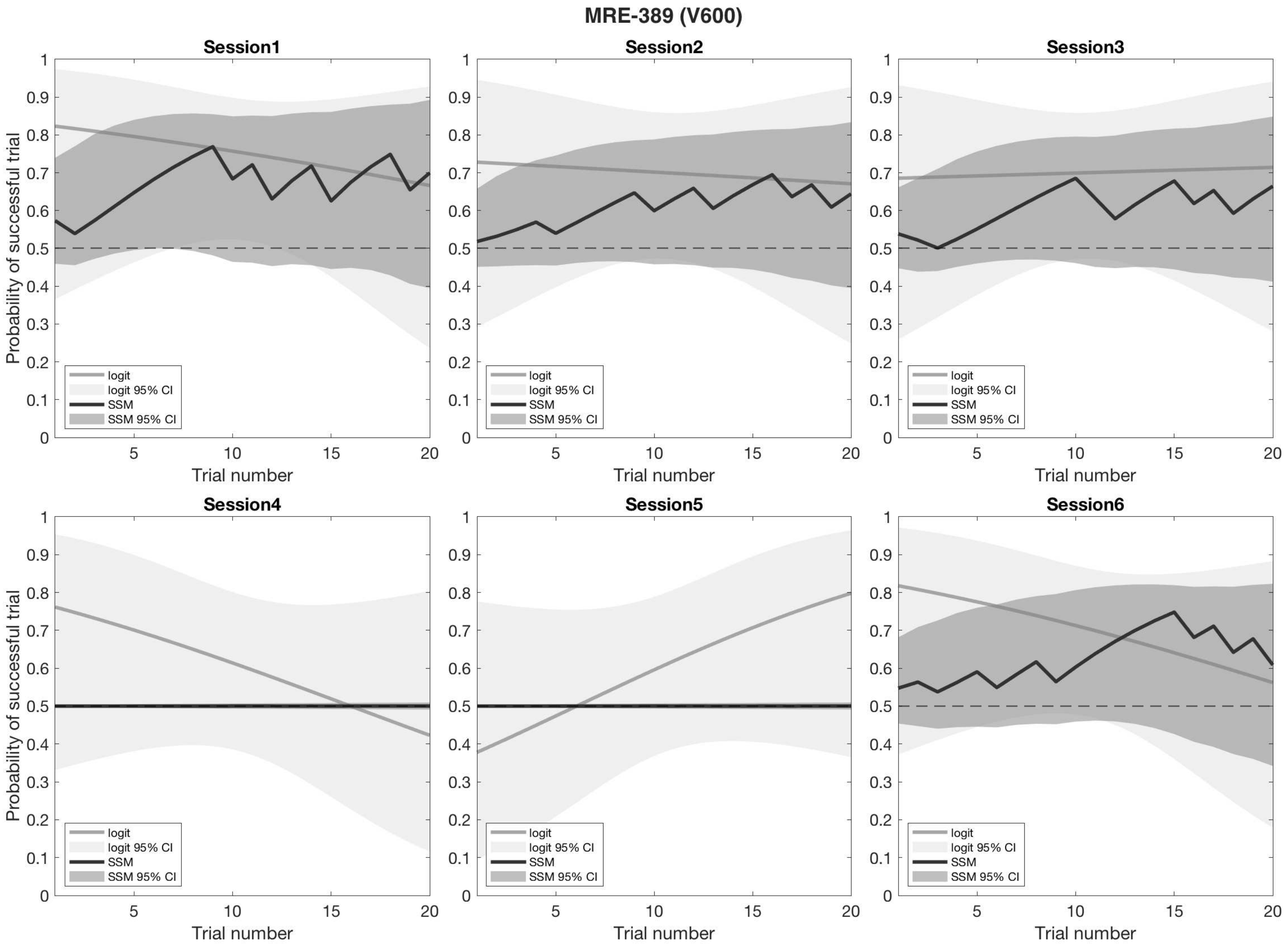
MRE-389 (V600)
Session1
Session2
Session3
Session4
Session5
Session6
Probability of successful trial
Trial number
logit
logit 95% CI
SSM
SSM 95% CI

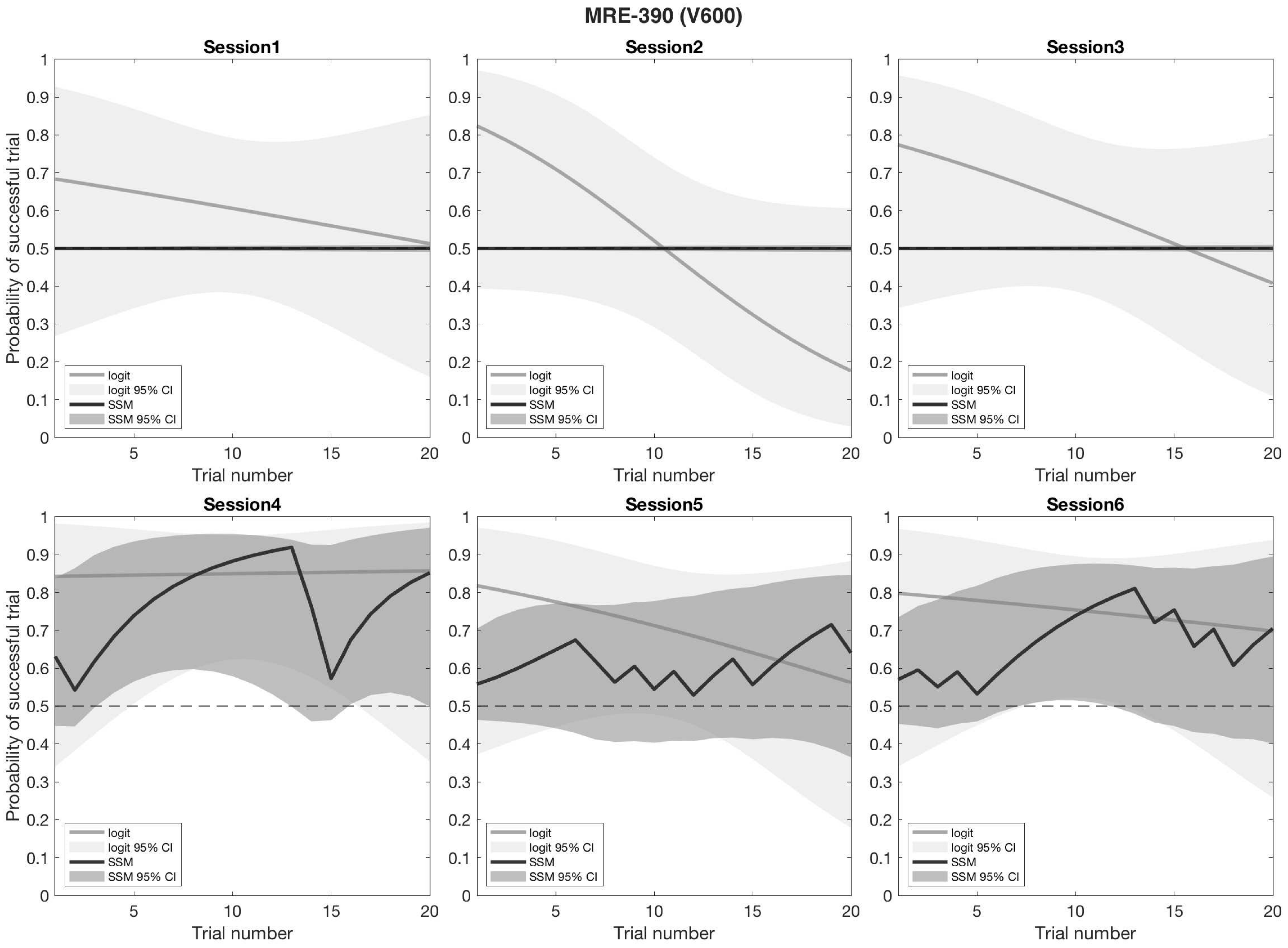

MRE-390 (V600)
Session1
Session2
Session3
Session4
Session5
Session6
Probability of successful trial
Trial number
logit
logit 95% CI
SSM
SSM 95% CI

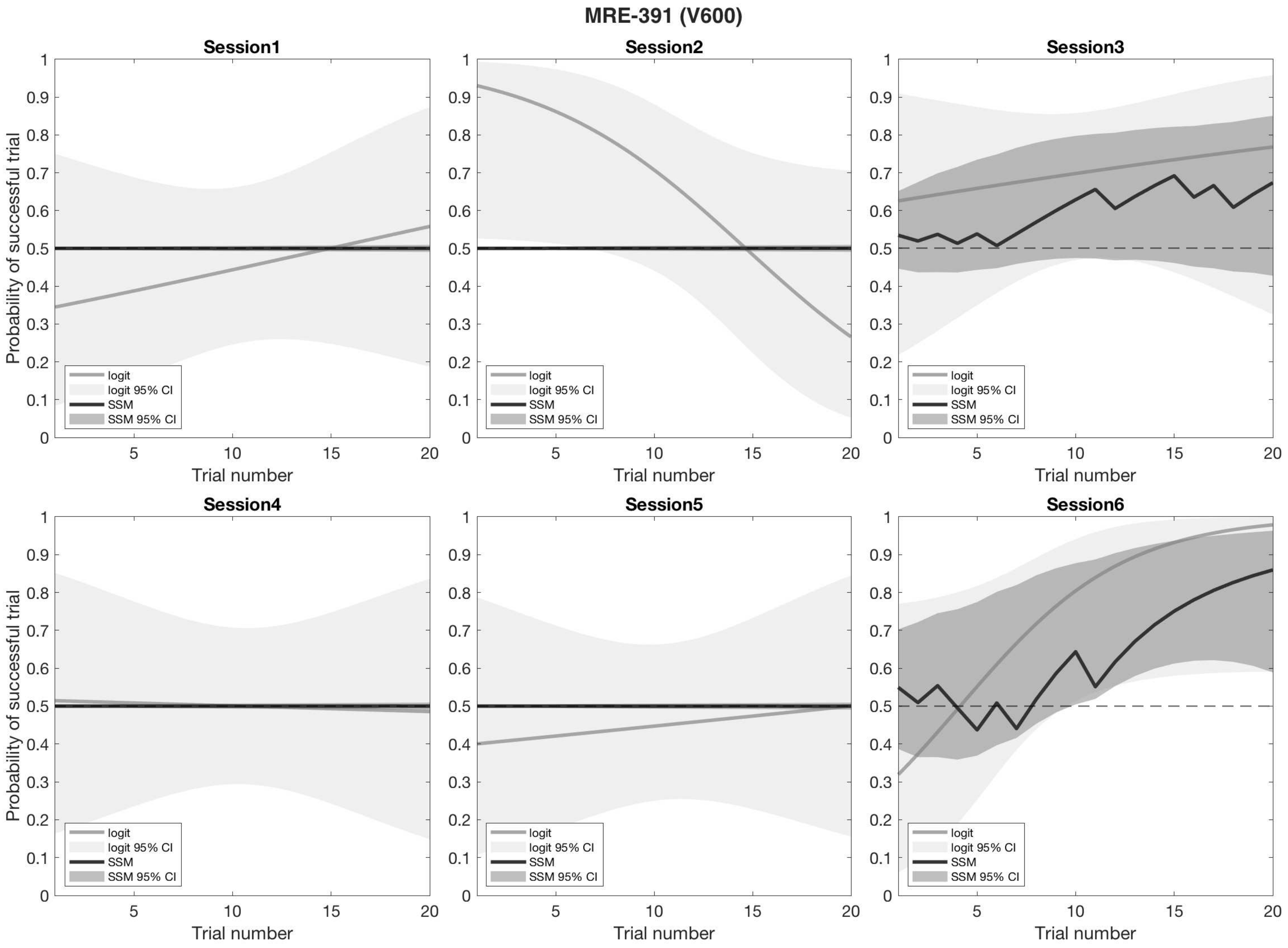

MRE-391 (V600)
Session1
Session2
Session3
Session4
Session5
Session6
Probability of successful trial
Trial number
logit
logit 95% CI
SSM
SSM 95% CI

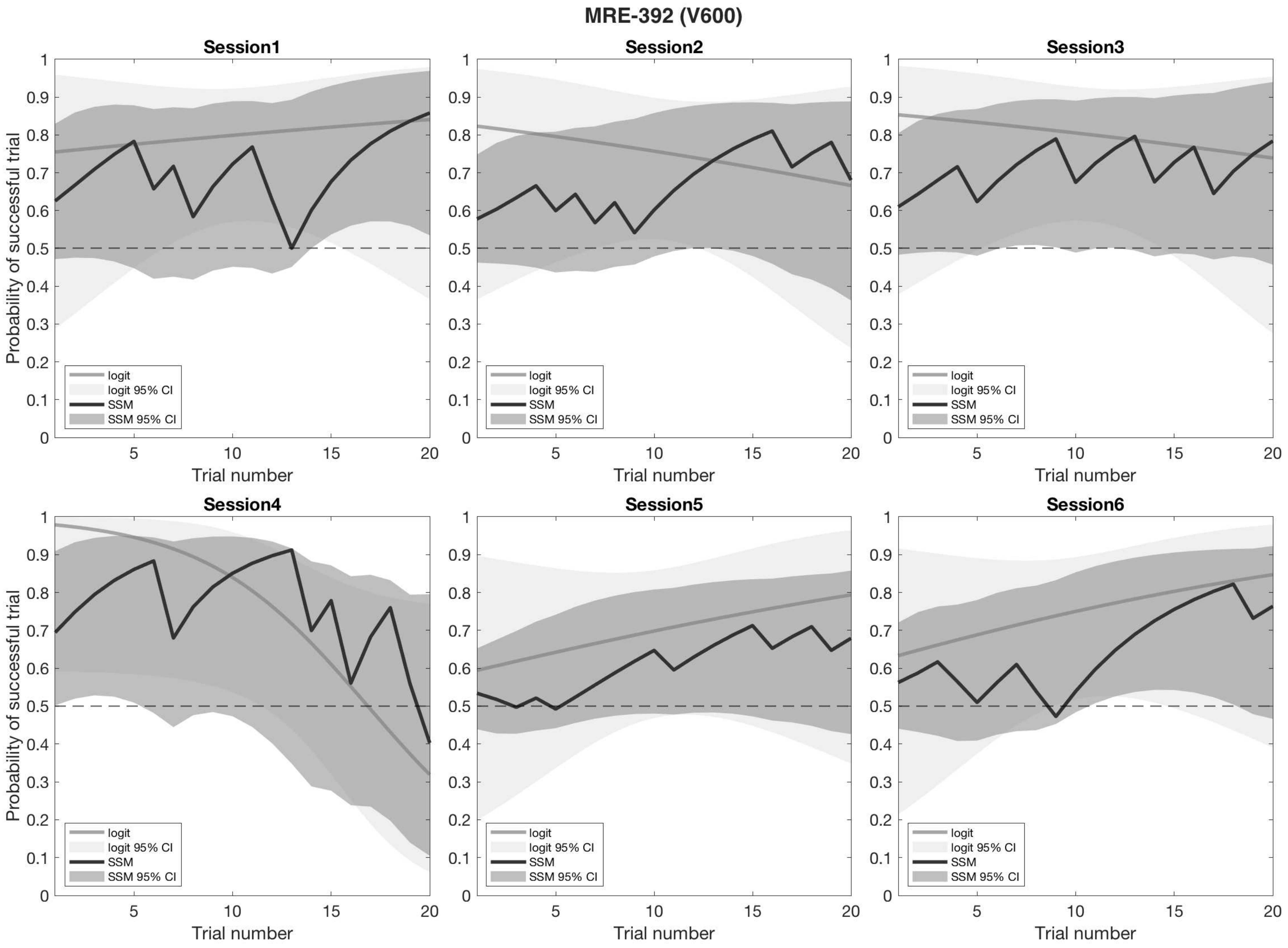
MRE-392 (V600)
Session1
Session2
Session3
Session4
Session5
Session6
Probability of successful trial
Trial number
logit
logit 95% CI
SSM
SSM 95% CI
0
0.1
0.2
0.3
0.4
0.5
0.6
0.7
0.8
0.9
1
5
10
15
20

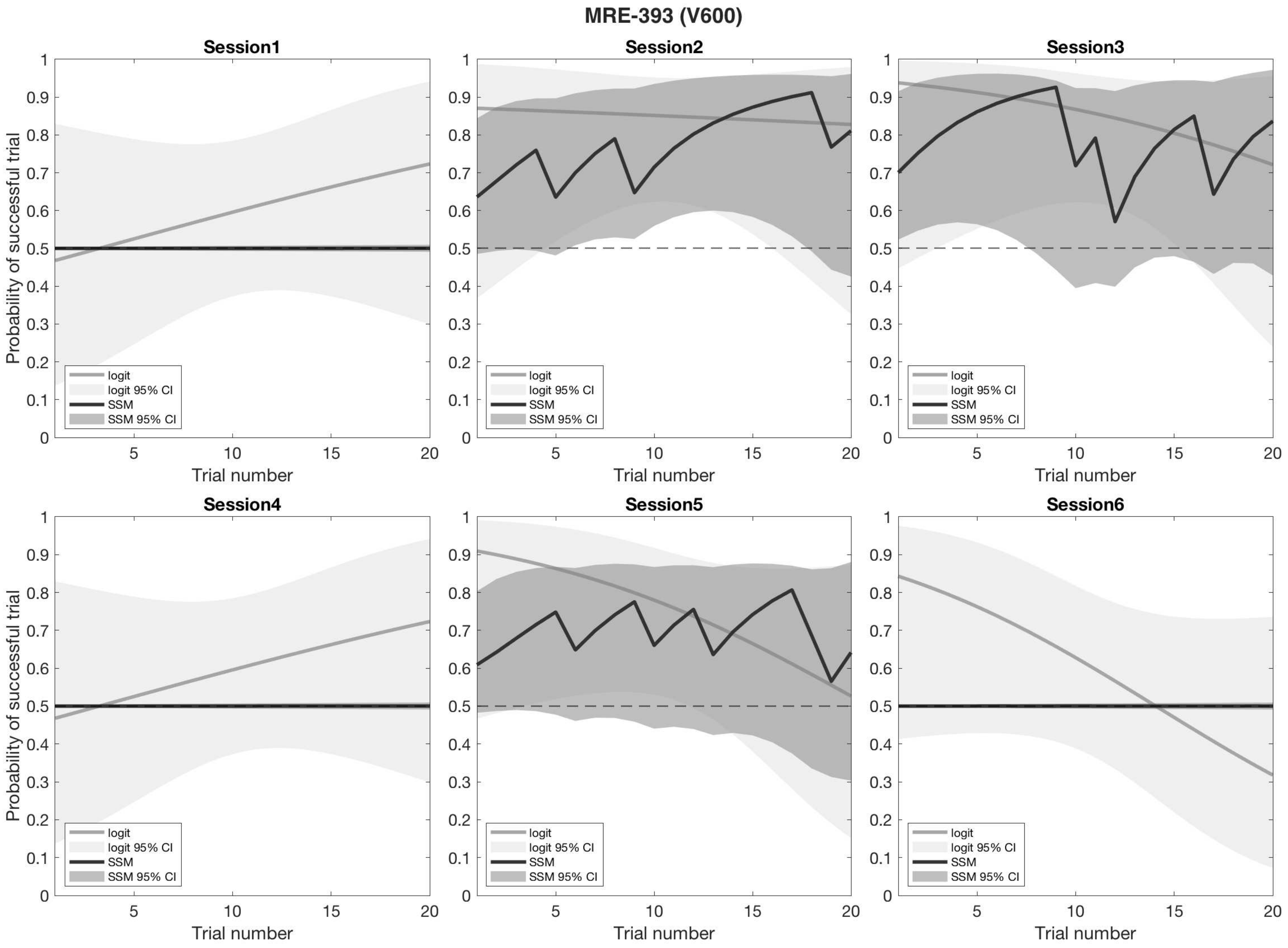

MRE-393 (V600)
Session1
Session2
Session3
Session4
Session5
Session6
Probability of successful trial
Trial number
logit
logit 95% CI
SSM
SSM 95% CI

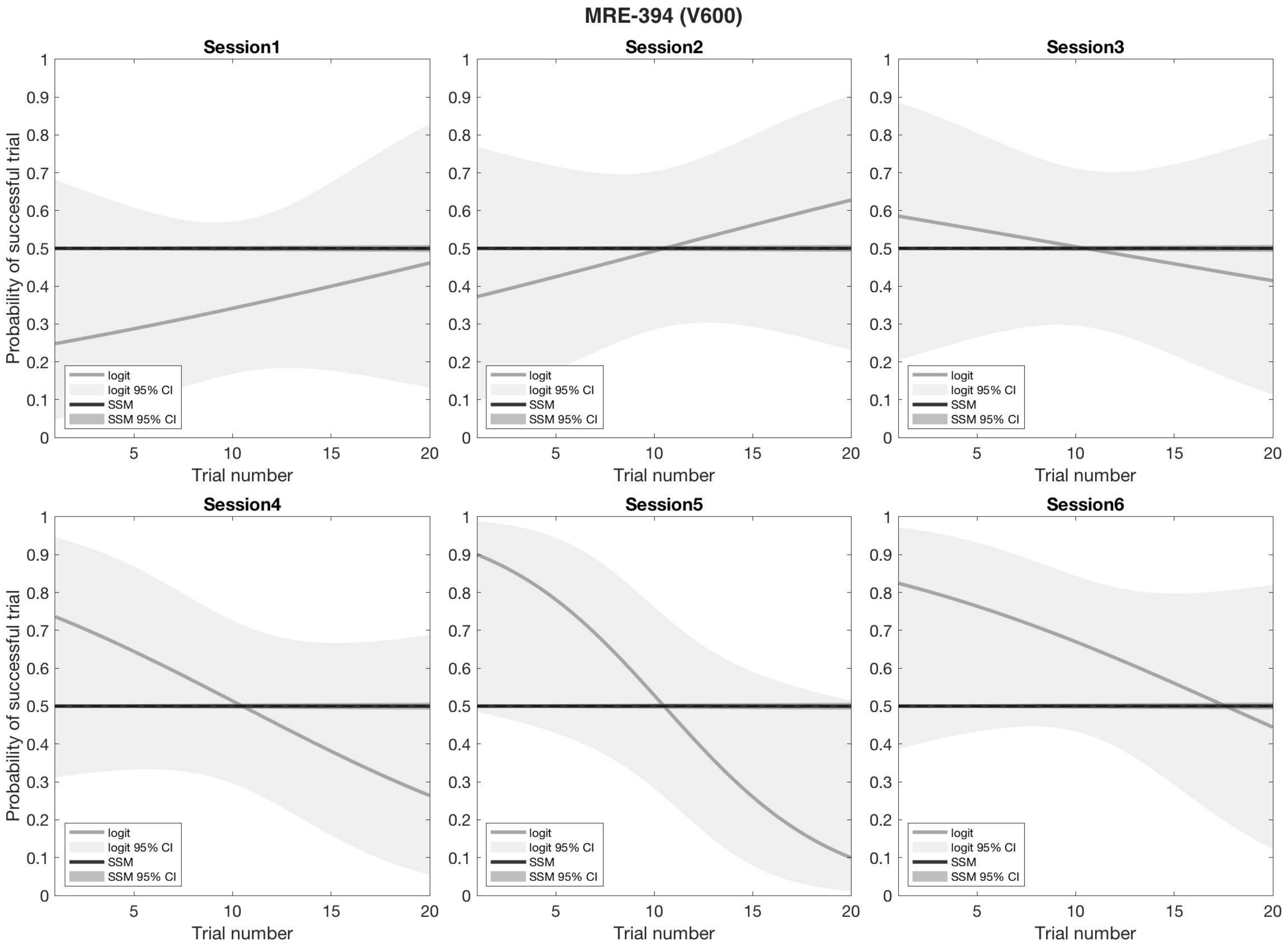
MRE-394 (V600)
Session1
Session2
Session3
Session4
Session5
Session6
Probability of successful trial
Trial number
logit
logit 95% CI
SSM
SSM 95% CI
0
0.1
0.2
0.3
0.4
0.5
0.6
0.7
0.8
0.9
1
5
10
15
20

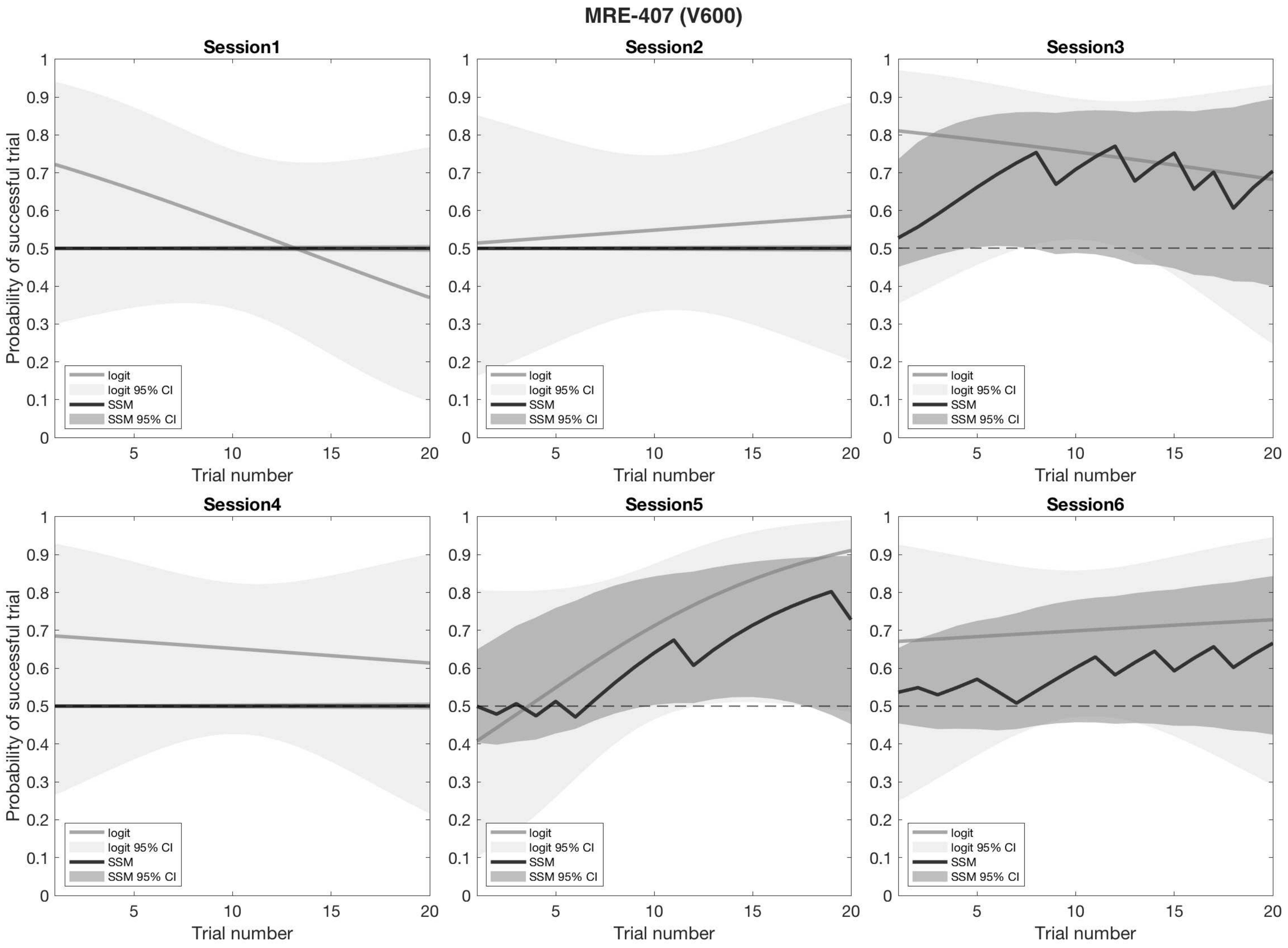
MRE-407 (V600)
Session1
Session2
Session3
Session4
Session5
Session6
Probability of successful trial
Trial number
logit
logit 95% CI
SSM
SSM 95% CI

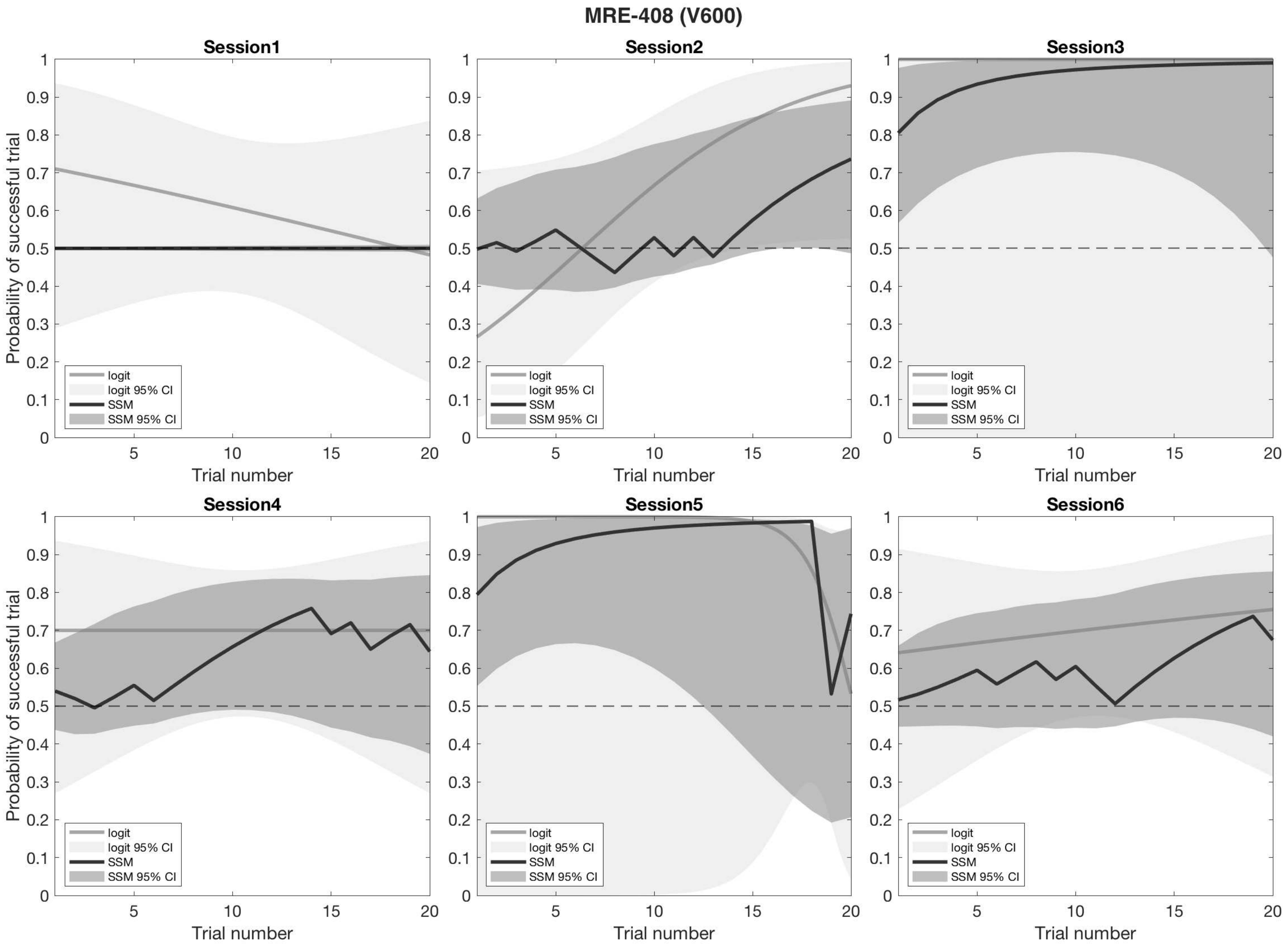
MRE-408 (V600)
Session1
Session2
Session3
Session4
Session5
Session6
Probability of successful trial
Trial number
logit
logit 95% CI
SSM
SSM 95% CI

### MRE-409 (V600)

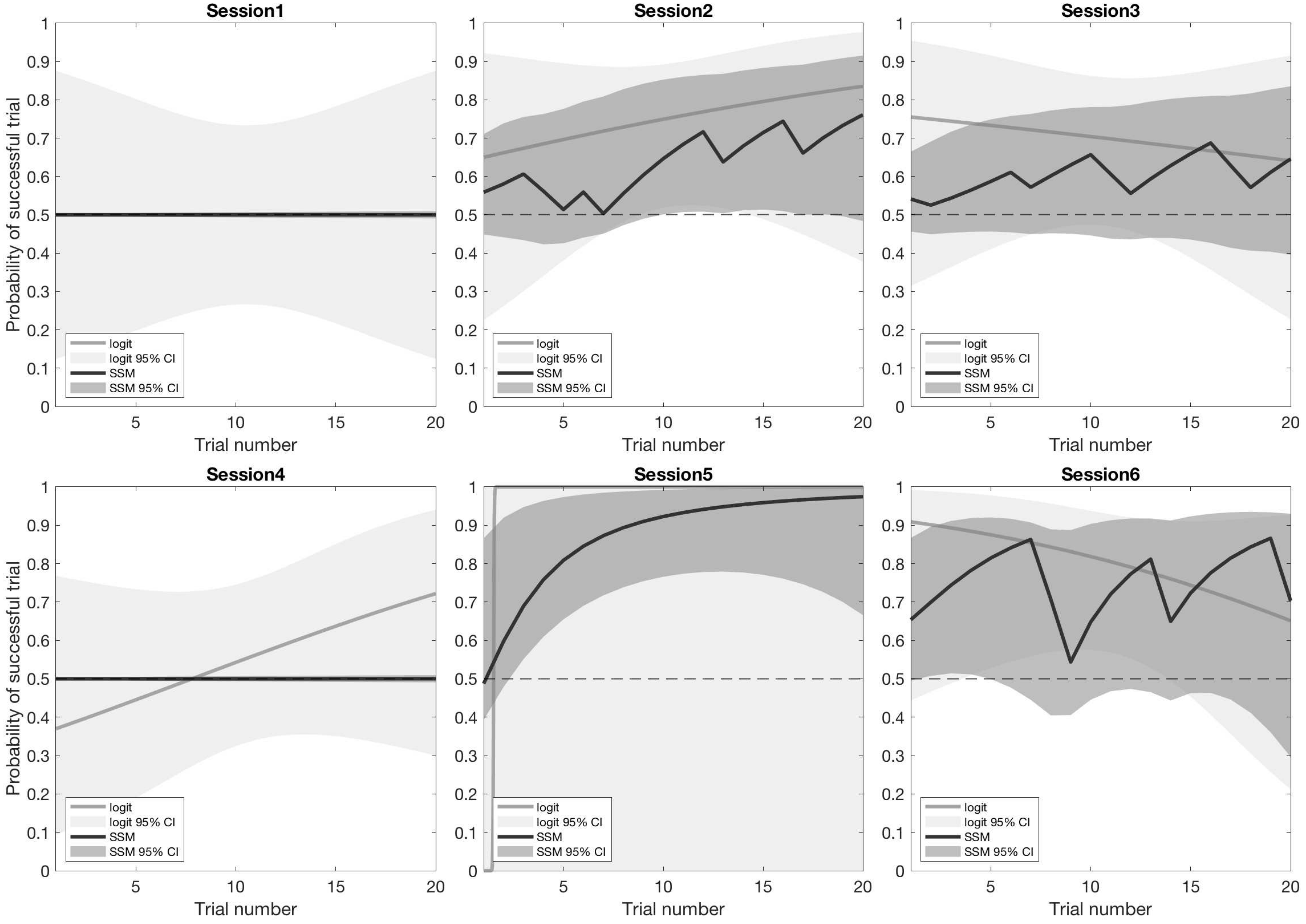

Session1
Session2
Session3
Session4
Session5
Session6
Probability of successful trial
Trial number
logit
logit 95% CI
SSM
SSM 95% CI

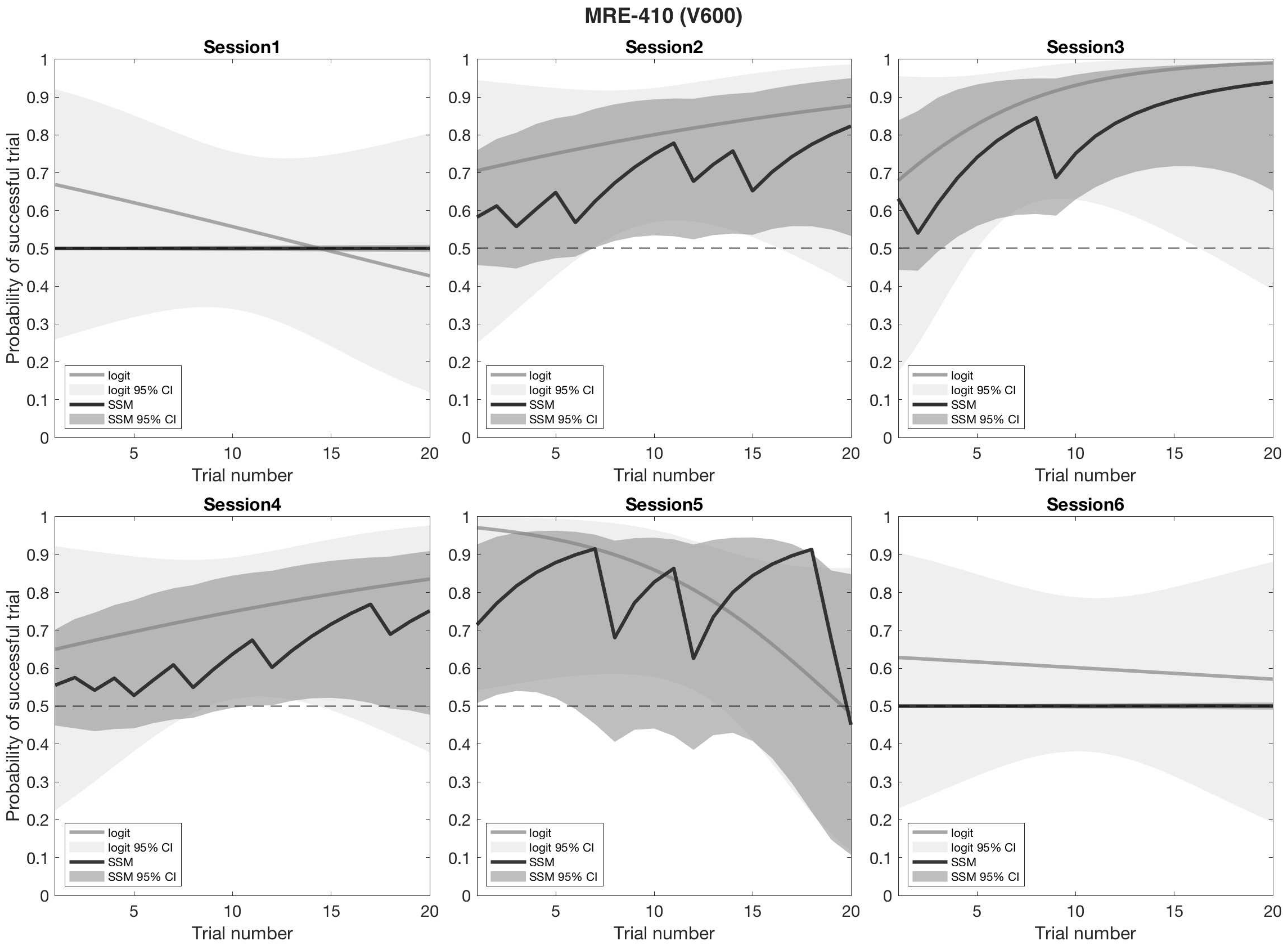
MRE-410 (V600)
Session1
Session2
Session3
Session4
Session5
Session6
Probability of successful trial
Trial number
logit
logit 95% CI
SSM
SSM 95% CI

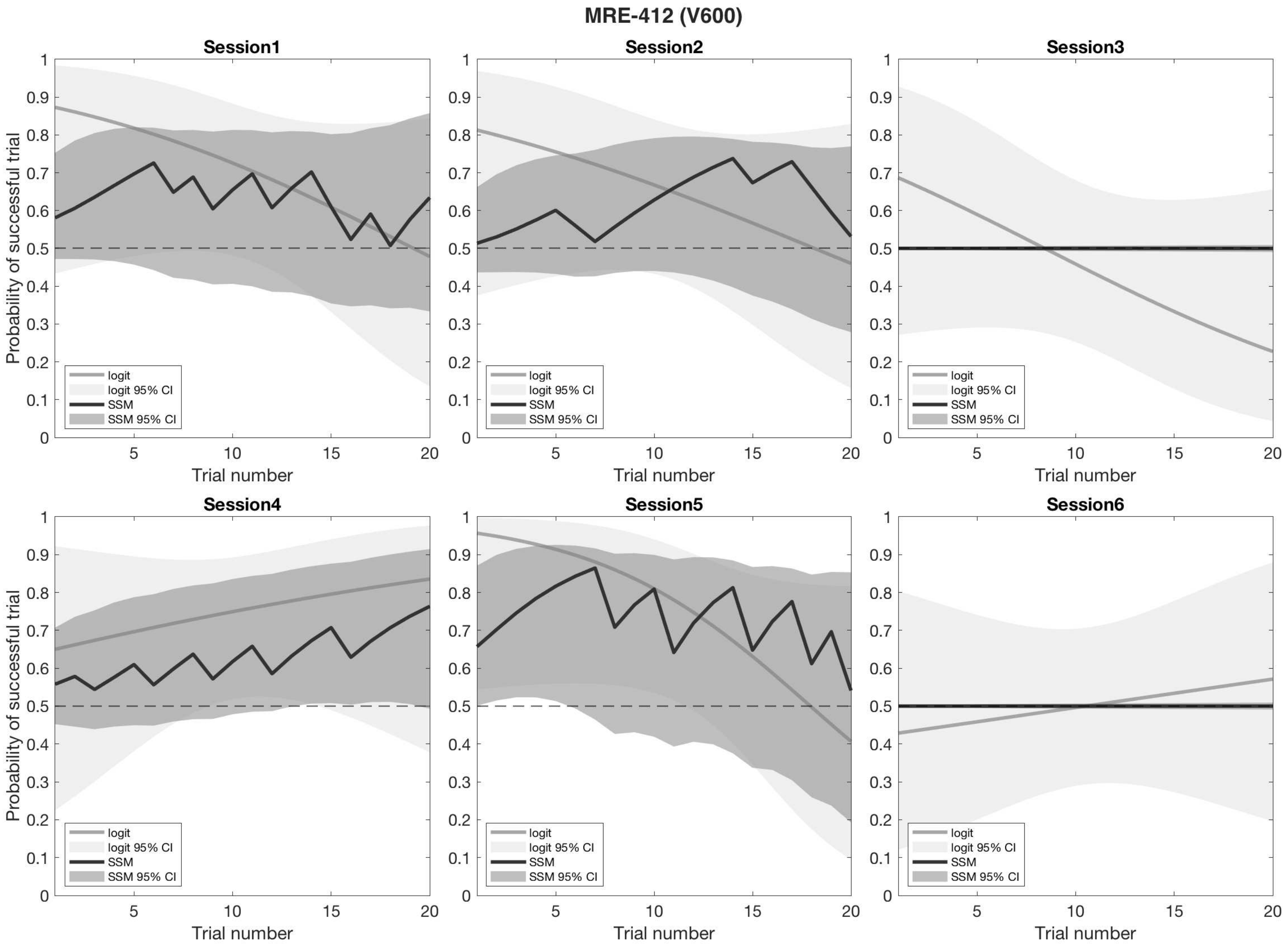

MRE-412 (V600)
Session1
Session2
Session3
Session4
Session5
Session6
Probability of successful trial
Trial number
logit
logit 95% CI
SSM
SSM 95% CI

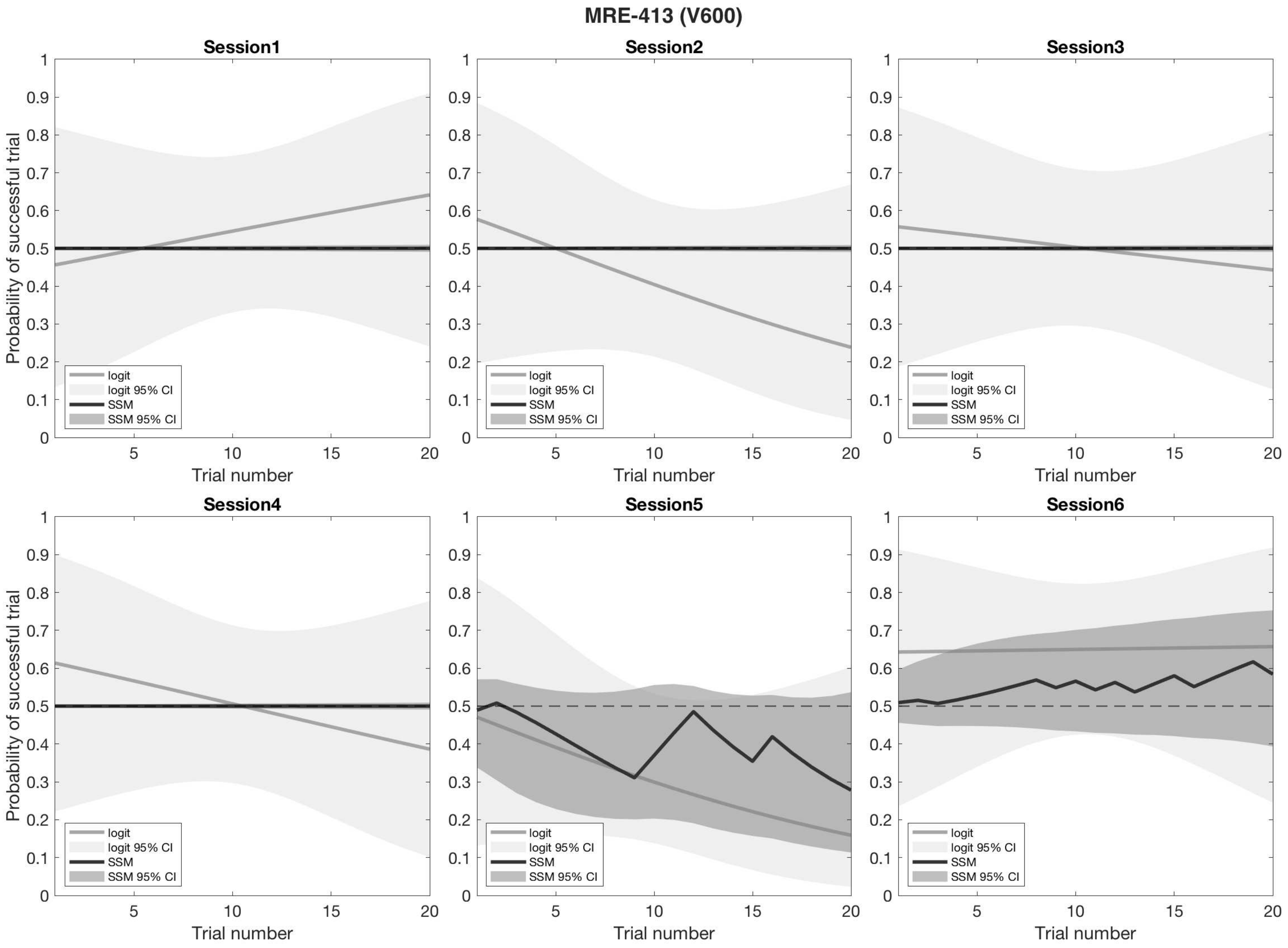
MRE-413 (V600)
Session1
Session2
Session3
Session4
Session5
Session6
Probability of successful trial
Trial number
logit
logit 95% CI
SSM
SSM 95% CI

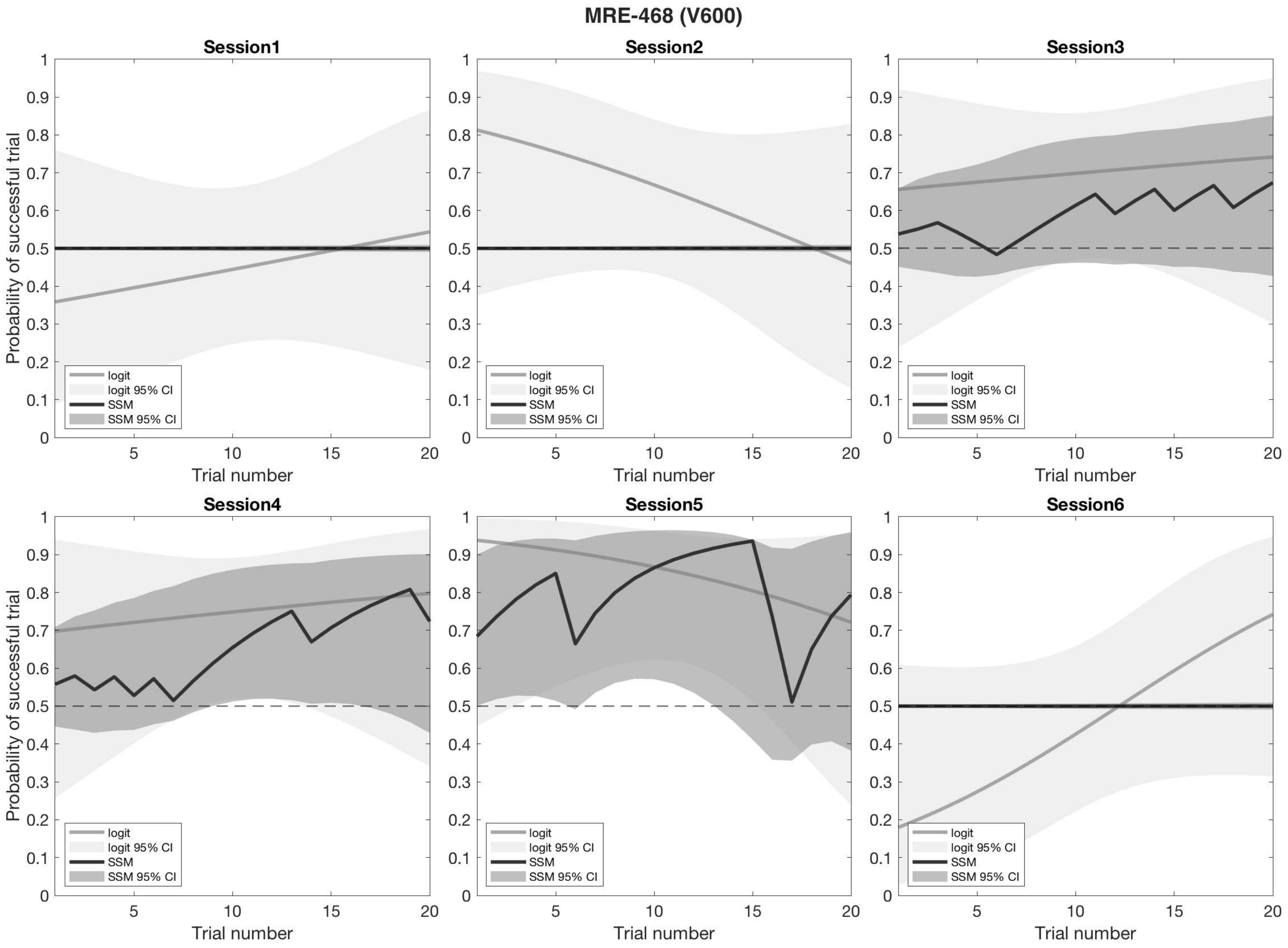

MRE-468 (V600)
Session1
Session2
Session3
Session4
Session5
Session6
Probability of successful trial
Trial number
0
0.1
0.2
0.3
0.4
0.5
0.6
0.7
0.8
0.9
1
5
10
15
20
logit
logit 95% CI
SSM
SSM 95% CI
